# Modelling the spatiotemporal spread of beneficial alleles using ancient genomes

**DOI:** 10.1101/2021.07.21.453231

**Authors:** Rasa Muktupavela, Martin Petr, Laure Ségurel, Thorfinn Korneliussen, John Novembre, Fernando Racimo

## Abstract

Ancient genome sequencing technologies now provide the opportunity to study natural selection in unprecedented detail. Rather than making inferences from indirect footprints left by selection in present-day genomes, we can directly observe whether a given allele was present or absent in a particular region of the world at almost any period of human history within the last 10,000 years. Methods for studying selection using ancient genomes often rely on partitioning individuals into discrete time periods or regions of the world. However, a complete understanding of natural selection requires more nuanced statistical methods which can explicitly model allele frequency changes in a continuum across space and time. Here we introduce a method for inferring the spread of a beneficial allele across a landscape using two-dimensional partial differential equations. Unlike previous approaches, our framework can handle time-stamped ancient samples, as well as genotype likelihoods and pseudohaploid sequences from low-coverage genomes. We apply the method to a panel of published ancient West Eurasian genomes to produce dynamic maps showcasing the inferred spread of candidate beneficial alleles over time and space. We also provide estimates for the strength of selection and diffusion rate for each of these alleles. Finally, we highlight possible avenues of improvement for accurately tracing the spread of beneficial alleles in more complex scenarios.

## Introduction

Understanding the dynamics of the spread of a beneficial allele through a population is one of the fundamental problems in population genetics ***(Ewens,* 2012).** We are often interested in knowing the location where an allele first arose and the way in which it spread through a population, but this is often unknown, particularly in natural, non-experimental settings where genetic sampling is scarce and uneven.

Patterns of genetic variation can be used to estimate how strongly natural selection has affected the trajectory of an allele and to fit the parameters of the selection process. The problem of estimating the age of a beneficial allele, for example, has yielded a rich methodological literature ***(Slatkin and Ranna/a,* 2000),** and recent methods have exploited fine-scale haplotype information to produce highly accurate age estimates ***(Mathieson and Mcvean, 2014; Platt et al., 2019; Albers and Mcvean,* 2020).** In contrast, efforts to infer the geographic origins of beneficial mutations are scarcer. These include ***Novembre et al.* (2005),** who developed a maximum likelihood method to model the origin and spread of a beneficial mutation and applied it to the CCRS-l:132 allele, which was, at the time, considered to have been under positive selection ***(Stephens et al., 1998; Sabeti et al., 2005; Novembre and Han, 2012).*** Similarly, ***/tan et al.* (2009)** developed an approximate Bayesian computation (ABC) approach using demic simulations, in order to find the geographic and temporal origins of a beneficial allele, based on present-day allele frequency patterns.

As ancient genome sequences become more readily available, they are increasingly being used to understand the process of natural selection (see reviews in ***Malaspinas et al. (2012); Dehasque et al.* (2020)).** However, few studies have used ancient genomes to fit spatial dynamic models of the spread of an allele over a landscape. Most spatiotemporal analyses which included ancient genomes have used descriptive modelling in order to learn the spatiotemporal covariance structure of allele frequencies ***(Segurel et al.,* 2020)** or hidden ancestry clusters ***(Racimo et al., 2020b),*** and then used that structure to hindcast these patterns onto a continuous temporally-evolving landscape. In contrast to descriptive approaches, dynamic models have the power to infer interpretable parameters from genomic data and perhaps reveal the ultimate causes for these patterns ***(Wikle et al., 2019)*.**

Dynamic models can also contribute to ongoing debates about the past trajectories of phenotypically important loci. For example, the geographic origin of the rs4988235(T) allele-upstream of the *LCT* gene and associated with adult lactase persistence in most of Western Eurasia ***(Enattah et al.,*** 2002)-remains elusive, as is the way in which it spread (an extensive review can be found in ***Segurel and Bon, 2017).*** The allele has been found in different populations, with frequencies ranging from 5% up to almost 100%, and its selection coefficient has been estimated to be among the highest in human populations ***(Bersag/ieri et al., 2004; Enattah et al., 2008; Tishkoff et al., 2007).*** However, the exact causes for its adaptive advantage are contested ***(Szpak et al., 2019),*** and it has been suggested that the selection pressures acting on the allele may have been different in different parts of the continent ***(Gerbault et al.,* 2009).** Ancient DNA evidence shows that the allele was rare in Europe during the Neolithic ***(Burger et al., 2007; Gamba et al., 2014; Allentoft et al., 2015; Mathieson et al., 2015)*** and only became common in Northern Europe after the Iron Age, suggesting a rise in frequency during this period, perhaps mediated by gene flow from regions east of the Baltic where this allele was more common during the onset of the Bronze Age ***(Kriittli et al., 2014; Margaryan et al.,* 2020). */tan et al.* (2009)** deployed their ABC approach to model the spatial spread of the rs4988235(T) allele and estimated that it was first under selection among farmers around 7,500 years ago possibly between the central Balkans and central Europe. Others have postulated a steppe origin for the allele ***(Allentoft eta/.,2015),*** given that the rise in frequency appears to have occurred during and after the Bronze Age migration of steppe peoples into Western Eurasia ***(Haak et al., 2015; Allentoft et al., 2015).*** However, the allele is at low frequency in genomes of Bronze Age individuals associated with Corded Ware and Bell Beaker assemblages in Central Europe who have high steppe ancestry ***(Mathieson et al., 2015; Margaryan et al.,* 2020),** complicating the story further ***(Segurel and Bon, 2017)*.**

The origins and spread dynamics of large-effect pigmentation-associated SNPs in ancient Eurasians have also been intensely studied ***Uu and Mathieson,* 2020).** Major loci of large effect on skin, eye and hair pigmentation have been documented as having been under recent positive selection in Western Eurasian history ***(Voight et al., 2006; Sabeti et al., 2007; Pickrell et al., 2009; Lao et al., 2007; Mathieson et al., 2015; Alonso et al., 2008; Hudjashov et al., 2013).*** These include genes *SLC45A2, OCA2, HERC2, SLC24A5* and *TYR.* While there is extensive evidence supporting the adaptive significance of these alleles, debates around their exact origins and spread are largely driven by comparisons of allele frequency estimates in population groups which are almost always discretized in time and/or space. Among these, selection at the *TYR* locus is thought to have occurred particularly recently, over the last 5,000 years ***(Stern et al., 2019),*** driven by a recent mutation ***(Albers and Mcvean,* 2020)** that may have spread rapidly in Western Eurasia.

Here, we develop a method to model the spread of a recently selected allele across both space and time, avoiding artificial discretization schemes to more rigorously assess the evidence for or against a particular dispersal process. We begin with the model proposed by ***Novembre et al. (2005),*** and adapt it in order to handle ancient low-coverage genomic data, and explore more complex models that allow for both diffusion and advection (i.e. directional transport) in the distribution of allele frequencies over space, as well asfor a change in these parameters at different periods of time. We apply the method to alleles in two of the aforementioned loci in the human genome, which have been reported to have strong evidence for recent positive selection: *LCT/MCM6* and *TYR.* We focus on Western Eurasia during the Holocene, where ancient genomes are most densely sampled, and infer parameters relevant to the spread of these alleles, including selection, diffusion and advection coefficients.

## Results

### Summary of model

We based our statistical inference framework on a model proposed by ***Novembre et al. (2005)*** to fit allele frequencies in two dimensions to present-day genotype data spread over a densely sampled map. We extend this model in several ways:

- We incorporate temporally sampled data (ancient genomes) to better resolve changes in frequency distributions over time
- We make use of genotype likelihoods and pseudohaploid genotypes to incorporate low-coverage data into the inference framework
- We permit more general dynamics by including advection parameters.
- We allow the selection, advection and diffusion parameters to be different in different periods of time. Specifically, to reflect changes in population dynamics and mobility before and after the Bronze Age ***(Loog et al., 2017; Racimo et al., 2020a),*** we partitioned the model fit into two time periods: before and after 5,000 years BP.

We explored the performance of two different spread models, which are extensions of the original model by ***Novembre et al. (2005),*** hereby called model A This is a diffusion model containing a selection coefficients (determining the rate of local allele frequency growth) and a single diffusion term (σ). A more general diffusion model - hereby model B - allows for two distinct diffusion parameters for latitudinal (σ_y_) and longitudinal (σ_x_) spread. Finally, model C is even more general and includes two advection terms *(v_x_* and *v_y_),* allowing the center of mass of the allele’s frequency to diverge from its origin over time. The incorporation of advection is meant to account for the fact that population displacements and expansions could have led to allele frequency dynamics that are poorly explained by diffusion alone.

In order to establish a starting time point for our diffusion process, we used previously published allele age estimates obtained from a non-parametric approach leveraging the patterns of haplotype concordance and discordance around the mutation of interest ***(Albers and Mcvean,* 2020).** In the case of the allele in the *LCTIMCM6* region, we also used age estimates based on an approximate Bayesian computation approach ***(/tan et al.,* 2009).**

### Performance on deterministic simulations

To characterize the accuracy of our inference method under different parameter choices we first generated deterministic simulations from several types of diffusion models. First, we produced an allele frequency surface map with a specified set of parameters from which we drew 1,040 samples matching the ages, locations and genotype calling format (diploid vs. pseudo-haploid) of the 1,040 genomes that we analyze below when studying the rs1042602(A) allele.

We generated six different simulations with different diffusion coefficients and afterwards ran our method assuming model B. The results (simulations B1-B6) are summarised in ***Figure 1, Figure 1-Figure Supplement 1, Figure 1-Figure Supplement 1, Figure 1-Figure Supplement 1, Figure 1-Figure Supplement 1, Figure 1-Figure Supplement*** Sand ***Table A1.*** Overall, the model is more accurate at correctly inferring the parameters for the time period before 5,000 years BP ***(Figure 1b)*,** with decreased performance when longitudinal diffusion is high ***(Figure 1-Figure Supplement 5*).**

**Figure 1.**
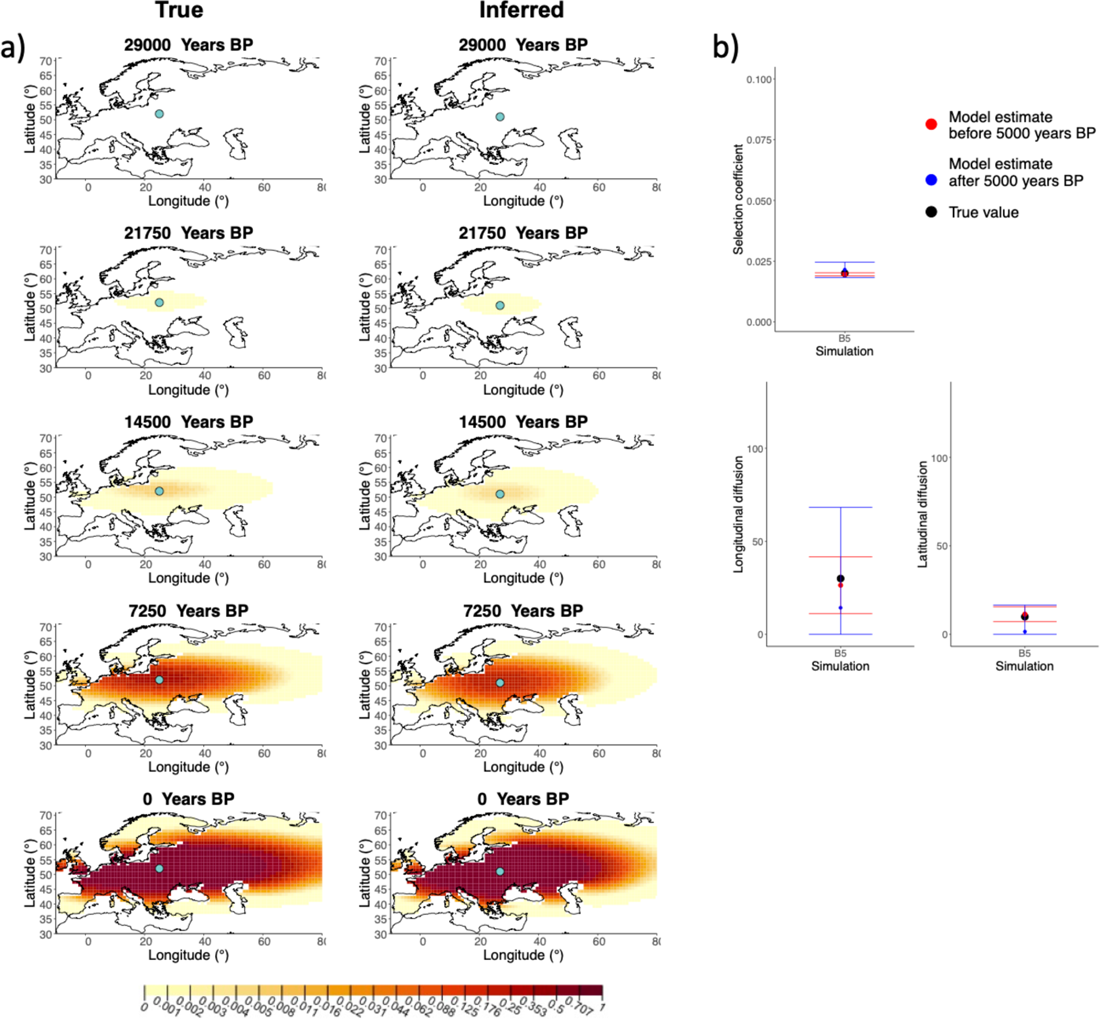
a) Comparison of true andinferred allele frequency dynamics for a simulation with diffusion and no advection (BS). The green dot corresponds to the origin of the allele. The parameter values used to generate the frequency surface maps are summarised in ***Table A1.*** b) Comparison of true parameter values and model estimates. Whiskers represent 95% confidence intervals. **Figure 1-Figure supplement 1**. Comparison of true and inferred allele frequency dynamics for simulation 81. Figure 1-Figure supplement 2. Comparison of true and inferred allele frequency dynamics for simulation 82. Figure 1-Figure supplement 3. Comparison of true and inferred allele frequency dynamics for simulation 83. Figure 1-Figure supplement 4. Comparison of true and inferred allele frequency dynamics for simulation 84. Figure 1-Figure supplement 5. Comparison of true and inferred allele frequency dynamics for simulation 86. Figure 1-Figure supplement 6. Comparison of true allele frequency dynamics for simulation 81 and those inferred by the model C. Figure 1-Figure supplement 7. Comparison of true allele frequency dynamics for simulation 84 and those inferred by the model C.

Next, we investigated the performance of model C, which includes advection coefficients. We generated four different simulations including advection (simulations C1-C4: ***Figure 2, Figure 2-Figure Supplement 2, Figure 2-Figure Supplement 2, Figure 2-Figure Supplement 2*** and ***Table* A2).** We found that our method is generally able to estimate the selection coefficient accurately. However, in some of the simulations, we found discrepancies between the estimated and true diffusion and advection coefficients, often occurring because of a misestimated origin forcing the other parameters to adjust in order to better fit the allele frequency distribution in later stages of the allele’s spread ***(Figure* 2).** Despite the disparities between the true and inferred parameter values, the resulting surface plots become very similar as we approach the present, suggesting that different combinations of parameters can produce similar present-day allele frequency distributions.

**Figure 2.**
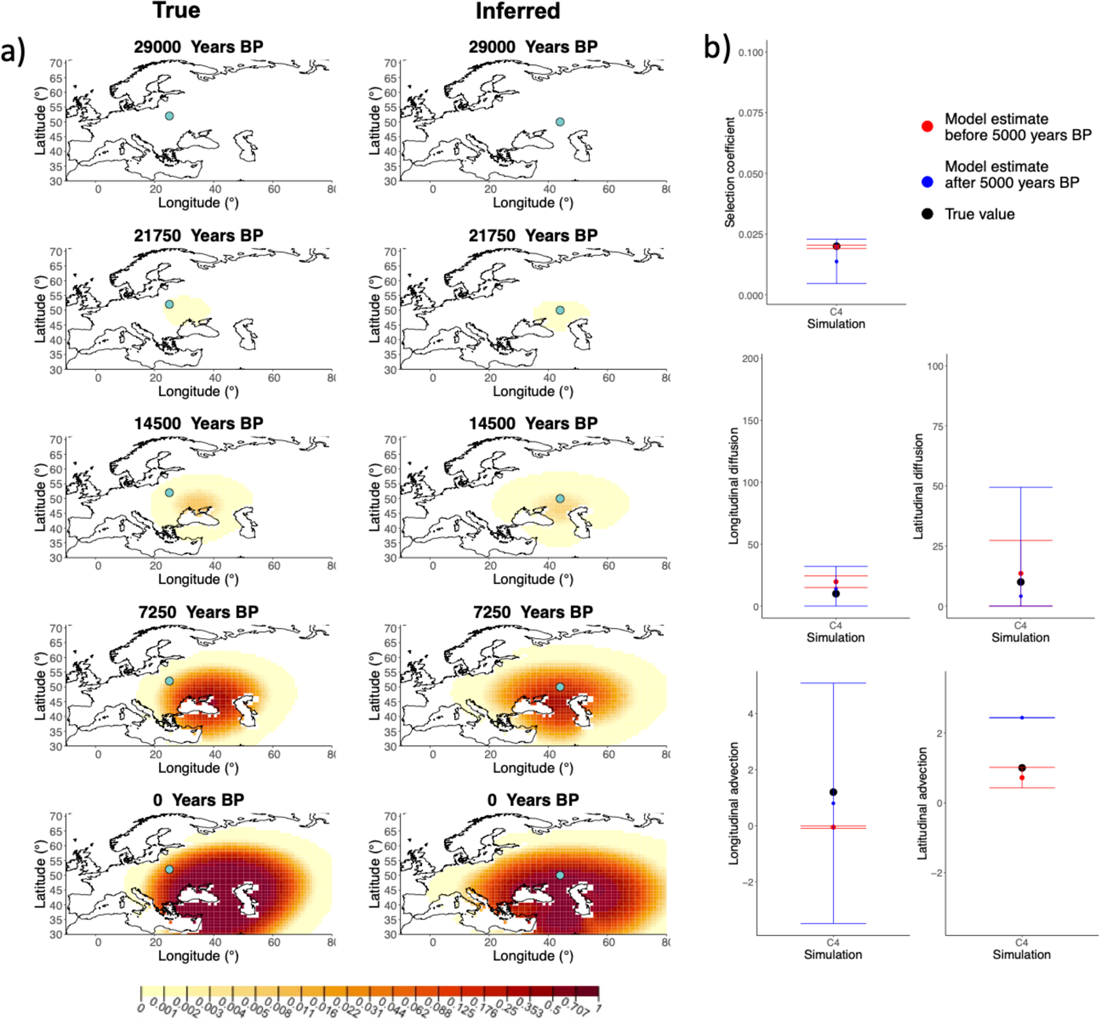
a) Comparison of true and inferred allele frequency dynamics for one of the simulations including advection (C4). The green dot corresponds to the origin of the allele. The parameter values used to generate the frequency surface maps are summarised in ***Table A2.*** b) Comparison of true parameter values and model estimates. Whiskers represent 95% confidence intervals. **Figure 2-Figure supplement 1.** Comparison of true and inferred allele frequency dynamics for simulation C1. **Figure 2-Figure supplement 2.** Comparison of true and inferred allele frequency dynamics for simulation C2. **Figure 2-Figure supplement 3.** Comparison of true and inferred allele frequency dynamics for simulation C3.

### Advection model application to non-advection simulations

We assessed the model performance when we apply the model C, which includes advection coefficient estimates, to simulations generated without advection (see ***Figure 1-Figure Supplement 1*** and ***Figure 1-Figure Supplement 7*).** We can observe that the advection coefficients are inferred to be non-zero ***(Figure 1-Figure Supplement 6b*** and ***Figure 1-Figure Supplement 7b*),** however the inferred allele frequency dynamic plots closely resemble the ones obtained with with true parameter values ***(Figure 1-Figure Supplement 1a*** and ***Figure 1-Figure Supplement 7a*).** This shows that complex interactions between the diffusion and advection coefficients can result in similar outcomes even when only diffusion is considered in the model.

The inference of the origin of the allele also differs when we compare the results for using model 8 and model C. In order to understand better how the model estimates the allele origin, we highlighted the first individual in simulations 81 and 84 that contains the derived allele. We can see that in case of simulation 81 the inferred origin of the allele is close to the first observance of the derived allele in the model which includes advection. In contrast when the advection is not included, the origin of the allele is inferred to be closer to where it is initially rising in frequency ***(Figure 1-Figure Supplement 1a*** and ***Figure 1-Figure Supplement 4a).*** However, this is not always the case. For instance, ifwe look at the results from the advection model on simulation B4, we can see that the origin of the allele is inferred relatively far from the sample known to have carried the first instance of the derived allele. Therefore, if there is a relatively large interval between the time when the allele originated and when the first ancient genomes are available, the beneficial allele can spread widely, but as this spread is not captured by any of the data points, inference of the precise origin of the selected allele is nearly impossible.

### Impact of sample clustering on parameter estimates

We evaluated the impact of different sampling and clustering schemes on our inferences that could potentially arise by aggregating aDNA data from studies with different sampling schemes. We used a deterministic simulation to create three different degrees of clustering which we will refer to as “homogeneous”, “intermediate” or “extreme” by varying the area from which we sample individuals to be used in our inferences ***(Figure3-FigureSupplement 1).*** Additionally, we also tested the impact of biased temporal sampling in the periods before and after 5000 year BP by oversampling in the ancient period (75%/25%), equal sampling in the two periods (50%/50%), and oversampling in the recent period (25%175%). Because we evaluated this temporal bias for each of the three spatial clustering sampling scenarios, this resulted in a total of 9 different sampling scenarios. We note that the third “extreme” spatial clustering scenarios is completely unrealistic and one would not expect inferences of any degree of accuracy from it, but we believe it gives a good idea of the behaviour of our method in the limit case of extremely restricted spatial sampling.

A comparison of allele frequency maps generated using true parameter values and using parameter estimates from the different sampling schemes are shown in ***Figure 3-Figure Supplement 2,Figure 3-Figure Supplement 3, Figure 3-Figure Supplement 3, Figure 3-Figure Supplement 5, Figure 3-Figure Supplement 3, Figure 3-Figure Supplement 7, Figure 3-Figure Supplement 8, Figure 3-Figure Supplement 3.* In *Figure 3*** we show the allele frequency map generated using the “intermediate 75%/25%” clustering scheme. Parameter estimates used to generate all these figures are summarised in ***Table A3.*** Overall we can see that the allele frequency maps inferred from these scenarios closely resemble the maps generated using the true parameter values, despite the challenges in finding accurate values for the individual point estimates of some of the parameters, highlighting that various combinations of diffusion and advection coefficients can produce similar underlying frequency maps (as discussed in the manuscript section “Performance on deterministic simulations”). This suggests that the joint spatiotemporal information encoded in the inferred maps (not just the individual parameters estimates) should be used in interpreting model outputs, particularly when it comes to the advection and diffusion parameters. The selection coefficient estimates are inferred highly accurately, regardless of the sampling scheme chosen, and lie close to the true value, with only a slight underestimation in the time period after 5000 years BP (with the exception of “extreme 25%/75%”).

**Figure 3.**
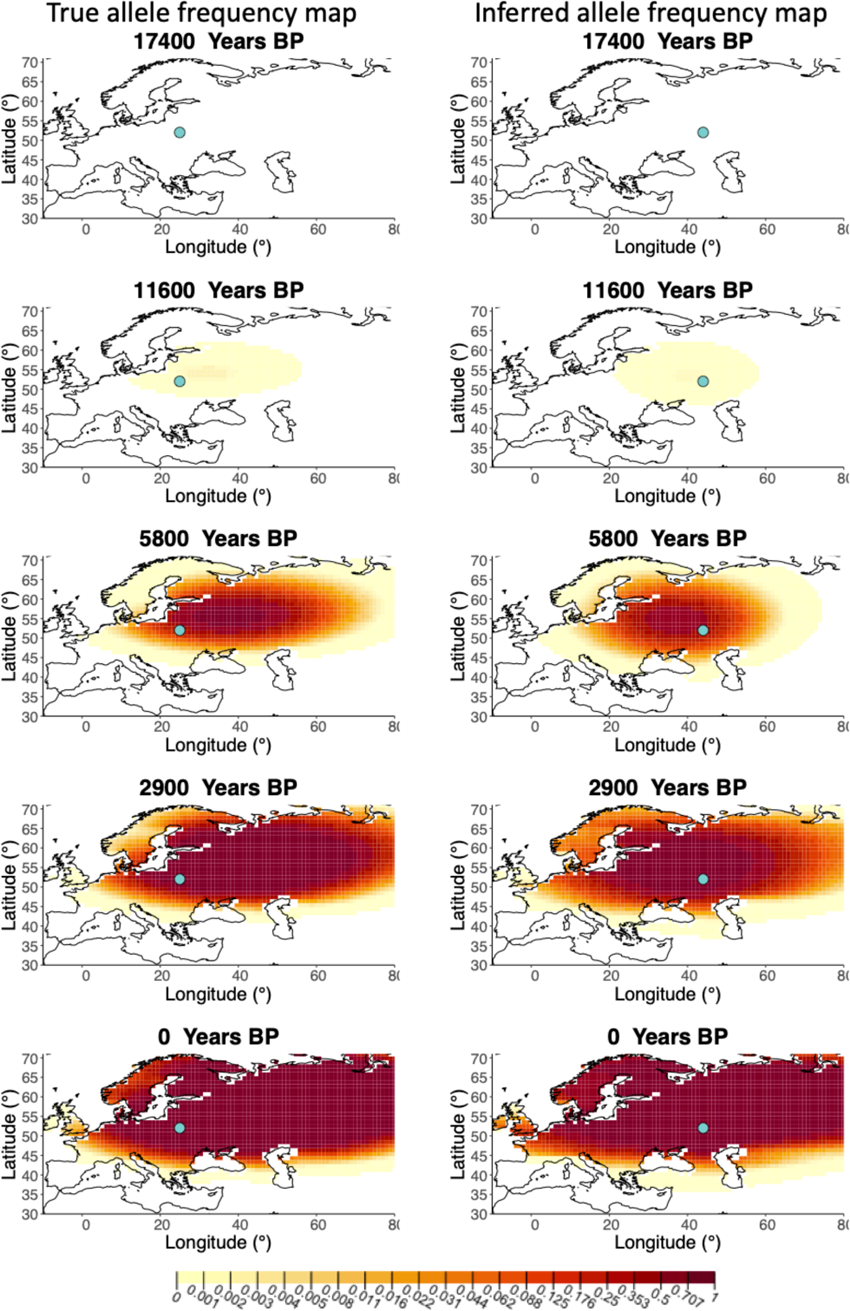
Left - Allele frequency map generated using true parameter values. Right - Allele frequency map generated using parameter estimates for “intermediate 75%/25%” clustering scheme. Parameter values used to generate the maps are summarised in ***Table A3.*** **Figure 3-Figure supplement 1.** Examples of spatial sampling scenarios for each of the three clustering schemes. **Figure 3-Figure supplement 2.** Allele frequency map generated using true parameter values and using parameter estimates for “homogeneous 75%/25%” clustering scheme **Figure 3-Figure supplement 3.** Allele frequency map generated using true parameter values and using parameter estimates for “homogeneous 50%/50%” clustering scheme **Figure 3-Figure supplement 4.** Allele frequency map generated using true parameter values and using parameter estimates for “homogeneous 25%175%” clustering scheme **Figure 3-Figure supplement 5.** Allele frequency map generated using true parameter values and using parameter estimates for “intermediate 50%/50%” clustering scheme **Figure 3-Figure supplement 6.** Allele frequency map generated using true parameter values and using parameter estimates for “intermediate 25%/75%” clustering scheme **Figure 3-Figure supplement 7.** Allele frequency map generated using true parameter values and using parameter estimates for “extreme 75%/25%” clustering scheme **Figure 3-Figure supplement 8.** Allele frequency map generated using true parameter values and using parameter estimates for “extreme 50%/50%” clustering scheme **Figure 3-Figure supplement 9.** Allele frequency map generated using true parameter values and using parameter estimates for “extreme 25%/75%” clustering scheme

### Spatially-explicit forward simulations

In addition to drawing simulated samples from a diffusion model, we used SUM ***(Haller and Messer (2019))*** to perform spatially explicit individual-basedforward-in-time simulations of selection acting on a beneficial allele, by leveraging an R interface for spatial population genetics now implemented in an R package *slendr **(Petr (2021))***.

We introduced a single beneficial additive mutation in a single individual and let it evolve across the European landscape. Before applying our method on the simulated data, we sampled 1,040 individuals whose ages were log-uniformly distributed, to ensure that there were more samples closer to the present, as in the real data. We transformed the diploid genotypes to pseudohaploid genotypes by assigning a heterozygous individual an equal probability of carrying the ancestral or the derived genotype. The parameter values estimated by our model to the simulations described in this section are summarised in ***Table A4*.**

We can see that the origin of the allele inferred by the model closely corresponds to the first observation of the derived allele in the simulation ***(Figure 4).*** The inferred selection coefficient is only slightly higher than the true value from the simulation (0.0366 vs 0.030). In general, the model accurately captures the spread of the allele centered in central Europe, though we observe some discrepancies due to differences between the model assumed in the simulation (which, for example, accounts for local clustering of individuals, ***Figure4-Figure Supplement 1),*** and that assumed by our diffusion-based inference.

**Figure 4.**
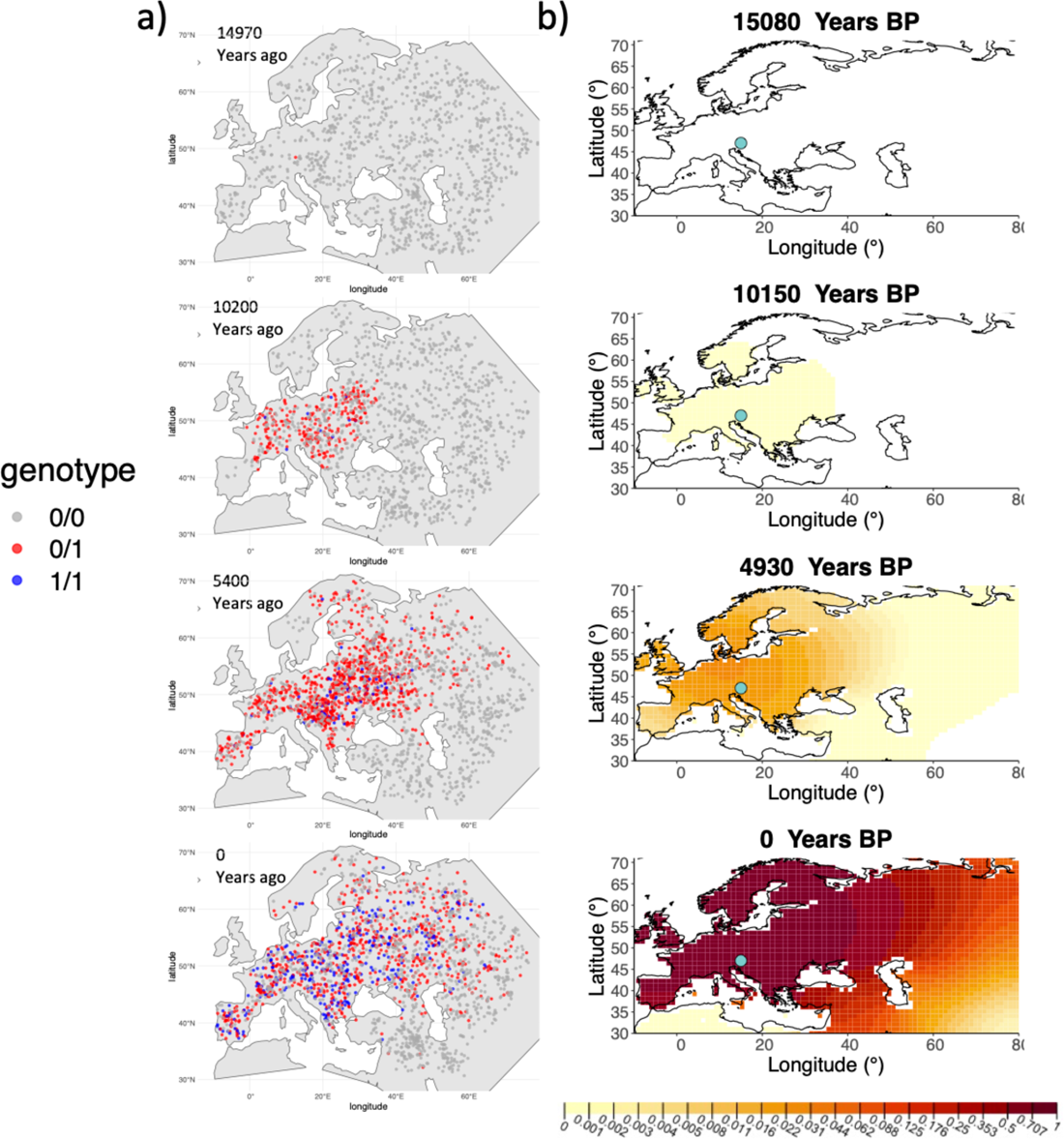
A) Individual-based simulation of an allele that arose in Central Europe 15,000 years ago with a selection coefficient of 0.03. Each dot represents a genotype from a simulated genome. To avoid overplotting, only 1,000 out of the total 20,000 individuals in the simulation in each time point are shown for each genotype category. B) Allele frequency dynamics inferred by the diffusion model on the individual-based simulation to the left, after randomly sampling 1,040 individuals from the simulation and performing pseudohaploid genotype sampling on them. The ages of sampled individuals were log-uniformly distributed. The estimated parameter values of the fitted model are shown in ***Table A4.*** **Figure 4-Figure supplement 1.** Distribution of individuals across the map under neutrality, showing the tendency of individuals to cluster together.

### Dynamics of the rs4988235(T) allele

Having tested the performance of our method on simulated data, we set out to infer the allele frequency dynamics of the rs4988235(T) allele (associated with adult lactase persistence) in ancient Western Eurasia. For our analysis, we used a genotype dataset compiled by ***Segurel et al.* (2020),** which amounts to 1,434 genotypes from ancient Eurasian genomes individuals, and a set of 36,659 genotypes from present-day Western and Central Eurasian genomes ***(Segurel and Bon, 2017; Heyer et al., 2011; Marchi et al., 2018; Liebert et al., 2017; Gallego Romero et al., 2012; /tan et al., 2010; Charati et al., 2019).*** After filtering out individuals falling outside of the range of the geographic boundaries considered in this study, we retained 1,332 ancient individuals. The locations of ancient and present-day individuals used in the analysis to trace the spread of rs4988235(T) are shown in ***Figures*.**

We used a two-period scheme by allowing the model to have two sets of estimates for the selection coefficient and the diffusion and advection coefficients in two different periods of time: before and after 5,000 years ago, reflecting the change in population dynamics and mobility before and after the Bronze Age transition ***(Loog et al., 2017; Racimo et al., 2020a).*** We used two allele age estimates as input: a relatively young one (7,441 years ago) obtained from ***/tan et al.* (2009),** and a relatively old one (20,106 years ago) obtained from ***Albers and Mcvean* (2020).** The results obtained for fitting the model on rs4988235(D are summarised in ***Table AS*** and ***Table A6,*** and in ***Figure 6b*** (younger age) and ***Figure 6-Figure Supplement 6*** (older age).

**Figure 5.**
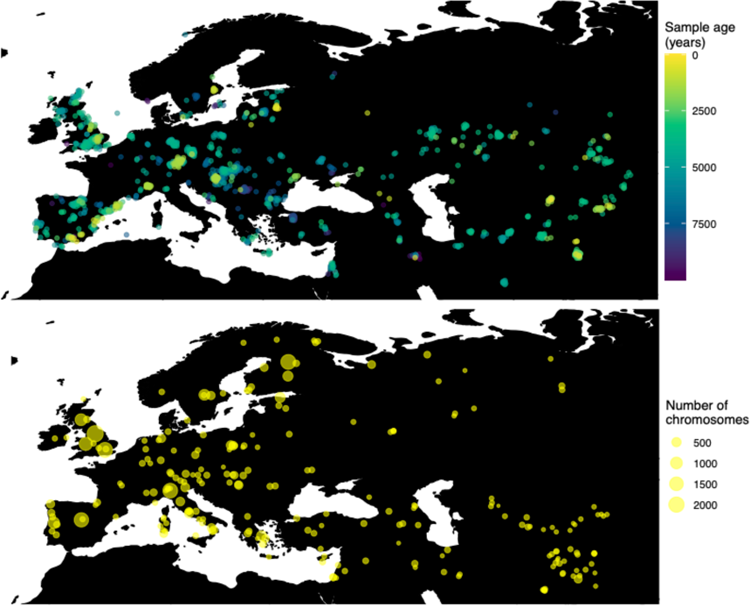
Locations of samples used to model the spread of the rs4988235(T) allele. The upper panel shows the spatiotemporal locations of ancient individuals, the bottom panel represents the locations of present-day individuals.

**Figure 6.**
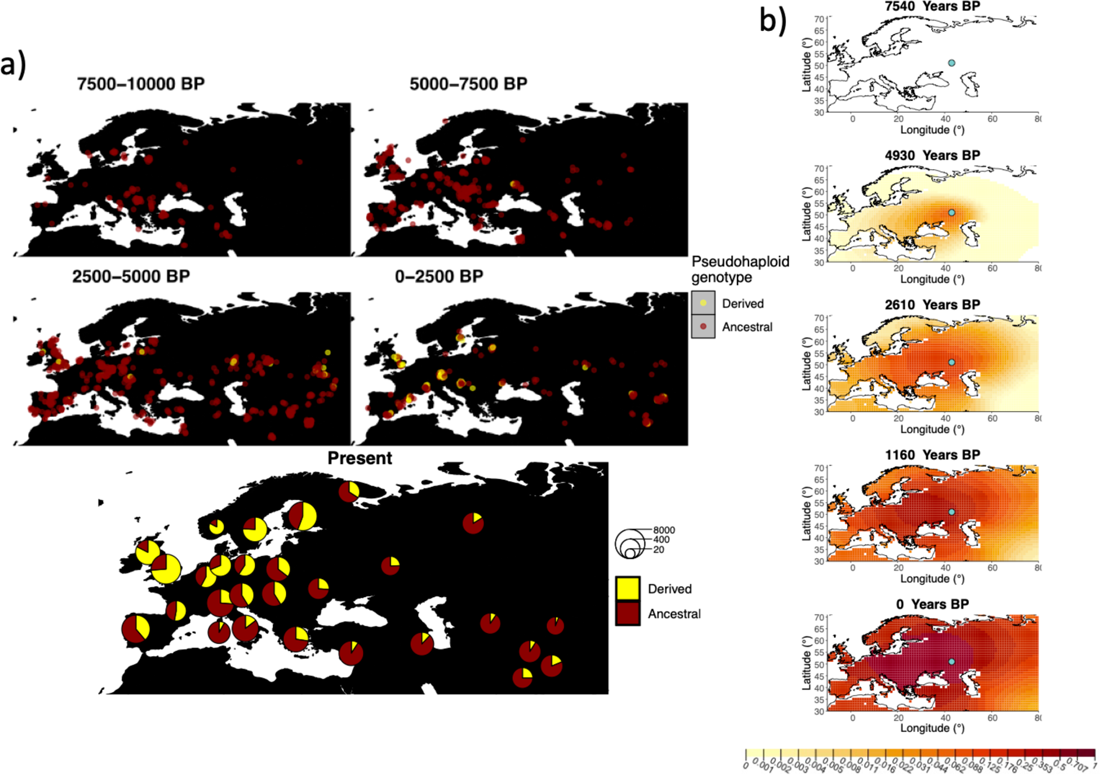
a) Top: Pseudohaploid genotypes of ancient samples at the rs4988235 SNP in different periods. Yellow corresponds to the rs4988235(T) allele. Bottom: allele frequencies of present-day samples represented as pie charts. The size of the pie charts corresponds to the number of available sequences in each region. b) Inferred allele frequency dynamics of rs4988235(T). The green dot indicates the inferred geographic origin of the allele. **Figure 6-Figure supplement 1.** Inferred frequency dynamics of rs4988235(T) using the allele age that was inferred in ***Albers and Mcvean* (2020).** **Figure 6-Figure supplement 2.** Inferred frequency dynamics of rs4988235(T) when the origin of the allele is moved 10 degrees west from the original estimate. **Figure 6-Figure supplement 3.** Inferred frequency dynamics of rs4988235(T) when the origin of the allele is moved 10 degrees east from the original estimate. **Figure 6-Figure supplement 4.** Inferred frequency dynamics of rs4988235(T) when the origin of the allele is moved 10 degrees north from the original estimate. **Figure 6-Figure supplement 5.** Inferred frequency dynamics of rs4988235(T) when the origin of the allele is moved 10 degrees south from the original estimate. **Figure 6-Figure supplement 6.** Inferred frequency dynamics of rs4988235(T) forcing the geographic origin of the allele to be at the location inferred in ***/tan et al.* (2009).** **Figure 6-Figure supplement 7.** Inferred frequency dynamics of rs4988235(T) assuming the allele age to be the lower end of the 95%credible interval for the allele age inferred in ***/tan et al.* (2009).** **Figure 6-Figure supplement 8.** Inferred frequency dynamics of rs4988235(T) assuming the allele age to be the higher end of the 95%credible interval for the allele age inferred in ***/tan et al.* (2009).** **Figure 6-Figure supplement 9.** Log-likelihood values for model runs using different ages of the rs4988235(T) allele as input.

Assuming the age estimate from ***(tan et al.,* 2009),** the origin of the allele is estimated to be north of the Caucasus, around what is now southwestern Russia and eastern Ukraine ***(Figure 6b*).** Given that this age is relatively young, our method fits a very strong selection coefficient(0.1) during the first period in order to accommodate the early presence of the allele in various points throughout Eastern Europe, and a weaker (but still strong) selection coefficient(0.03) in the second period. We also estimate stronger diffusion in the second period than in the first, to accommodate the rapid expansion of the allele throughout Western Europe, and a net westward advection parameter, indicating movement of the allele frequency’s center of mass to the west as we approach the present.

Assuming the older age estimate from ***Albers and Mcvean* (2020),** the origin of the allele is estimated to be in the Northeast of Europe ***(Figure 6-Figure Supplement 1),*** which is at a much higher latitude than the first occurrence of the allele, in Ukraine. Due to the deterministic nature of the model, the frequency is implicitly imposed to expand in a region where there are no actual observed instances of the allele. The model compensates for this by placing the origin in an area with a lower density of available aDNA data and thus avoiding an overlap of the increasing allele frequencies with individuals who do not carry the derived rs4988235(T) allele (see ***Figure 6a*).** As the model expands rapidly in the southern direction ***(Table A6)*** it eventually reaches the sample carrying the derived variant in Ukraine.

### Dynamics of the rs1042602(A) allele

Next, we investigated the spatiotemporal dynamics of the spread of an allele at a pigmentationassociated SNP in the *TYR* locus (rs1042602(A)), which has been reported to be under recent selection in Western Eurasian history ***(Stern et al., 2019).*** For this purpose, we applied our method to the Allen Ancient DNA Resource data ***(Reich and Mallick, 2019),*** which contains randomly sampled pseudohaploid genotypes from 1,513 published ancient Eurasian genomes (listed **in** Supplementary File 1), from which we extracted those genomes that had genotype information at this locus in Western Eurasia. We merged this dataset with diploid genotype information from high-coverage present-day West Eurasian genomes from the Human Genome Diversity Panel (HGDP) ***(Bergstrom et al., 2020),*** which resulted in a total of 1,040 individuals with genotype information at rs1042602, which were as input to our analysis. Geographic locations of individuals in the final dataset are shown in ***Figure* 7.**

**Figure 7.**
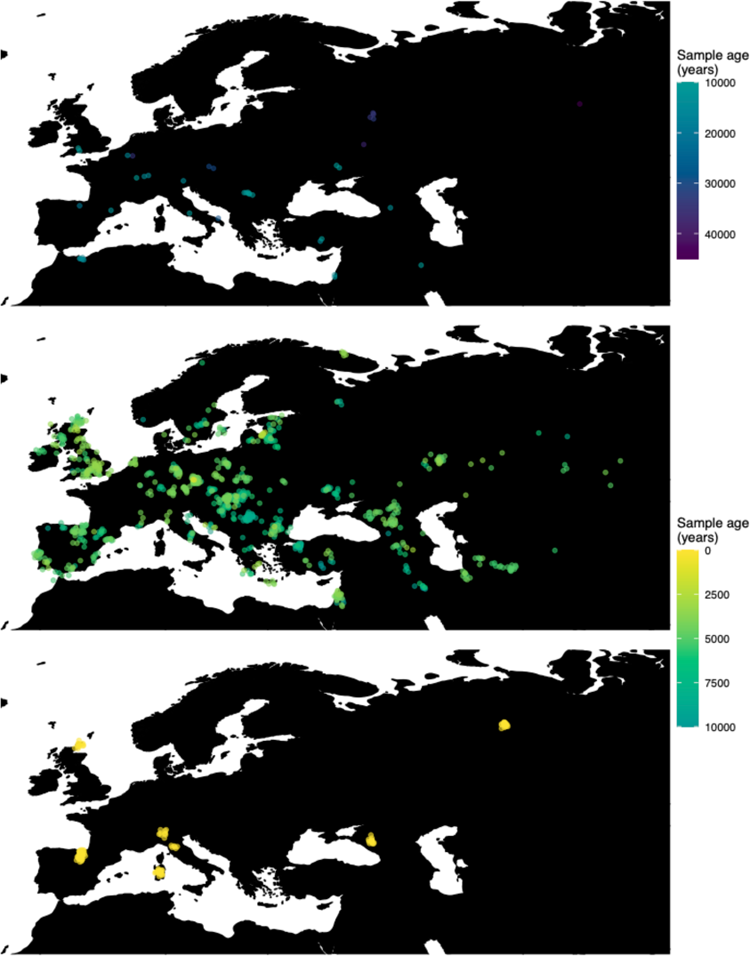
Spatiotemporal sampling locations of sequences used to model the rs1042602(A) allele in Western Eurasia. Upper panel: ancient individuals dated as older than 10,000 years ago. Middle panel: ancient individuals dated as younger than 10,000 years ago. Bottom panel: present-day individuals from HGDP.

Similarly to our analysis of the spread of the allele in rs4988235(T), we inferred the dynamics of the rs1042602(A) allele separately for the time periods before and after 5,000 years BP and assuming the age of the allele to be 26,361 years ***(Albers and Mcvean, 2020).*** The inferred parameters for both time periods are summarised in ***Table Al*** and the allele frequency surface maps generated using these parameters are shown in ***Figure* Bb.** The origin of the rs1042602(A) corresponds closely to the region where the allele initially starts to segregate in the time period between 7,500 and 10,000 years BP as seen in ***Figure Ba.*** Estimates of the selection coefficient for both time periods (0.0221 and 0.0102 for the period before and after 5000 years BP, respectively) suggest that selection acting on the allele has decreased after 5000 years BP.

### Robustness of parameters to the inferred geographic origin of allele

We carried out an analysis to characterize how sensitive the selection, diffusion and advection parameters are to changes in the assumed geographic origin of the allele. For the rs4988235(T) allele, we forced the origin of the allele to be 10 degrees away from our inferred origin in each cardinal direction, while assuming the allele age from ***/tan et al.* (2009) *(Table AB).*** In ***Figure 6-Figure Supplement 6, Figure 6-Figure Supplement 6, Figure 6-Figure Supplement 6*** and ***Figure 6-Figure Supplement 6,*** we can see the allele frequency dynamics of these four scenarios, respectively. We also forced the allele origin to be at the geographic origin estimated in ***/tan et al.* (2009) *(Table A9, Figure 6-Figure Supplement 6),*** which is westward of our estimate. In all five cases during the period prior 5,000 years BP, the allele is inferred to expand in the direction of the first sample that is observed to carry the rs4988235(T) allele and is located in Ukraine. During the time period after 5000 years BP, the patterns produced by the model are rather similar, although the parameters associated with diffusion and advection differ, in order to account for the different starting conditions.

We also investigated how the results are affected when the estimated geographic origin of the rs1042602(A) allele is moved with respect to the initial estimate. We set the allele to be 10 degrees east, 10 degrees north and 10 degrees south of the original estimate as shown in ***Figure 8-Figure Supplement 8, Figure 8-Figure Supplement 8*** and ***Figure 8-Figure Supplement 8,*** respectively. We did not look at a scenario in which the origin of the allele is moved to the west, since it would either end up in the Black sea or more westwards than 10 degrees. The selection coefficient remains similar to the original estimate throughout all three scenarios. The way the allele spreads across the landscape is also similar in all cases and, as in the case of rs4988235(T), the model accounts for the different origins of the allele by adjusting the diffusion and advection coefficients in the time period after 5000 years BP.

**Figure 8.**
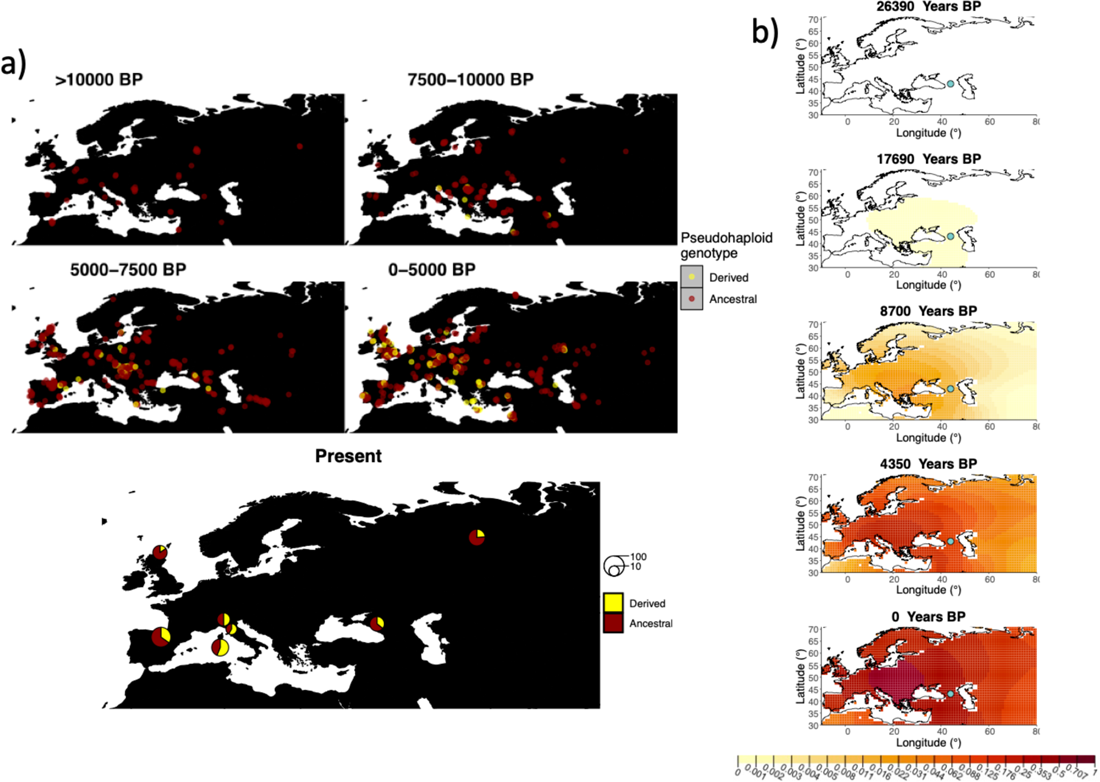
a) Top: Pseudohaploid genotypes of ancient samples of the rs1042602 in different periods. Yellow corresponds to the A allele. Bottom: diploid genotypes of present-day samples. b) Inferred allele frequency dynamics of rs1042602(A). The green dot corresponds to the inferred geographic origin of the allele. **Figure 8-Figure supplement 1.** Inferred frequency dynamics of rs1042602(A) when the origin of the allele is moved 10 degrees east from the original estimate. **Figure 8-Figure supplement 2.** Inferred frequency dynamics of rs1042602(A) when the origin of the allele is moved 10 degrees north from the original estimate. **Figure 8-Figure supplement 3.** Inferred frequency dynamics of rs1042602(A) when the origin of the allele is moved 10 degrees south from the original estimate. **Figure 8-Figure supplement 4.** Inferred frequency dynamics of rs1042602(A) assuming the allele age to be the lower end of the 95%confidence interval for the allele age inferred in ***Albers and Mcvean (2020).*** **Figure 8-Figure supplement 5.** Frequency dynamics of rs1042602(A) assuming the allele age to be the higher end of the 95%confidence interval for the allele age inferred in ***Albers and Mcvean (2020).*** **Figure 8-Figure supplement 6.** Log-likelihood values for model runs using different ages of the rs1042602(A) allele as input.

### Robustness of parameters to the assumed age of the allele

In order to investigate how sensitive our inferences are to the point estimates of allele ages we obtained from the literature ***(Albers and Mcvean, 2020; /tan et al., 2009),*** we also fitted our model using the upper and lower ends of the 95% confidence intervals or credible intervals for each age estimate (depending on whether the inference procedure in the literature was via a maximum likelihood or a Bayesian approach). For the rs4988235(T) allele, the reported credible intervals for the ***/tan et al. (2009)*** age are 8,683 and 6,256 years BP. For the rs1042602(A) allele, the reported confidence intervals for the age are 27,315 and 25,424 years BP ***(Albers and Mcvean,* 2020).**

When re-fitting the model for the rs4988235(T) allele, we found that the inferred selection coefficient is slightly lower when the allele age is assumed to be at the lower bound of the 95% credi· ble interval (0.0867 vs 0.0993 before 5000 years BP and 0.0321 vs 0.0328 after 5000 years BP) and slightly higher when assumed to be at the upper bound (0.0994 vs 0.0993 before 5000years BP and 0.0572 vs 0.0328 after 5000 years BP) ***(Table AS*** and ***Figure 6-Figure Supplement 7*** and ***Figure 6-Figure Supplement 6).*** This occurs because the selection intensity must be higher or lower when there is more or less time, respectively, for the allele to reach the allele frequencies observed in the data. In the case of the rs1042602(A) allele, this only affects the earlier time period ***(Table A]).*** The rs4988235(T) allele’s geographic distribution in the more recent time periods is also less extended geographically when the age is assumed to be young. The inferred geographic origin of both alleles slightly differs under different assumed ages ***(Figure 8-Figure Supplement 8*** and ***Figure 8-Figure Supplement 5*).**

In addition, we assessed the likelihood of the best fitted models with varying the ages of the rs4988235(T) and rs1042602(A) alleles ***(Figure 6-Figure Supplement 6*** and ***Figure 8-Figure Supplement 6,*** respectively). We can see that in the case of rs4988235(T) allele the allele age used in this study (7,441 years) gives the most likely solution among the explored ages. In case of the rs1042602(A) allele, we found that there are multiple nearly equally likely ages when looking at ages at least as old as 15,000 years.

## Discussion

A spatially explicit framework for allele frequency diffusion can provide new insights into the dynamics of selectedvariants across a landscape. We have shown that under the conditions of strong, recent selection, our method can infer selection and dispersal parameters, using a combination of ancient and present-day human genomic data. However, when allowing for advection, the inferred location tends to become less accurate. This suggests that migration events early in the dispersal of the selected allele could create difficulties in finding the true allele origin if net directional movement (i.e. via major migratory processes) had a large effect in this dispersal. This issue could be alleviated with the inclusion of more ancient genomes around the time of the origin of the mutation, perhaps in combination with a more fine-scaled division into periods where advection may have occurred in different directions.

The inferred geographic origin of the rs4988235(T) allele reflects the best guess of our framework given the constraints provided by its input, namely the previously inferred age of the allele and the observed instances of this allele throughout Western Eurasia. We are also assuming that the allele must have arisen somewhere within the bounding box of our studied map. When assuming a relatively young allele age (7,441 years ago, ***/tan et al.* (2009)),** the origin of the allele is placed north of the Caucasus, perhaps among steppe populations that inhabited the area at this time ***(Haak et al., 2015; Allentoft et al., 2015).*** This origin is further east than the geographic origin estimate from ***/tan et al.* (2009),** likely reflecting additional ancient DNA information that is available to us, and indicates an early presence of the allele in eastern Europe. When assuming a relatively old allele age (20,106 years ago, ***Albers and Mcvean* (2020)),** the age is placed in northeast Europe, perhaps among Eastern hunter-gatherer groups that inhabited the region in the early Holocene. We note that the number of available genomes for eastern and northeastern Europe during the early Holocene is scarce, so the uncertainty of the exact location of this origin is relatively high. Regardless of the assumed age, we estimate a net westward displacement of the allele frequency’s center of mass, and a rapid diffusion, particularly in the period after 5,000 years ago.

Various studies have estimated the selection coefficient for the rs4988235(T) allele, and these range from as low as 0.014 to as high as 0.19 ***(Enattah et al., 2008; Mathieson and Mathieson, 2018; Mathieson, 2020; Stern et al., 2019; Burger et al., 2020; Peter et al., 2012; Gerbault et al., 2009; /tan et al., 2009; Bersaglieri et al., 2004).*** Recent papers incorporating ancient DNA estimate the selection coefficient to be as low as O (in certain regions of Southern Europe) and as high as 0.06 ***(Mathieson and Mathieson, 2018;Mathieson, 2020;Burger et al.,* 2020).** It isalso likely that the selection coefficient was different for different regions of Europe, perhaps due to varying cultural practices ***(Mathieson,* 2020).** In our case, the estimated selection coefficient during the first period - before 5,000 years ago - depends strongly on the assumed allele age (s = 0.0993 vs. s = 0.0285). As in the case of the geographic origin, these estimates should be taken with caution as the number of available allele observations in the early Holocene is fairly low. The estimates for the second period - after 5,000 years ago - are more robust to the assumed age: s = 0.0328 (95% Cl: 0.0327-0.0329) if we assume the younger allele age (7,441 years ago) and s = 0.0255 (95% Cl: 0.0252-0.0258) if we assume the older allele age (20,106 years ago). These estimates are also within the range of previous estimates.

In the case of the rs1042602(A) allele, our estimated selection coefficients of 0.0221 (95% Cl: 0.0216-0.0227) and 0.0102 (95% Cl: 0.0083-0.0120) for the time periods before and after 5000 years BP, respectively, are generally in agreement with previous results. ***Wilde et al. (2014)*** used a forward simulation approach to infer a point estimate of 0.026. Another study using an approximate Bayesian computation framework ***(Nakagome et al., 2019)*** estimated the strength of selection acting on rs1042602 to be 0.013 (0.002-0.029). Although both studies utilized ancient DNA data, the estimates were obtained without explicitly modelling the spatial dimension of the selection process.

Our estimates of the longitudinal advection parameter are negative for both the SNPs in the *TYR* and *LCT* loci: the mutation origins are always to the east of the center of mass of the allele frequency distribution seen in present-day data. This perhaps reflects common migratory processes, like the large-scale Neolithic and Bronze Age population movements from east to west, affecting the allele frequencies at these loci across the Eurasian landscape ***(A/lentoft et al.,2015; Haak et al.,2015).*** As a form of regularization, we kept the range of explored values for the advection parameters to be small (−2.5 to 2.5 km per generation), while allowing the diffusion parameters to be explored over a much wider range of values. In certain cases, like the second period of the rs4988235(T) spread when the allele age is assumed to be young ***(Table AS),*** we find that the advection parameters are fitted at the boundary of the explored range, because the allele needs to spread very fast across the landscape to fit the data.

A future improvement to our method could include other forms of regularization that better account for the joint behavior of the advection and diffusion processes, or the use of priors for these parameters under a Bayesian setting, which could be informed by realistic assumptions about the movement of individuals on a landscape. Bayesian parameter fitting would likely provide a more robust understanding of the uncertainty of the estimates as well as an opportunity to formally compare different models using Bayes factors, although at the cost of an increase of computational intensity.

When investigating the robustness of the geographic origin of both rs4988235(D and rs1042602(A), we found that parameters related to the beneficial allele’s expansion change in response to different assumed origins of the allele. The resulting allele frequency surface plots, however, appear very similar throughout the later stages of the process, showing that the model tends to adjust the diffusion and advection coefficients in a way such that the allele will end up expanding into the same areas regardless of the origin.

As we apply these methods to longer time scales and broader geographic areas, the assumptions of spatiotemporal homogeneity of the parameters seem less plausible. There may be cases where the allele may have been distributed over a wide geographic area but remained at low frequencies for an extended period of time, complicating the attempts to pinpoint the allele’s origin. In our study, we estimated diffusion and selection coefficients separately for two time periods before and after 5000 years ago to account for changes in mobility during the Bronze Age, but this approach may still be hindered by uneven sampling, especially when the allele in question exists at very low frequencies. Notably, our results for the spread of the rs4988235(T) allele during the older time period should be interpreted with caution, since they may be affected by sparse sampling in the early Holocene.

Potential future extensions of our method could incorporate geographic features and historical migration events that create spatially or temporally varying moderators of gene flow. An example of this type of processes is the retreat of glaciers after the last Glacial maximum, which allowed migration of humans into Scandinavia ***(Giinther et al., 2018).*** These changing geographic features could lead to changes in the rate of advection or diffusion across time or space. They could also serve to put more environmentally-aware constraints on the geographic origin of the allele, given that it cannot have existed in regions uninhabitable by humans, and to extend our analyses beyond the narrow confines of the Western Eurasian map chosen for this study. One could also envision incorporating variation in population densities over time, or known migration processes in the time frames and regions of interest. These might have facilitated rapid, long-range dispersal of beneficial alleles ***(Bradburd et al., 2016; Hallatschek and Fisher, 2014)*** or caused allelic surfing on the wave of range expansions ***(Klop/stein et al., 2006).*** Additional information like this could come, for example, from previously inferred spatiotemporal demographic processes (e.g. ***Racimo et al. (2020b))*.**

As described above, our model only accounts for diffusion in two directions. Further extension of our model could therefore incorporate anisotropic diffusion ***(Othmer et al., 1988; Painter and Hillen, 2018).*** Another possibility could be the introduction of stochastic process components, in order to convert the partial differential equations into stochastic differential equations ***(Brown et al.,* 2000).** Stochastic components could serve to induce spatial autocorrelation and capture local patterns of allele frequency covariance in space that might not be well modeled by the deterministic PDEs ***(Cressie and Wikle, 2015).*** They could also serve to induce stochasticity in allele frequency changes over time as a consequence of genetic drift ***(Crow et al., 1970),*** allowing one to model the dynamics of more weakly selected variants, where drift plays an important role. Eventually, one could perhaps combine information across loci to jointly model the spatiotemporal frequency surfaces at multiple loci associated with the same trait. This could help clarify the dynamics of polygenie adaptation and negative selection on complex traits ***(Irving-Pease et al., 2021),*** and perhaps hindcast the genetic value of traits across a landscape.

The availability of hundreds of ancient genomes ***(Marciniak and Perry, 2017)*** and the increasing interest in spatiotemporal method development ***(Bradburd and Ralph, 2019),*** such as the one described in this manuscript, will likely lead researchers to posit new questions and hypotheses about the behavior of natural selection. In the case of a beneficial allele spreading on a landscape, new ontologies and vocabulary for describing positive selection in time and space will be needed.

Abundant terms exists to classify the initial conditions and dynamics of a selective sweep in a single population (hard sweep, multiple origin soft sweep, single origin soft sweep, partial sweep) ***(Hermisson and Pennings, 2005; Pritchard and Di Rienzo, 2010; Hermisson and Pennings, 2017).*** In contrast, there is a lack of vocabulary for distinguishing between a scenario of strong selection that is locally constrained in space from a scenario of widespread selection extended over a landscape, or a model of neutral diffusion in space followed by parallel non-neutral increases in frequency at multiple locations. For example, ***Ralph and Coop (2010)*** showed how multiple localized hard sweeps may be seen as a soft sweep at a larger population-wide scale. Existing vocabulary for spatiotemporal genetic processes is clearly not enough, limiting the types of questions or hypotheses we can pose about them.

Population genetic models that explicitly account for space and time are an important area of future methodological development ***(Bradburd and Ralph, 2019).*** We believe that methods such as the one described in this study show great promise at broadening the horizon of our understanding of natural selection across space and time in humans and other species. As in the case of demographic reconstruction ***(Ray and Excoffier,* 2009),** spatiotemporal information can greatly help improve our knowledge of how natural selection operated in the past.

## Methods

### The model

To describe the allele frequency dynamics in time and space, we first begin by usinga deterministic model based on a two-dimensional partial differential equation (PDE) ***(Fisher, 1937; Kolmogorov et al., 1937; Novembre et al.,* 2005).** This PDE represents the distribution *p(x, y,t)* of the allele frequency across a two dimensional *(x,* y) landscape at time t:

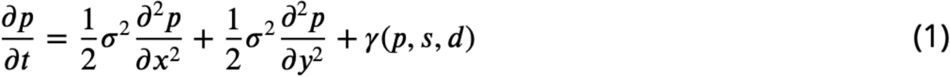

where

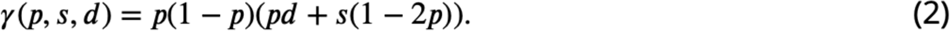

Here, *u* is the diffusion coefficient, *s* is the selection coefficient, and *d* is the dominance coefficient ***(Novembre et al.,2005).*** We assumed an additive model and fixed *d* = 2s in all analyses below. We call this “model A”, but we also evaluated the fit of our data under more complex models which are more flexible, and are described below.

Model B is a more general diffusion-reaction model, which incorporates distinct diffusion terms in the longitudinal and latitudinal directions *(ux* and *uy,* respectively):

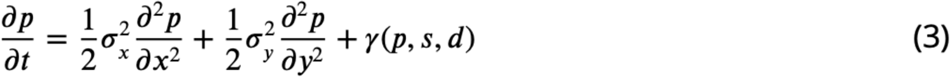

Model C is a generalization of model B that incorporates advection terms in the longitudinal and latitudinal directions (see e.g. ***Cantrell and Cosner* (2004)** for a motivation of this type of model in the context of spatial ecology):

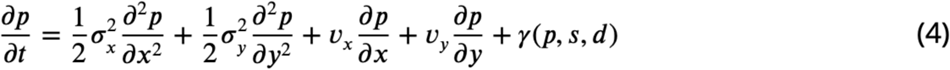

Here, *v_x_* and *v_Y_* represent the coefficients for advective velocity along the longitude and latitude respectively.

In the Appendix, we motivate the construction of these equations using model C as an example, and show that equation 4 can be obtained by taking an infinitesimal limit of a random walk on a two-dimensional lattice, after including a reaction term due to selection. Models A and Bare then shown to be special cases of model C.

For evaluating the likelihood of the observed data, we use a binomial genotype sampling model. Let *g_1_* E 0,1,2 be the genotype of individual i at the locus of interest, let *a_1_* be the number of reads carrying ancestral alleles, let *d_1_* be the number of reads carry derived reads. Let *(x_1_, y;)* be the coordinates of the location from which individual i was sampled, and *1_1_* its estimated age (e.g. from radiocarbon dating). Then, the likelihood for individual i can be computed as follows:

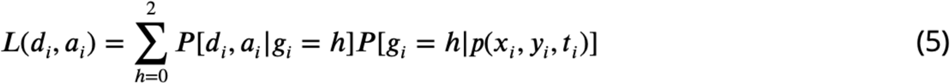

Here, *p(x_i_, y_i_, t_i_)* is the solution to one of the partial differential equations described above (equations (1), (2) or (4), depending on the process model chosen), evaluated at location *(x_i_, y_i_)* and time t*_i_*. In turn, *P[d_i_,a_i_lg_i_* = h] is the likelihood for genotype i. Furthermore, *P[g_i_* = *hlp(x_i_,y_i_,t_i_)]* is a binomial distribution, where *n* represents the ploidy level, which in this case is 2:

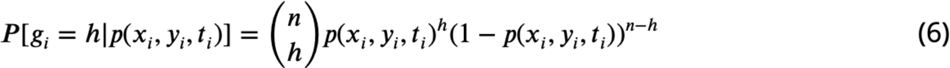

Then, the likelihood of the entire data can be computed as

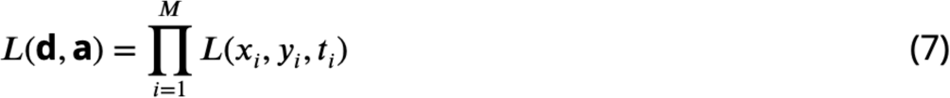

where **M** is the total number of individuals for which we have data, **d** is the vector containing the derived read count for each individual and **a** is the vector containing the ancestral read count for each individual. We computed genotype likelihoods directly on the BAM file read data, using the SAMtools genotype model ***(Li, 2011)*** implemented in the software ANGSD ***(Korneliussen et al., 2014)*.**

When only randomly sampled pseudohaploid allele counts are available, we used a Bernoulli sampling likelihood (conditional on the genotype *g)* on the right-hand side of equation 6 instead. Briefly, assuming that the probability of an individual having genotype g at a particular locus given the underlying allele frequency *p* follows a binomial distribution and that the probability of samplinga read given the genotype of an individual follows a Bernoulli distribution with probability of success g, then the probability of sampling a read given the genotype follows a Bernoulli distribution with probability of success *p*.

### Map

We restricted the geographic area explored by our model fit to be between 30°N to 75°N, and between 10°w and 80°E. For numerical calculations, we used a grid constructed using a resolution of approximately 1 grid cell per latitude and longitude. We used Harvesine functions in order to transform the distance from degrees to kilometers between two geographic points. The diffusion of the allele frequency was disallowed in the map regions where the topology is negative (i.e. regions under water), based on ETOPO5 data ***(NOAA (1988)).*** For this reason we added land bridges between the European mainland and Sardinia, and between the mainland and Great Britain, in order to allow the allele to diffuse in these regions (see ***Figure A1)*.**

### Parameter search

Parameter optimization was done via maximum likelihood estimation with a two-layer optimization set-up. The first layer consists of a simulated annealing approach ***(Belisle (1992))*** starting from 50 random points in the parameter space. The initial 50 points are sampled using latin hypercube sampling to ensure an even spread across the parameter space. The output of this fit was then fed to the L-BFGS-B algorithm to refine the parameter estimates around the obtained maximum and obtain confidence intervals for the selection, diffusion and advection parameters ***(Byrd et al. (1995))*.**

The parameters optimised were:

- the selection coefficient (s), restricted to the range 0.001-0.1
- two dispersal parameters *<Jx* and *<JY* in the longitudinal and latitudinal directions respectively, restricted to the range of 1-100 square-kilometers per generation
- the longitudinal and latitudinal advection coefficients *vx* and *vY* respectively. As a form of regularization, we set the range of explored values to be narrowly centered around zero: −2.5 to 2.5 kilometers per generation
- the geographic origin of the allele, which is randomly initialized to be any of the 28 spatial points shown in ***Figure A2*** at the start of the optimization process

We chose to construct our method in a way that uses the age of the allele as an input parameter rather than estimating it. We do this since there are multiple equally possible solutions with various combinations of allele age and selection coefficient values as shown in ***Figure A3.*** The latitude and longitude are discretized in our model in order to solve the differential equations numerically, thus the origin of a mutation is measured in terms of discrete units. For this reason, when using the L-BFGS-B algorithm, we fixed the previously estimated origin of the allele, and did not explore it during this second optimization layer. For numerical calculations we used the Livermore Solver for Ordinary Differential Equations ***(Hindmarsh, 1983)*** implemented in R package “deSolve” ***(Soetaert et al., 2010a),*** which is a general purpose solver that can handle both stiff and nonstiff systems. In case of stiff problems the solver uses aJacobian matrix. Absorbing boundary conditions were used at the boundaries of the map. For visualisation purposes we masked the allele frequencies from areas with negative topology (i.e. areas covered by large bodies of water). Time was measured in generations, assuming 29 years per generation. During the optimization we scaled the time and the parameters by a factor of 10, which allowed us to decrease the execution time of the model.

We initialized the grid by setting the initial allele frequency to be *p_0_* in a grid cell where the allele originates and O elsewhere. *p_0_* was calculated as 1/(2* *D* * *A),* where *D* is the population density and is equal to 2.5 inhabitants per square-kilometer, which is the estimated population density in Europe in 1000 B.C. ***(Colin McEvedy, 1978; Novembre et al., 2005).*** In the equation, Dis multiplied by 2 because we assume that the allele originated in a single chromosome in a diploid individual. *A* is the area in square-kilometers of the grid cell where the allele emerged.

Asymptotic 95% confidence intervals for a given parameter *o_i_* were calculated using equation

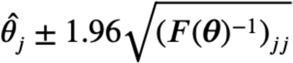

where *F(θ)* is an estimate of the observed Fisher information matrix ***(Fisher, 1922; Efron and Hastie, 2016; Casella and Berger, 2021)*.**

### Implementation

The above described model was implemented in R version 3.6. To numerically solve the differential equations and obtain maximum likelihood estimates, we used the libraries *deSo/ve* ***(Soetaert et al., 2010b),*** *ReacTran* ***(Soetaert and Meysman, 2012)*** and *bbmle* ***(Balker and R Development Core Team,* 2020).** Scripts containing the code used in this paper are available on github: https://github.com/RasaMukti/stepadna

### lndvidual-based simulations

For the individual-based spatiotemporal forward simulations, we first defined a spatial boundary for a population spread across a broad geographic region of Europe. In order to ensure a reasonably uniform distribution of individuals across this spatial range throughout the course of the simulation, we set the maximum distance for spatial competition and mating choice between individuals to 250 km (translated, on a SUM level, to the interaction parameter *maxDistance),* and the standard deviation of the normal distribution governing the spread of offspring from their parents at 25 km (leveraged in SLiM’s *modifyChild()* callback function) ***(Haller and Messer, 2019).*** We note that we have chosen the values of these parameters merely to ensure a uniform spread of individuals across a simulated landscape. They are not intended to represent realistic estimates for these parameters at any time in human history.

After defining the spatial context of the simulations and ensuring the uniform spread of individuals across their population boundary, we introduced a single beneficial additive mutation in a single individual. In order to test how accurately our model can infer the parameters of interest, we simulated a scenario in which the allele appeared in Central Europe 15,000years ago with the selection coefficient of the beneficial mutation set to 0.03. Over the course of the simulation, we tracked the position of each individual that ever lived together with its location on a two-dimensional map, as well as its genotype (i.e. zero, one, or two copies of the beneficial allele). We then used this complete information about the spatial distribution of the beneficial allele in each time point to study the accuracy of our model in inferring the parameters of interest.

## Acknowledgments

We thank Graham Gower, Evan Irving-Pease, Montgomery Slatkin, the members of the Racimo group and two anonymous reviewers for helpful comments and advice. FR and RM were funded by a Viiium Fonden Young Investigator award to FR (project no. 00025300). MP and FR were supported by a Lundbeckfondengrant (R302-2018-2155)and a Novo Nordisk Fonden grant (NNF18SA0035006) to the GeoGenetics Centre. TSK was funded by a Carlsberg grant (CF19-0712). JN was funded by NIH grant R01 GM132383.

## Appendix

### Appendix 1

Here, we motivate the construction of model Casa large scale limit of a random walk model on a lattice ***(Karlin and Taylor, 1975; Cantrell and Cosner, 2004).*** We think of the allele frequency as a variable *p* that can increase in magnitude due to its inherent advantage (selection), spread across a landscape (diffusion) or move directionally as a consequence of migration (advection). We imagine a lattice composed of small square cells of size *t:u* x *dy,* where a certain amount of allele frequency *p* can occur at a given time point *t.* At each small time step (of duration dt), inflow and outflow of p can occur in the x-direction with probability h or in they-direction with probability 1-h, and the magnitude of these flows depend on the amount of *p* present in neighboring cells. If flow of pis along the x-axis, it does so in the positive direction with probability *a* and in the negative direction with probability **1-a**.

If flow of p is along the y-axis, it does so in the positive direction with probability *p* and in the negative direction with probability 1 - *p.* The allele frequency can also increase in magnitude locally, via a function rO that depends on its dominance (d), selection coefficient (s) and current magnitude *(p(x, y,*t)). Then, we obtain:

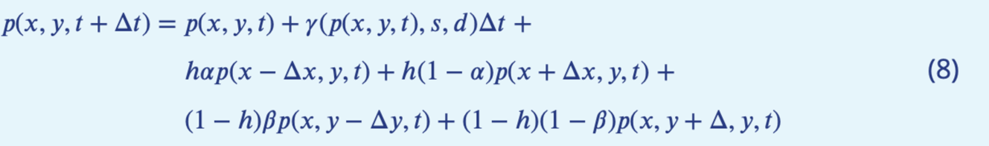

We can also write this as:

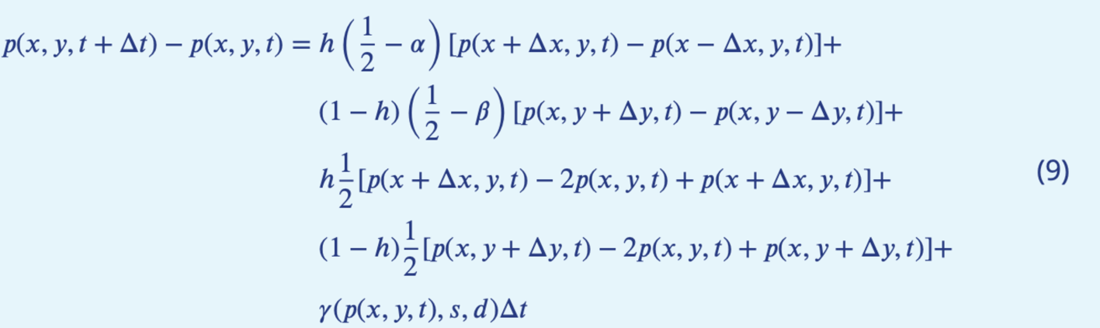

If we divide both sides by *dt* and take the limit of infinitesimally small *Δx, Δy* and *Δt,* while assuming that, in this limit, 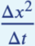 and 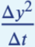 are finite ***(Okubo et al., 1980),*** we obtain:

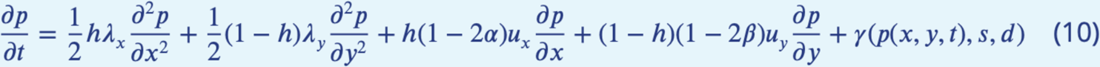

where 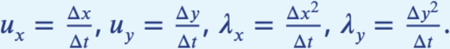

If we let 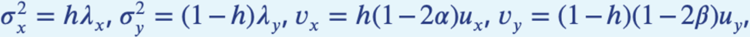 then we obtain equation 4. Thus, we can see that the squared diffusion coefficient σ^2^_x_ depends on the square of the length of the cells in the x-axis relative to the duration of a time step (λ_x_), and on the probability that flows occurs in the x-axis at a given time step (*h*). Similarly, the squared diffusion coefficient σ^2^_y_ depends on the square of the length of the cells in the y-axis relative to the duration of a time step (λ_y_), and on the probability that flows occurs in the y-axis at a given time step (1 - h). The advection coefficient *v_x_* depends on the advective velocity along the x-axis (u_x_) as well as on the probability of flow occurring along the x-axis (h) and the directional bias 1 *-2α,* which depends on the probability that flow occurs in the positive x-direction (a). Finally, the advection coefficient *vy* depends on the advective velocity along the y-axis *(u_y_)* as well as on the probability of flow occurring along the y-axis (1 - h) and the directional bias 1 - *2β,* which depends on the probability that flow occurs in the positive y-direction (β).

We can recover model Bas a special case of model C ifwe fix *α = α = 1/2,* assuming isotropy n ht two directions, so that Δx = Δy. We can also recover model A if we additionally fix *h* = 1/2.

## Appendix 2

**Figure A1.**
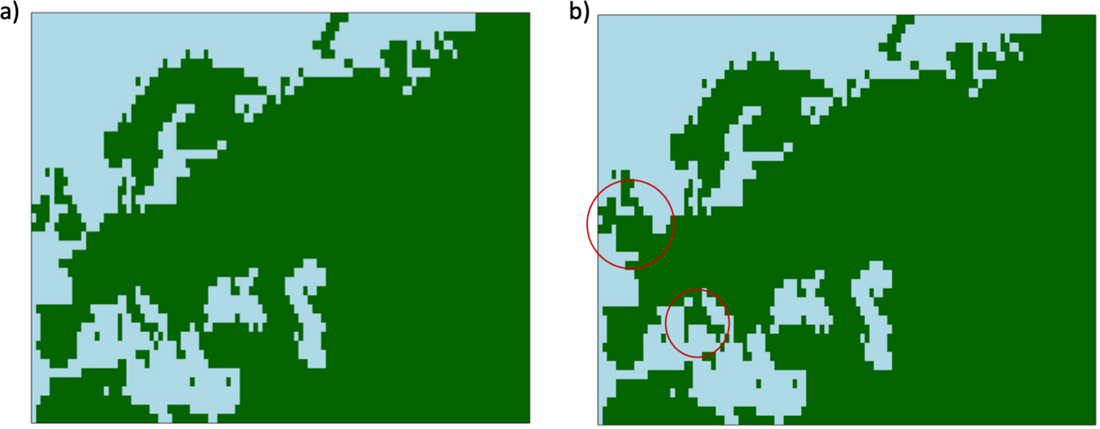
Maps showing areas where diffusion in the model is allowed {green) and where it is forbidden {blue). Figure a) map without land bridges. Figure b) map containing land bridges indicated with red circles.

**Figure A2.**
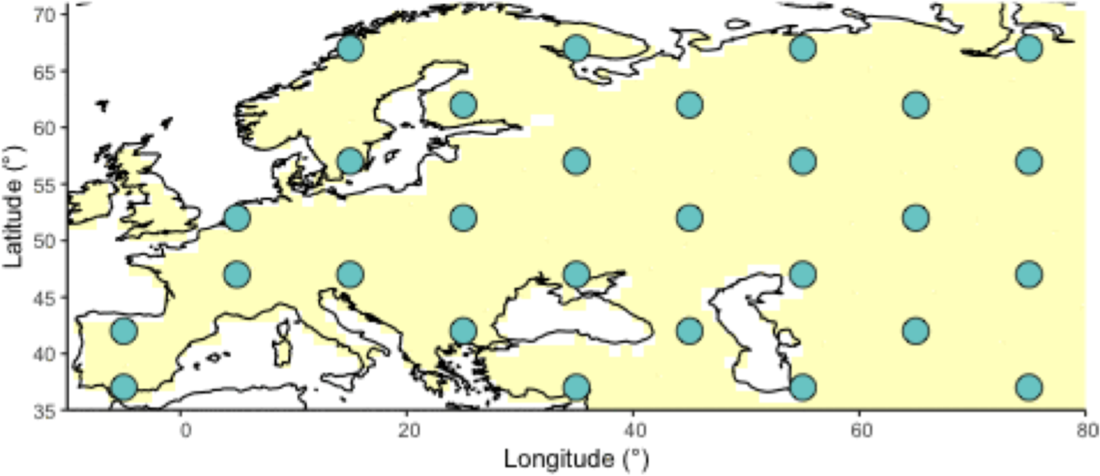
Geographic locations for points used as potential origins of the allele at the initialization of the simulated annealing optimization algorithm. Note that, after initialization, the algorithm can continuously explore any points on the map grid that are not necessarily included in this point set

**Figure A3.**
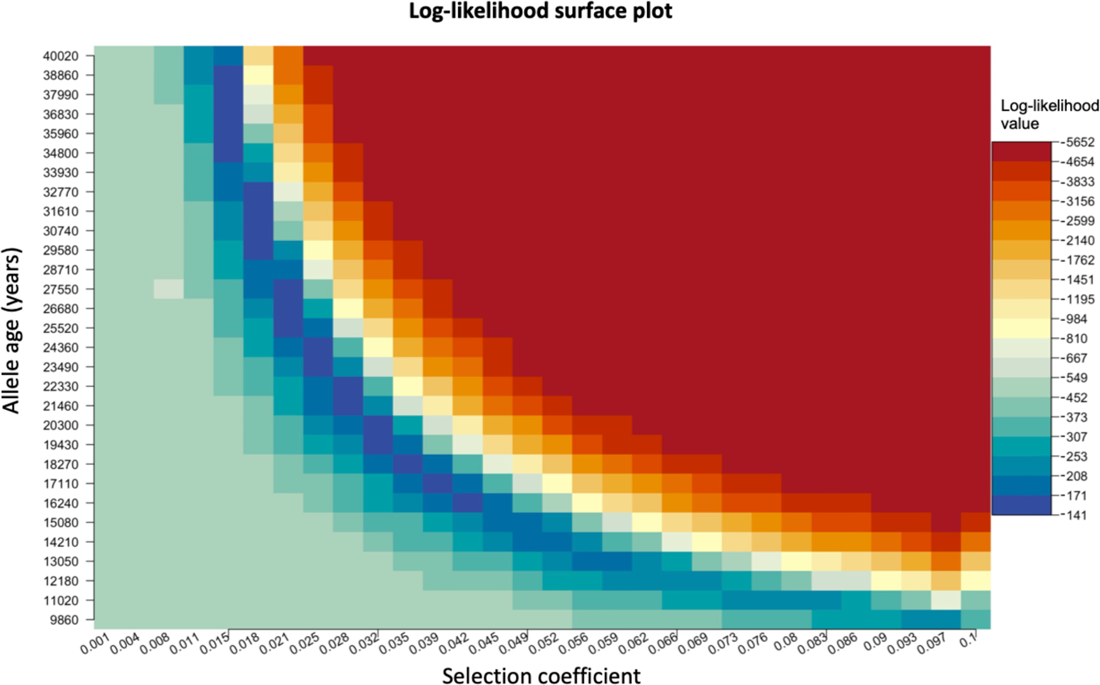
Log-likelihood as a function of selection coefficient and age of the allele. Dark blue regions correspond to optimal solutions.

**Table A1.** Parameter values used to generate simulations using numerical solutions to equation 3, compared to parameter estimates assuming model B. The age of the allele was set to 29,000 years in all simulations. Columns named “Long” and “Lat” indicate the longitude and latitude of the geographic origin of the allele, respectively.

| Simulation | | $s$ | $\sigma_x$<br>(km/gen) | $\sigma_y$<br>(km/gen) | Long | Lat |
| --- | --- | --- | --- | --- | --- | --- |
|  |  | True/Pred<br>(95% CI) | True/Pred<br>(95% CI) | True/Pred<br>(95% CI) | True/Pred | True/Pred |
| B1 | Sample age >5000 | 0.02/0.0192<br>(0.0187–0.0196) | 10/15.244<br>(2.5042–27.9828) | 20/16.963<br>(11.9263–21.9993) | 25/24 | 52/52 |
|  | Sample age <5000 | 0.02/0.0027<br>(0–0.0074) | 10/8.805<br>(0.5631–17.0468) | 20/97.432<br>(97.2566–97.6081) | – | – |
| B2 | Sample age >5000 | 0.02/0.0193<br>(0.0189–0.0198) | 10/15.348<br>(0–95.5192) | 30/20.427<br>(0–51.514) | 25/25 | 52/51 |
|  | Sample age <5000 | 0.02/0.001<br>(0–0.0059) | 10/10.015<br>(1.6837–18.3472) | 30/100<br>(99.9933–100.0067) | – | – |
| B3 | Sample age >5000 | 0.02/0.0196<br>(0.0191–0.02) | 10/8.149<br>(6.1551–10.143) | 40/49.432<br>(0–135.9428) | 25/24 | 52/51 |
|  | Sample age <5000 | 0.02/0.0265<br>(0.0145–0.0386) | 10/7.855<br>(0–19.751) | 40/100<br>(99.9933–100.0067) | – | – |
| B4 | Sample age >5000 | 0.02/0.0188<br>(0.0188–0.0188) | 20/19.037<br>(19.0311–19.0435) | 10/19.254<br>(19.2439–19.2638) | 25/25 | 52/51 |
|  | Sample age <5000 | 0.02/0.0142<br>(0.0111–0.0173) | 20/17.354<br>(4.4083–30.2991) | 10/1.489<br>(0–18.5993) | – | – |
| B5 | Sample age >5000 | 0.02/0.0196<br>(0.019–0.0202) | 30/26.409<br>(11.1997–41.6184) | 10/11.429<br>(7.1825–15.6759) | 25/27 | 52/51 |
|  | Sample age <5000 | 0.02/0.0215<br>(0.0183–0.0246) | 30/14.3<br>(0–68.176) | 10/1.554<br>(0–16.5985) | – | – |
| B6 | Sample age >5000 | 0.02/0.0199<br>(0.0192–0.0206) | 40/85.415<br>(41.6058–129.2248) | 10/9.02<br>(7.2853–10.7538) | 25/6 | 52/51 |
|  | Sample age <5000 | 0.02/0.0163<br>(0.0112–0.0213) | 40/10.403<br>(0–22.533) | 10/22.623<br>(12.0841–33.1614) | – | – |

**Table A2.** Parameter values used to generate simulations using numerical solutions to equation 4, compared to parameter estimates assuming model C. The age of the allele was set to 29,000 years in all simulations. Columns named “Long” and “Lat” indicate the longitude and latitude of the geographic origin of the allele, respectively.

| Simulation | | $s$ | $\sigma_x$<br>(km/gen) | $\sigma_y$<br>(km/gen) | $v_x$<br>(km/gen) | $v_y$<br>(km/gen) | Long | Lat |
| --- | --- | --- | --- | --- | --- | --- | --- | --- |
|  |  | True/Pred<br>(95% CI) | True/Pred<br>(95% CI) | True/Pred<br>(95% CI) | True/Pred<br>(95% CI) | True/Pred<br>(95% CI) | True/Pred | True/Pred |
| C1 | Sample age >5000 | 0.02/0.0189<br>(0.0188–0.0189) | 20/52.246<br>(52.2051–52.2872) | 20/16.373<br>(16.332–16.4139) | -2/-1.675<br>(-1.6771–-1.6722) | -2/-2.067<br>(-2.0702–-2.0639) | 25/4 | 52/45 |
|  | Sample age <5000 | 0.02/0.0231<br>(0.023–0.0233) | 20/11.086<br>(10.9286–11.2441) | 20/15.606<br>(15.3659–15.8467) | -2/0.399<br>(0.3946–0.4037) | -2/-2.491<br>(-2.5458–-2.436) | – | – |
| C2 | Sample age >5000 | 0.02/0.0185<br>(0.0176–0.0195) | 10/19.434<br>(5.6736–33.1952) | 20/18.727<br>(4.8938–32.5605) | -1.2/1.579<br>(1.2671–1.8905) | 1.9/-0.801<br>(-1.1684–-0.4331) | 25/-6 | 52/38 |
|  | Sample age <5000 | 0.02/0.0205<br>(0.0175–0.0234) | 10/38.144<br>(10.3123–65.9749) | 20/51.094<br>(14.0489–88.1388) | -1.2/-1.299<br>(-2.7247–0.1266) | 1.9/2.493<br>(2.4929–2.4933) | – | – |
| C3 | Sample age >5000 | 0.02/0.0255<br>(0.0254–0.0256) | 30/59.237<br>(59.1269–59.347) | 10/6.604<br>(6.5991–6.6087) | 1.8/2.195<br>(2.1918–2.1985) | -0.8/0.438<br>(0.4381–0.4387) | 25/65 | 52/66 |
|  | Sample age <5000 | 0.02/0.0079<br>(0–0.0165) | 30/86.511<br>(0–194.0772) | 10/40.693<br>(24.3946–56.9905) | 1.8/-2.498<br>(-2.4983–-2.498) | -0.8/-0.014<br>(-3.4481–3.4204) | – | – |
| C4 | Sample age >5000 | 0.02/0.0197<br>(0.0191–0.0204) | 10/19.647<br>(14.975–24.3197) | 10/13.585<br>(0–27.2936) | 1.2/-0.054<br>(-0.0968–-0.0111) | 1/0.72<br>(0.4278–1.0124) | 25/44 | 52/50 |
|  | Sample age <5000 | 0.02/0.0137<br>(0.0046–0.0229) | 10/14.151<br>(0–32.1031) | 10/4.093<br>(0–49.4651) | 1.2/0.8<br>(-3.4903–5.0895) | 1/2.434<br>(2.4299–2.4387) | – | – |

**Table A3.**
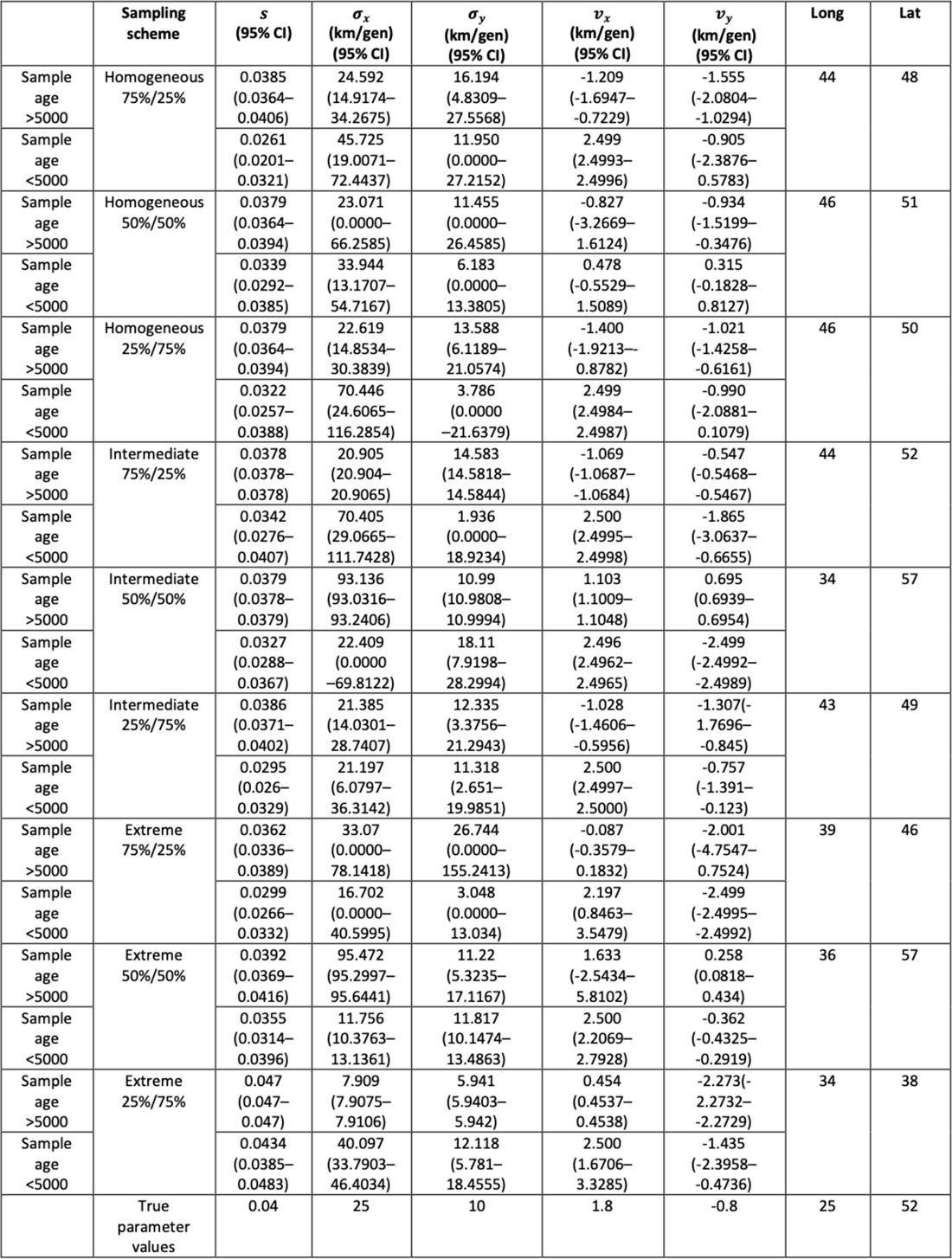
Parameter value estimates for each of the 9 clustering schemes and true parameter values used to generate the deterministic simulation. The age of the allele was set to 17,400 years. Columns named “Long” and “Lat” indicate the longitude and latitude of the geographic origin of the allele, respectively.

**Table A4.** Parameter values estimated using model C for the forward simulation created using SUM. Columns named “Long” and “Lat” indicate the longitude and latitude of the geographic origin of the allele, respectively.

| $s$<br>(95% CI) | $\sigma_x$<br>(km/gen)<br>(95% CI) | $\sigma_y$<br>(km/gen)<br>(95% CI) | $v_x$<br>(km/gen)<br>(95% CI) | $v_y$<br>(km/gen)<br>(95% CI) | Long | Lat | Allele age<br>(years) |
| --- | --- | --- | --- | --- | --- | --- | --- |
| 0.0366<br>(0.0357–0.0375) | 58.583<br>(49.1983–67.9669) | 63.733<br>(3.6601–123.8056) | -0.436<br>(-0.8077–-0.0649) | -1.564<br>(-3.0915–-0.0355) | 15 | 47 | 15000 |

**Table A5.**
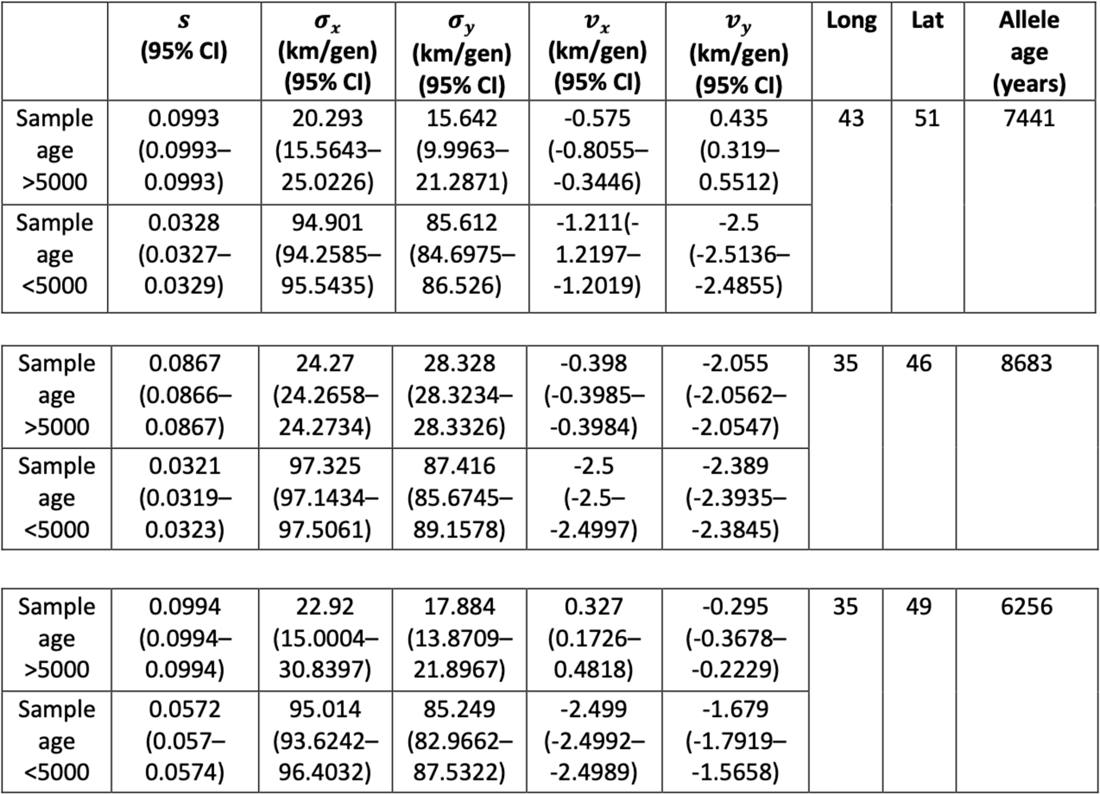
Summary of parameter estimates for rs4988235(T). The upper two rows correspond to results obtained assuming the allele age to be the point estimate from ***/tan et al.* (2009):** 7,441 years ago. The middle two rows and the bottom two rows show results assuming the age to be either the lower or the higher ends of the allele age’s 95%confidence interval from ***/tan et al.* (2009).** Columns named “Long” and “Lat” indicate the longitude and latitude of the geographic origin of the allele, respectively.

**Table A6.** Parameter estimates for rs4988235(T) using the allele age inferred in ***Albers and Mcvean* (2020).** Columns named “Long” and “Lat” indicate the longitude and latitude of the geographic origin of the allele, respectively.

| | $s$<br>(95% CI) | $\sigma_x$<br>(km/gen)<br>(95% CI) | $\sigma_y$<br>(km/gen)<br>(95% CI) | $v_x$<br>(km/gen)<br>(95% CI) | $v_y$<br>(km/gen)<br>(95% CI) | Long | Lat | Allele<br>age<br>(years) |
| --- | --- | --- | --- | --- | --- | --- | --- | --- |
| Sample<br>age<br>>5000 | 0.0285<br>(0.0285–<br>0.0285) | 1.25<br>(1.2492–<br>1.25) | 44.619<br>(44.5944–<br>44.6445) | -0.177<br>(-0.1773–<br>-0.1771) | 1.925<br>(1.9247–<br>1.9262) | 32 | 66 | 20106 |
| Sample<br>age<br><5000 | 0.0255<br>(0.0252–<br>0.0258) | 92.545<br>(91.6963–<br>93.3941) | 87.545<br>(85.3525–<br>89.7369) | -2.499<br>(-2.4992–<br>-2.4989) | -2.271<br>(-2.4127–<br>-2.1297) |  |  |  |

**Table A7.** Summary of parameter estimates for rs1042602(A). Upper two rows corresponds to model fit when allele age is set to be the point estimate ***Albers and Mcvean* (2020):** 26,367 years ago. The middle two rows and the bottom two rows show results assuming the age to be either the lower or the higher ends of the allele age’s 95%confidence interval from ***Albers and Mcvean* (2020).** Columns named “Long” and “Lat” indicate the longitude and latitude of the geographic origin of the allele, respectively.

| | $s$<br>(95% CI) | $\sigma_x$<br>(km/gen)<br>(95% CI) | $\sigma_y$<br>(km/gen)<br>(95% CI) | $v_x$<br>(km/gen)<br>(95% CI) | $v_y$<br>(km/gen)<br>(95% CI) | Long | Lat | Allele<br>age<br>(years) |
| --- | --- | --- | --- | --- | --- | --- | --- | --- |
| Sample<br>age<br>>5000 | 0.0221<br>(0.0216–<br>0.0227) | 71.668<br>(24.7274–<br>118.6092) | 50.434<br>(25.6535–<br>75.2136) | -2.268<br>(-3.006–<br>-1.5304) | -0.486<br>(-0.8661–<br>-0.1053) | 44 | 43 | 26367 |
| Sample<br>age<br><5000 | 0.0102<br>(0.0083–<br>0.012) | 69.25<br>(14.0247–<br>124.4756) | 95.281<br>(95.1087–<br>95.453) | 0.849<br>(-0.0783–<br>1.7769) | -0.503<br>(-0.929–<br>-0.076) |  |  |  |
| Sample<br>age<br>>5000 | 0.0214<br>(0.0205–<br>0.0223) | 57.914<br>(0–<br>131.3177) | 83.846<br>(0–<br>246.6688) | -2.111<br>(-2.8784–<br>-1.3429) | 1.305<br>(-0.8411–<br>3.4519) | 46 | 51 | 27315 |
| Sample<br>age<br><5000 | 0.01<br>(0.0078–<br>0.0121) | 88.218<br>(0–<br>190.105) | 96.216<br>(96.0422–<br>96.3898) | 1.19<br>(-0.7489–<br>3.1293) | -0.88<br>(-2.0897–<br>0.3299) |  |  |  |
| Sample<br>age<br>>5000 | 0.023<br>(0.023–<br>0.0231) | 75.857<br>(75.8065–<br>75.9071) | 48.992<br>(48.9166–<br>49.0674) | -2.362<br>(-2.3655–<br>-2.3593) | -0.837<br>(-0.8371–<br>-0.8362) | 43 | 42 | 25424 |
| Sample<br>age<br><5000 | 0.0099<br>(0.0085–<br>0.0112) | 72.847<br>(67.7991–<br>77.8949) | 92.867<br>(75.4925–<br>110.2412) | 0.497<br>(0.2717–<br>0.7214) | -0.685<br>(-0.8076–<br>-0.5628) |  |  |  |

**Table A8.**
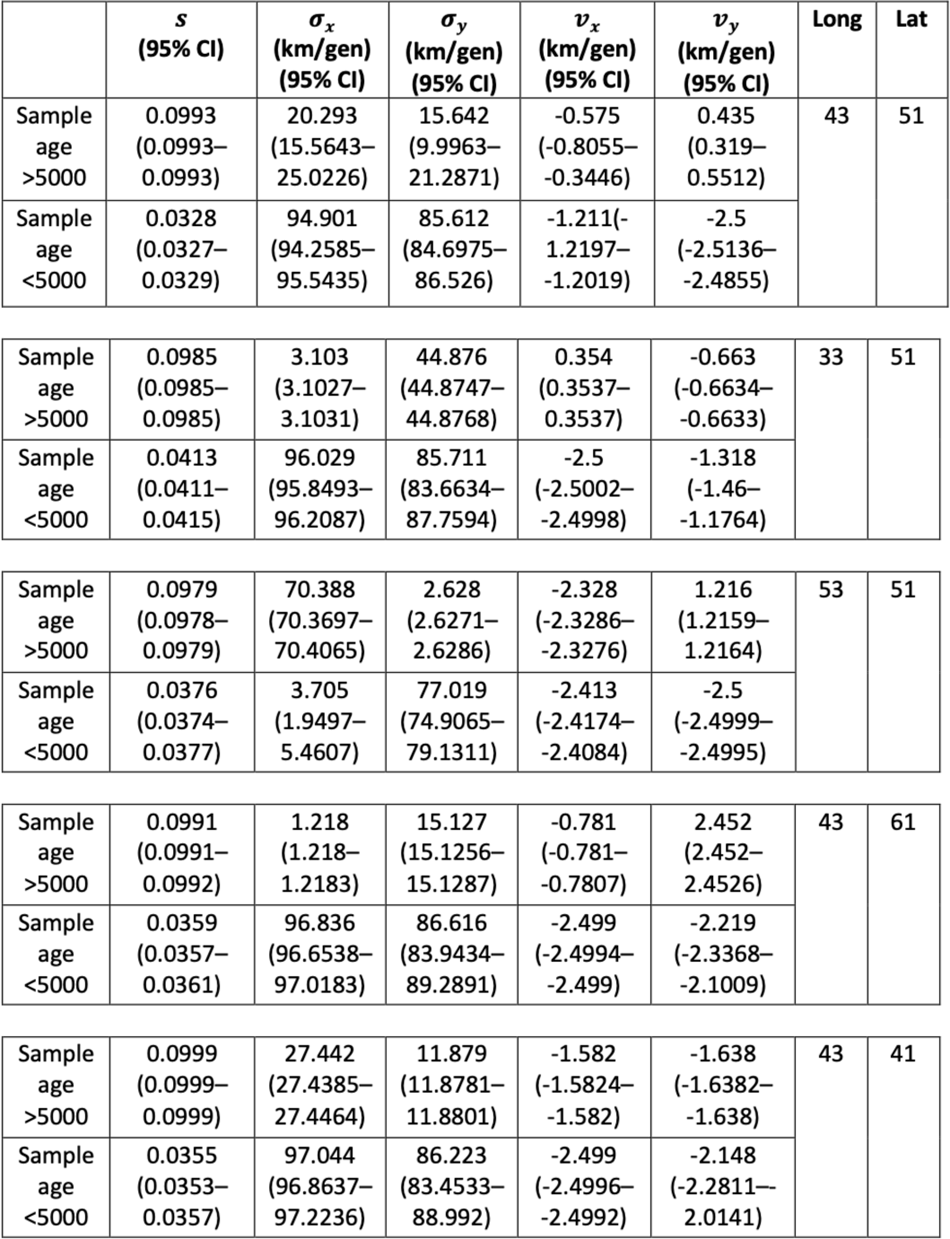
Summary of parameter estimates for rs4988235(T) when the origin of the allele is forced to be at different points in the map (top panel corresponds to the original fit for the geographic position). In all cases, the estimated age of allele that was inputted into the model is 7,441 years ago. Columns named “Long” and “Lat” indicate the longitude and latitude of the geographic origin of the allele, respectively.

**Table A9.**
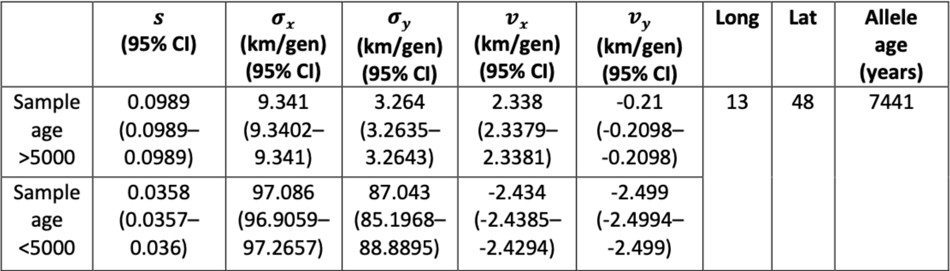
Parameter estimates for rs4988235(T) using the geographic origin of the allele inferred in ***/tan et al.* (2009).** Columns named “Long” and “Lat” indicate the longitude and latitude of the geographic origin of the allele, respectively.

**Table A10.** Summary of parameter estimates for rs1042602(A) when the origin of the allele is forced to be at different points in the map (top panel corresponds to the original fit for the geographic position). In all cases, the estimated age of allele that was inputted into the model is 26,367 years ago. Columns named “Long” and “Lat” indicate the longitude and latitude of the geographic origin of the allele, respectively.

| | $s$<br>(95% CI) | $\sigma_x$<br>(km/gen)<br>(95% CI) | $\sigma_y$<br>(km/gen)<br>(95% CI) | $v_x$<br>(km/gen)<br>(95% CI) | $v_y$<br>(km/gen)<br>(95% CI) | Long | Lat |
| --- | --- | --- | --- | --- | --- | --- | --- |
| Sample age >5000 | 0.0221<br>(0.0216–0.0227) | 71.668<br>(24.7274–118.6092) | 50.434<br>(25.6535–75.2136) | -2.268<br>(-3.006–-1.5304) | -0.486<br>(-0.8661–-0.1053) | 44 | 43 |
| Sample age <5000 | 0.0102<br>(0.0083–0.012) | 69.25<br>(14.0247–124.4756) | 95.281<br>(95.1087–95.453) | 0.849<br>(-0.0783–1.7769) | -0.503<br>(-0.929–-0.076) |  |  |
| Sample age >5000 | 0.0227<br>(0.0223–0.0231) | 42.745<br>(33.6354–51.8541) | 96.993<br>(96.8183–97.1683) | -2.437<br>(-2.4412–-2.4324) | -0.266<br>(-0.4848–-0.0468) | 54 | 43 |
| Sample age <5000 | 0.0095<br>(0.007–0.0119) | 93.477<br>(7.6582–179.2965) | 99.634<br>(0–205.4586) | -2.499<br>(-3.2101–-1.7873) | 2.057<br>(-0.7888–4.903) |  |  |
| Sample age >5000 | 0.0221<br>(0.0221–0.0221) | 47.691<br>(47.6686–47.7127) | 71.367<br>(71.336–71.3986) | -2.164<br>(-2.1652–-2.1637) | 1.839<br>(1.8387–1.8392) | 44 | 53 |
| Sample age <5000 | 0.0112<br>(0.0093–0.0131) | 87.959<br>(0–215.8939) | 88.951<br>(25.5422–152.3589) | 2.108<br>(-0.2061–4.4227) | -2.237<br>(-5.7828–1.3083) |  |  |
| Sample age >5000 | 0.0219<br>(0.0209–0.0229) | 73.106<br>(38.1699–108.043) | 76.835<br>(24.0025–129.6684) | -2.429<br>(-2.4335–-2.4248) | -1.474<br>(-2.8769–-0.0706) | 44 | 33 |
| Sample age <5000 | 0.0102<br>(0.0083–0.0121) | 88.216<br>(0–192.1057) | 95.401<br>(95.2283–95.573) | 0.871<br>(-0.2474–1.9893) | -1.026<br>(-2.6161–0.564) |  |  |

---

**Figure 1-Figure supplement 1.**
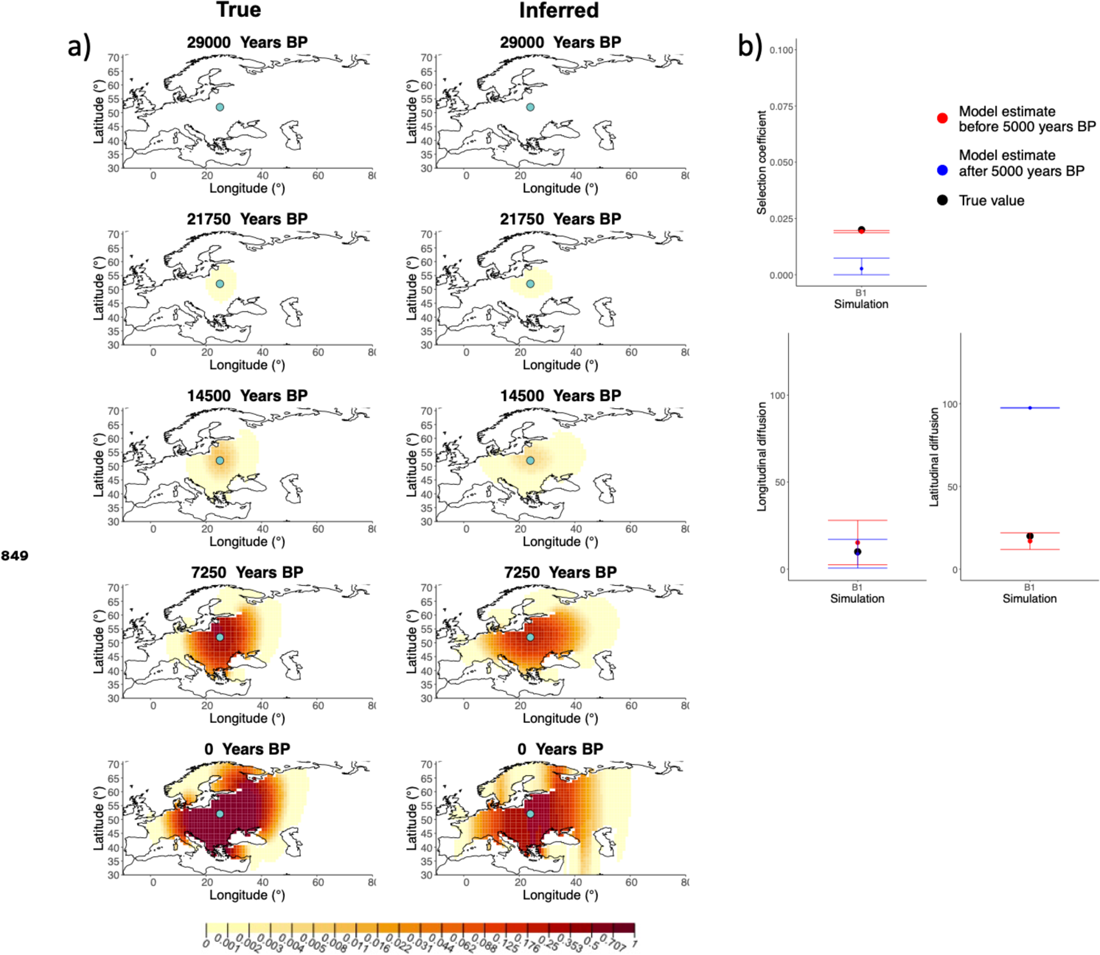
a) Comparison of true and inferred allele frequency dynamics for simulation B1. The green dot corresponds to the origin of the allele. The parameter values used to generate the frequency surface maps are summarised in **Table A1.** b) Comparison of true parameter values and model estimates. Whiskers represent 95% confidence intervals.

**Figure 1-Figure supplement 2.**
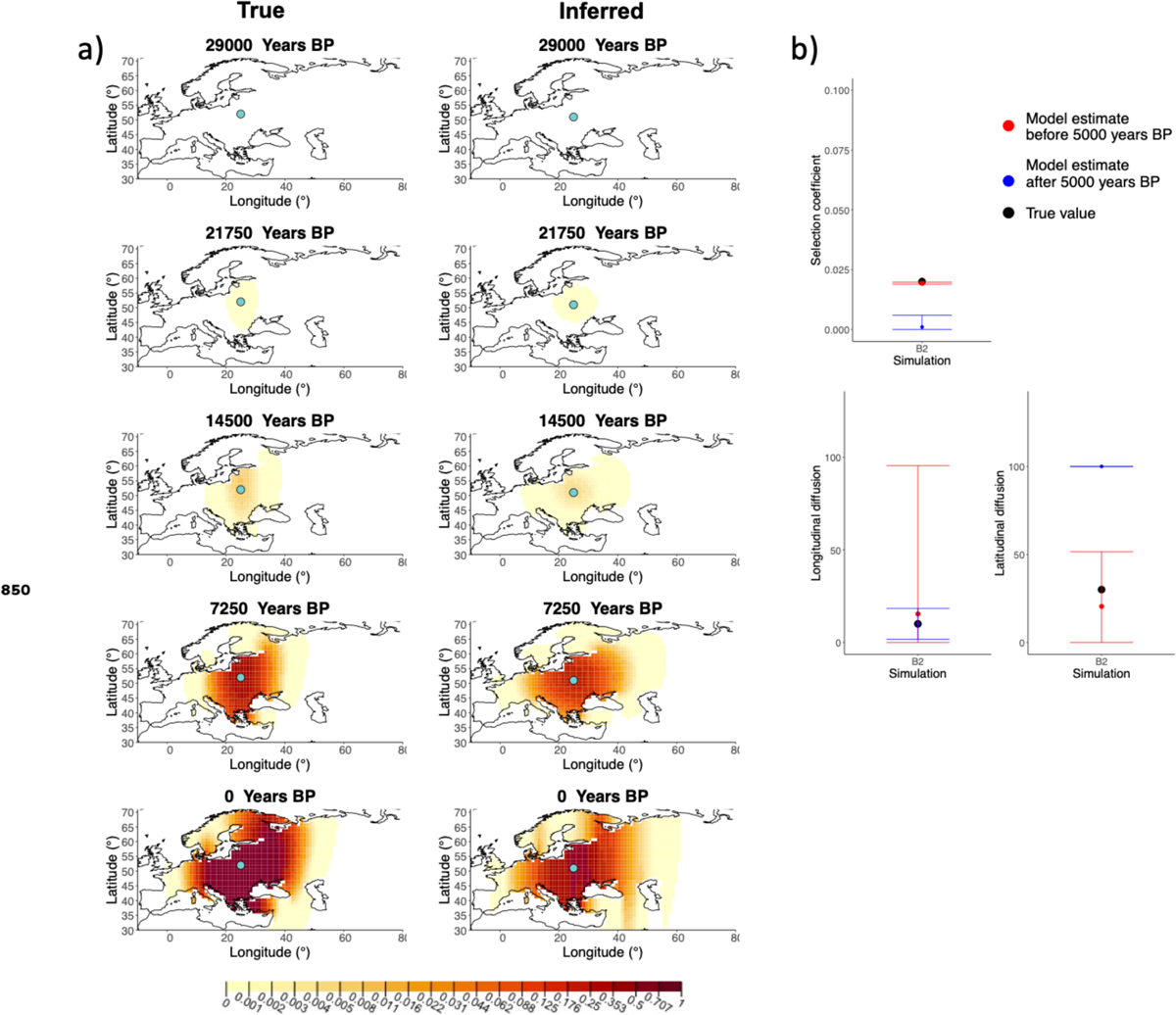
a) Comparison of true and inferred allele frequency dynamics for simulation B2. The green dot corresponds to the origin of the allele. The parameter values used to generate the frequency surface maps are summarised in **Table A1.** b) Comparison of true parameter values and model estimates. Whiskers represent 95% confidence intervals.

**Figure 1-Figure supplement 3.**
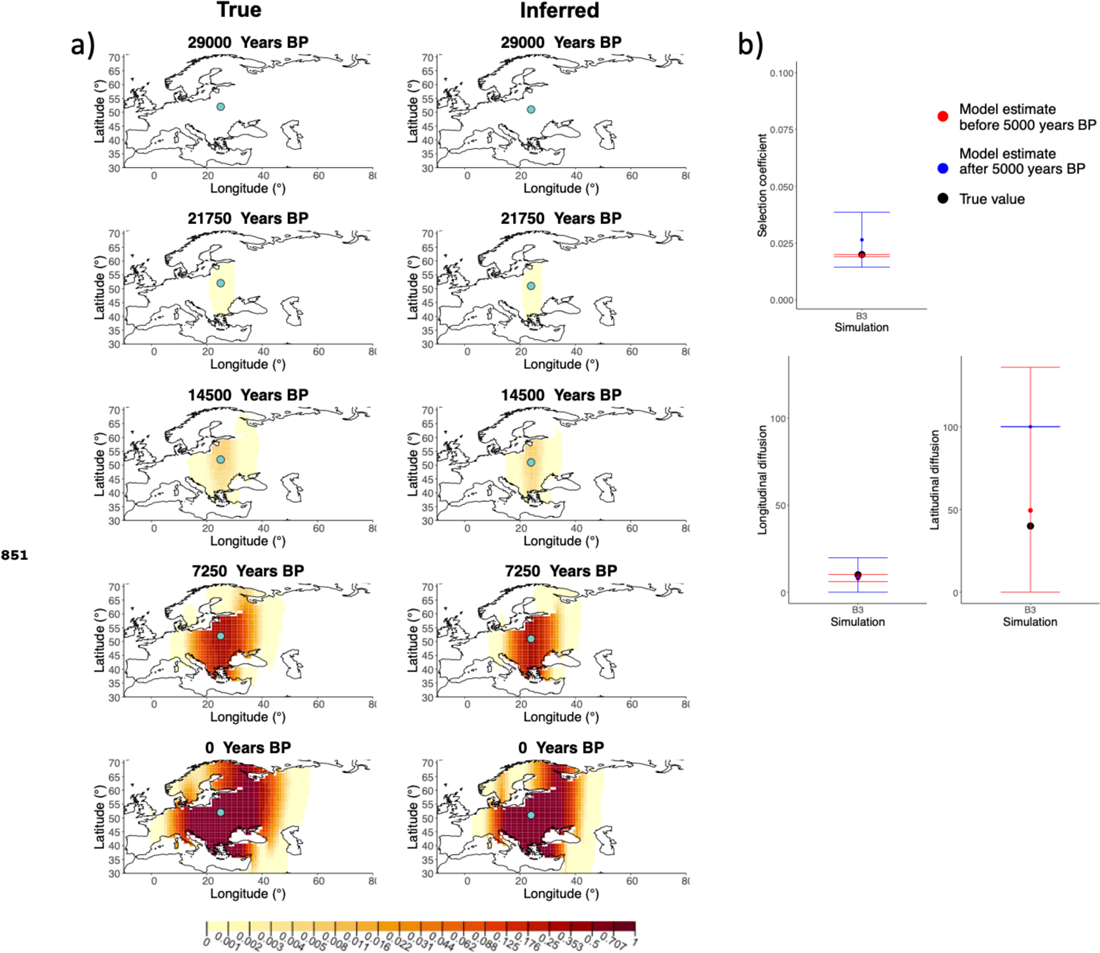
a) Comparison of true and inferred allele frequency dynamics for simulation B3. The green dot corresponds to the origin of the allele. The parameter values used to generate the frequency surface maps are summarised in **Table A1.** b) Comparison of true parameter values and model estimates. Whiskers represent 95% confidence intervals.

**Figure 1-Figure supplement 4.**
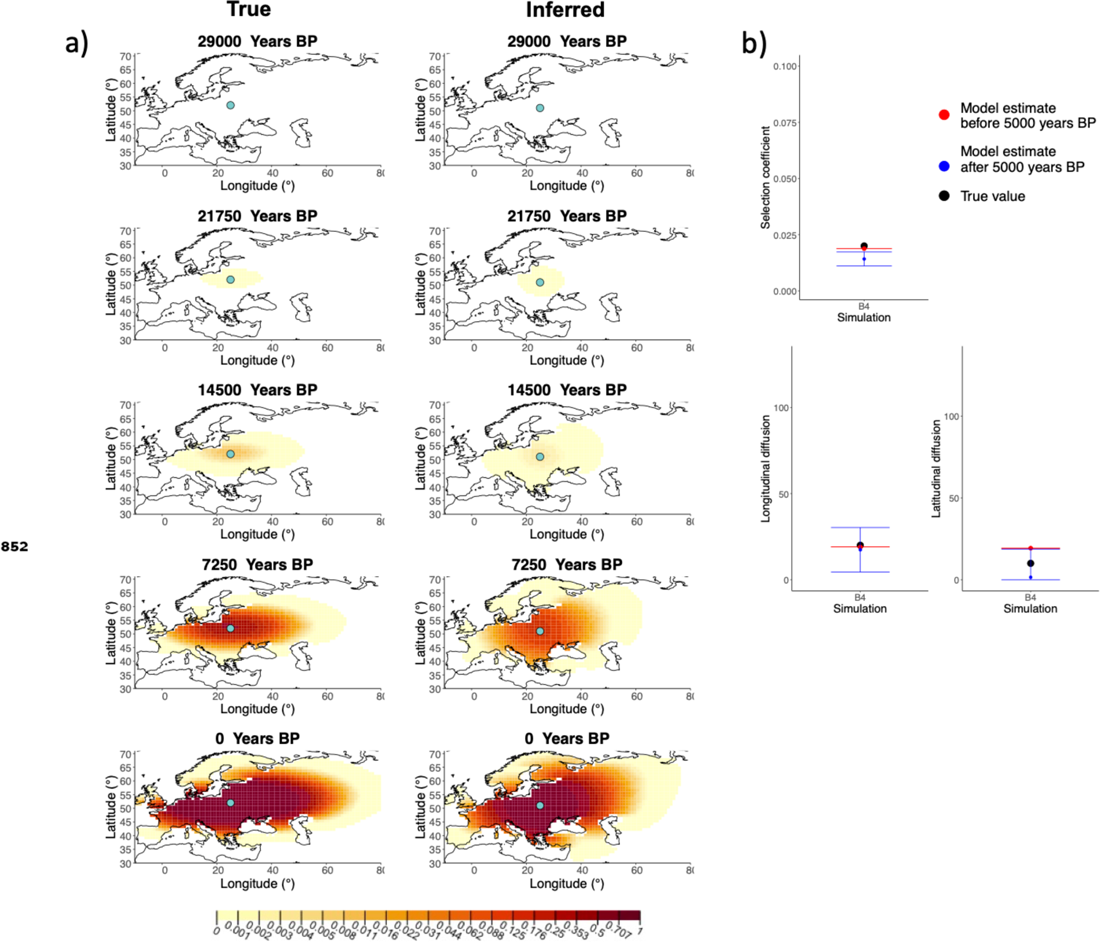
a) Comparison of true and inferred allele frequency dynamics for simulation B4. The green dot corresponds to the origin of the allele. The parameter values used to generate the frequency surface maps are summarised in **Table A1.** b) Comparison of true parameter values and model estimates. Whiskers represent 95% confidence intervals.

**Figure 1-Figure supplement 5.**
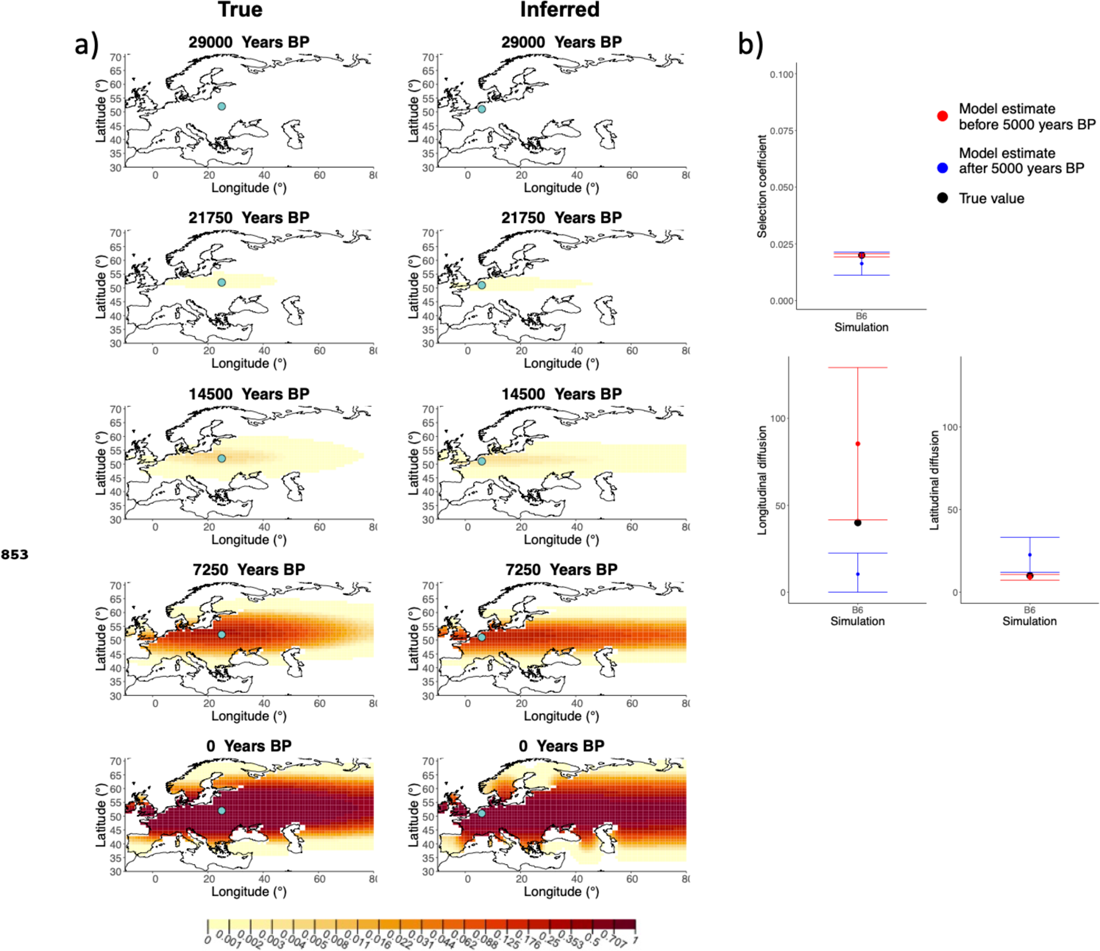
a) Comparison of true and inferred allele frequency dynamics for simulation B6. The green dot corresponds to the origin of the allele. The parameter values used to generate the frequency surface maps are summarised in **Table A1.** b) Comparison of true parameter values and model estimates. Whiskers represent 95% confidence intervals.

**Figure 1-Figure supplement 6.**
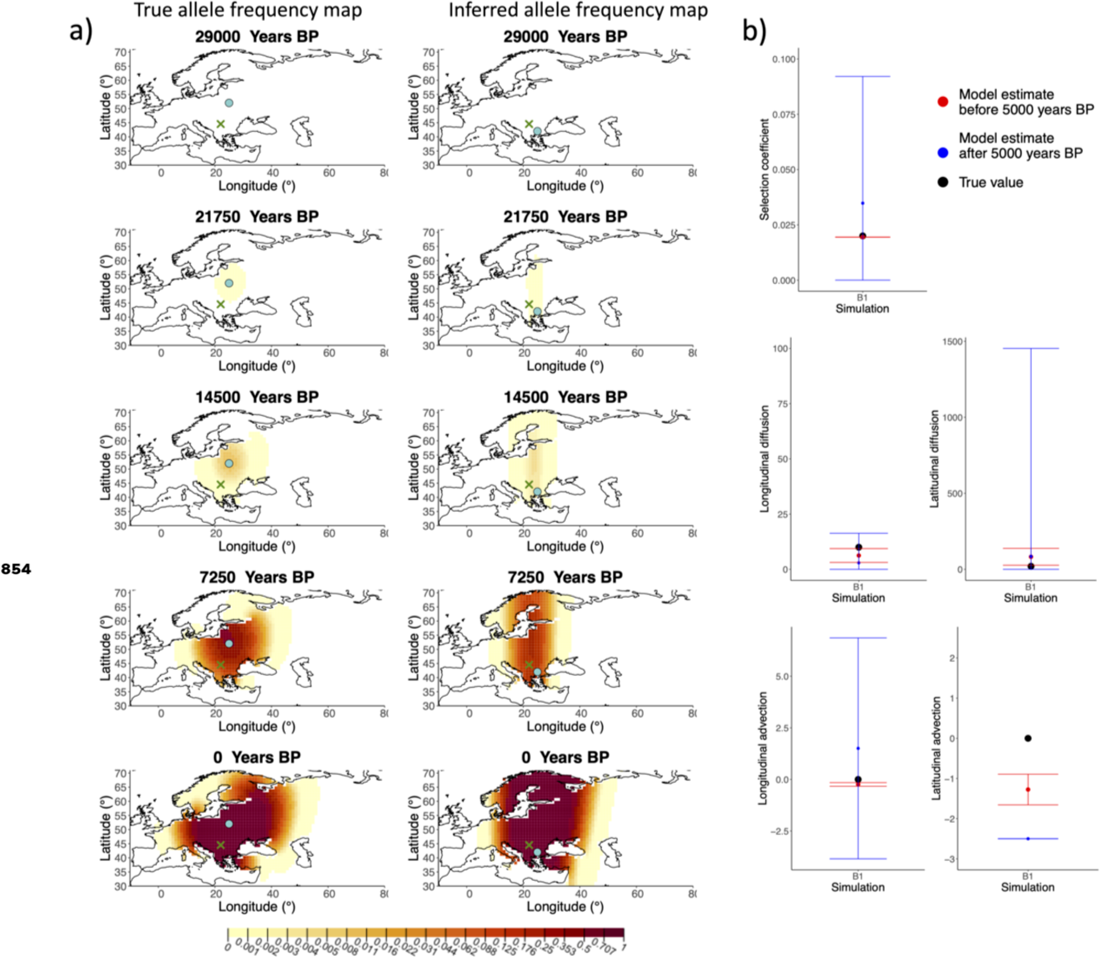
a) Comparison of true allele frequency dynamics for simulation B1 and those inferred by the model C. The green dot shows the origin of the derived allele and the cross represents the location of the first individual that carried it. b) Comparison of true parameter values and model estimates. Whiskers represent 95% confidence intervals.

**Figure 1-Figure supplement 7.**
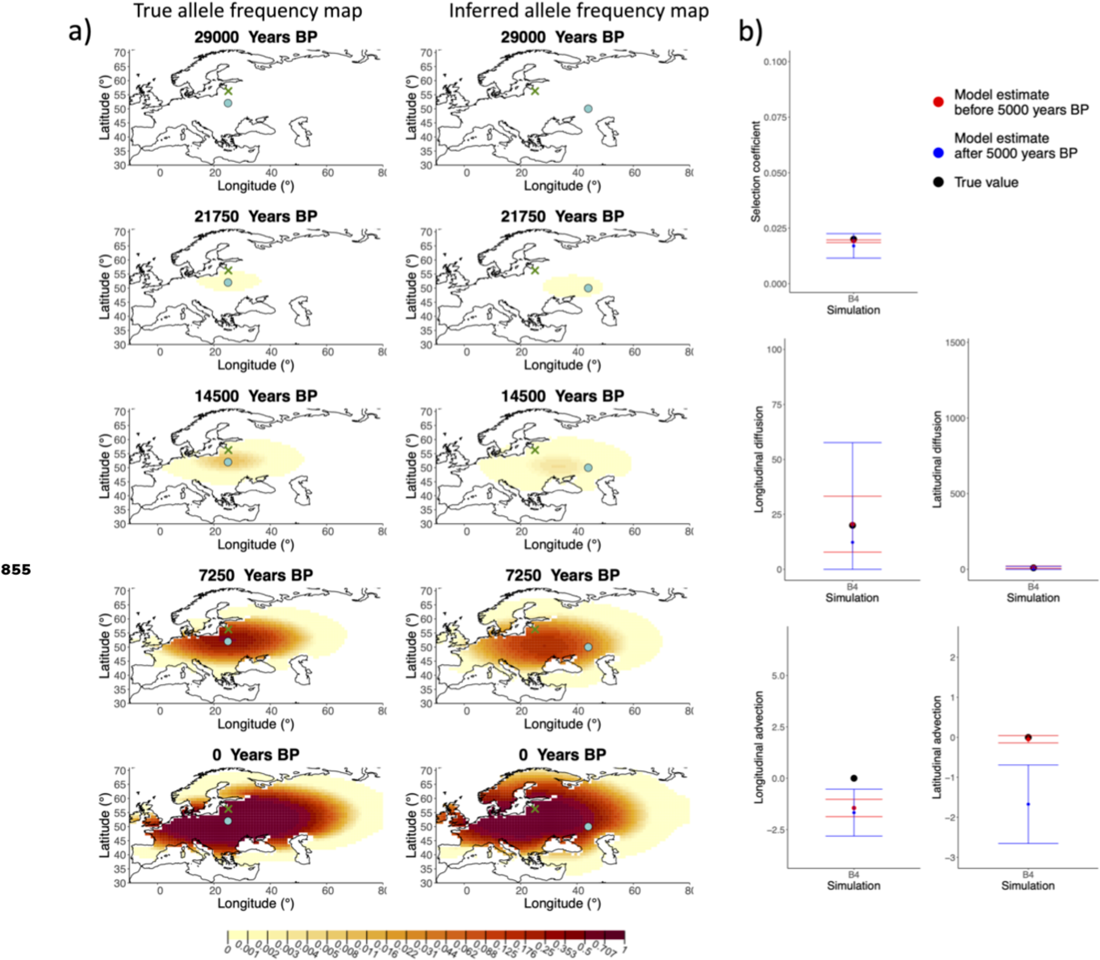
a) Comparison of true allele frequency dynamics for simulation B4 and those inferred by the model C. The green dot corresponds to the origin of the allele and the cross represents the first sample having the derived variant. b) Comparison of true parameter values and model estimates. Whiskers represent 95% confidence intervals.

**Figure 2-Figure supplement 1.**
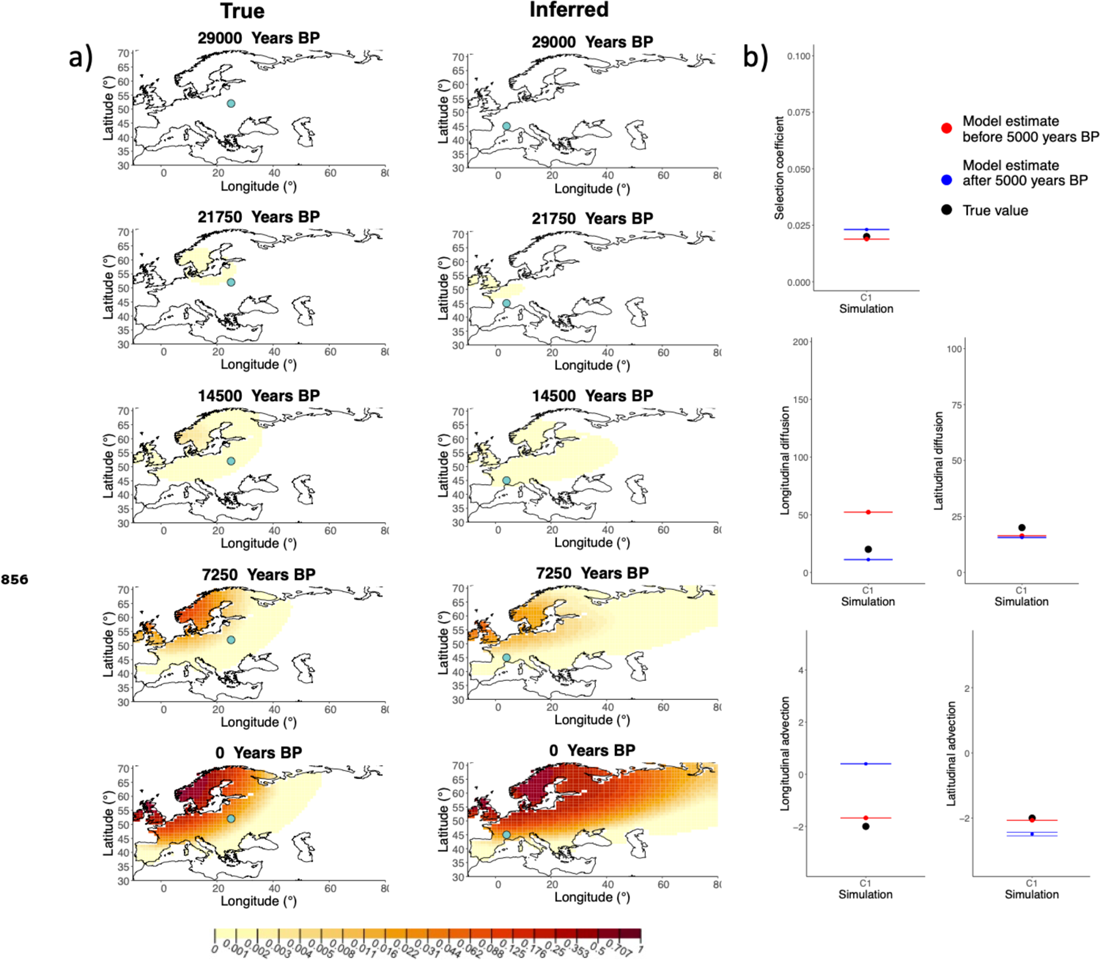
a) Comparison of true and inferred allele frequency dynamics for one of the simulations including advection (C1). The green dot corresponds to the origin of the allele. The parameter values used to generate the frequency surface maps are summarised in **Table A2.** b) Comparison of true parameter values and model estimates. Whiskers represent 95% confidence intervals.

**Figure 2-Figure supplement 2.**
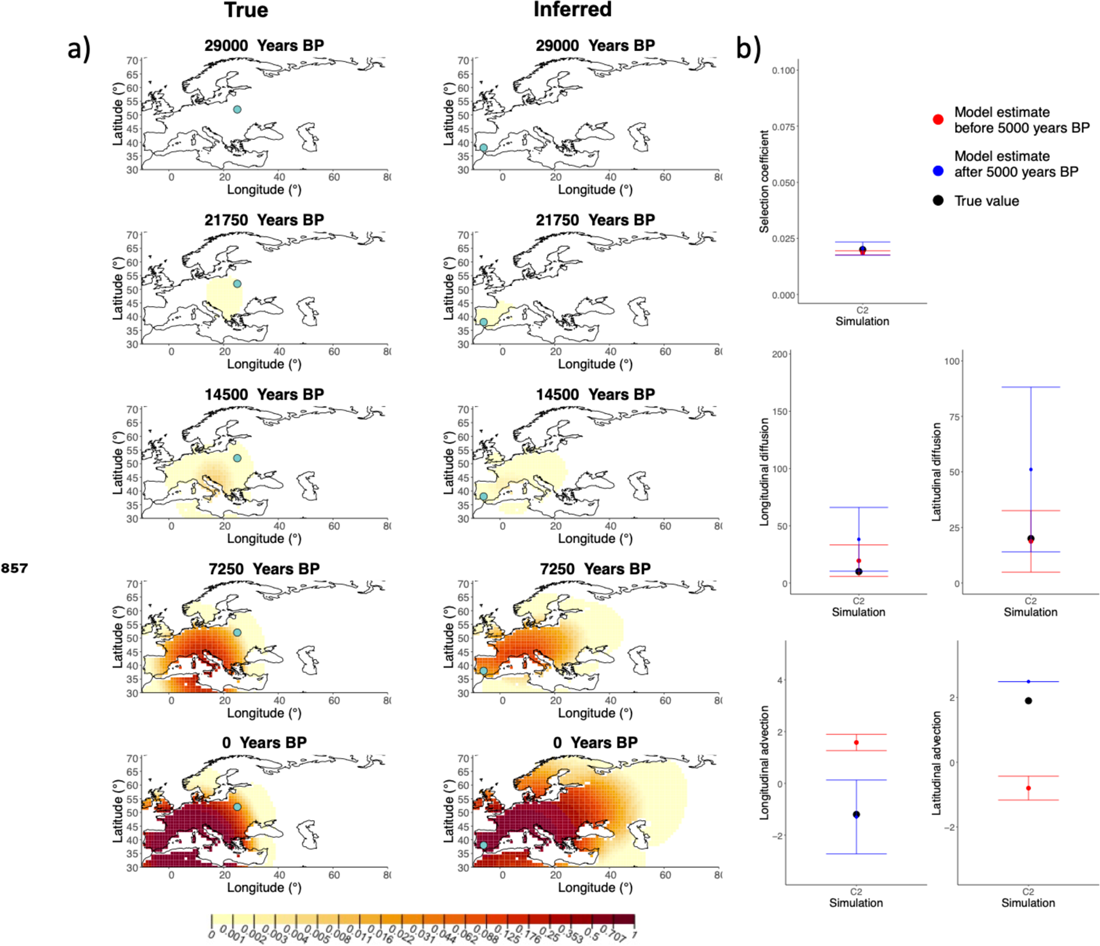
a) Comparison of true and inferred allele frequency dynamics for one of the simulations including advection (C2). The green dot corresponds to the origin of the allele. The parameter values used to generate the frequency surface maps are summarised in **Table A2.** b) Comparison of true parameter values and model estimates. Whiskers represent 95% confidence intervals.

**Figure 2-Figure supplement 3.**
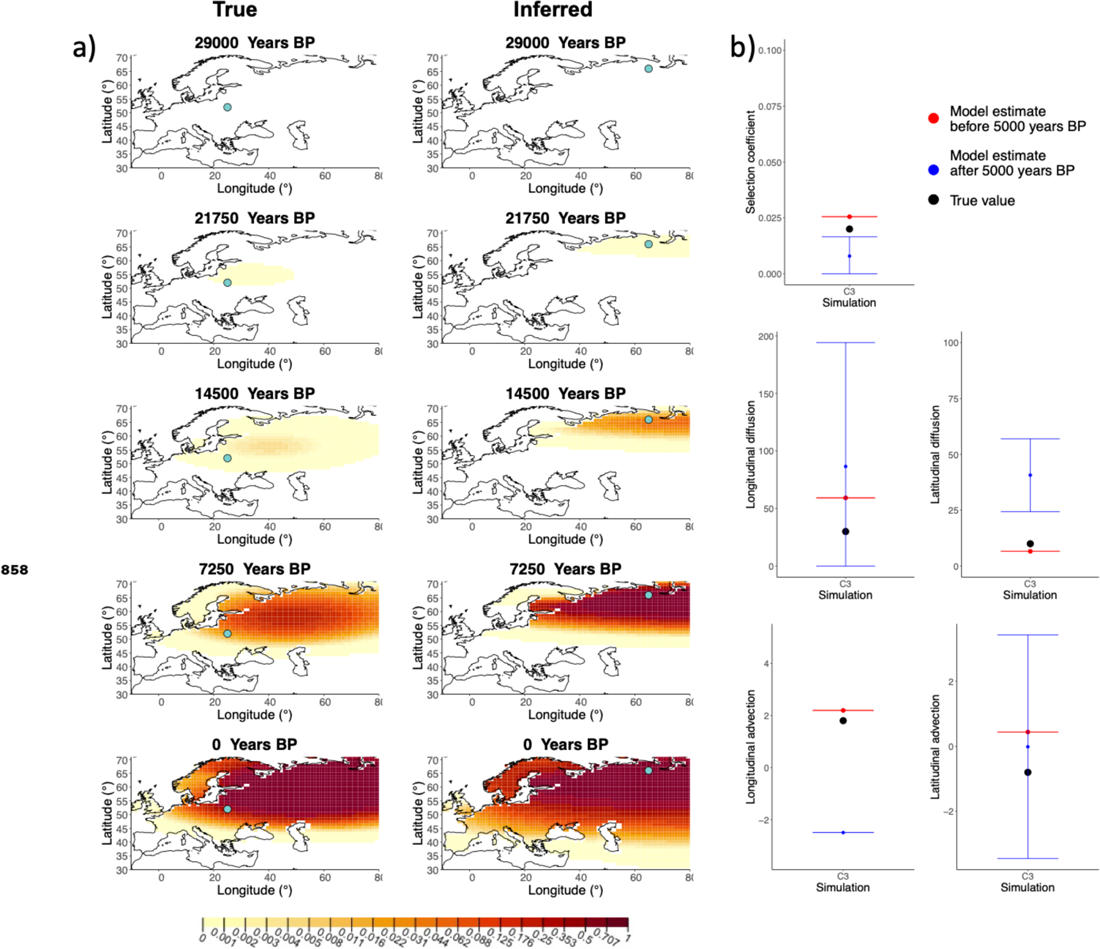
a) Comparison of true and inferred allele frequency dynamics for one of the simulations including advection (C3). The green dot corresponds to the origin of the allele. The parameter values used to generate the frequency surface maps are summarised in **Table A2.** b) Comparison of true parameter values and model estimates. Whiskers represent 95% confidence intervals.

**Figure 3-Figure supplement 1.**
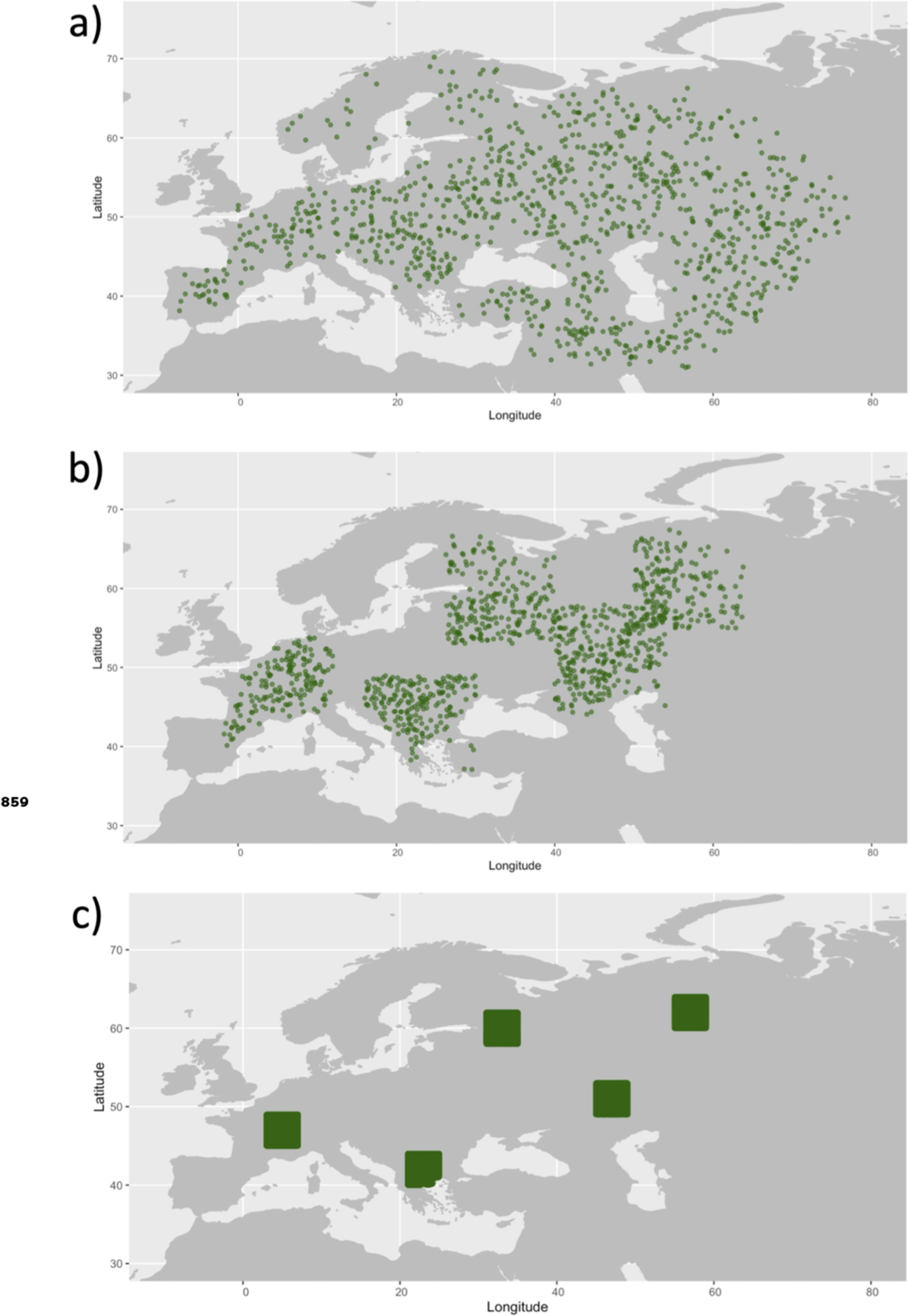
Examples of spatial sampling scenarios for each of the three clustering schemes. We chose five locations and increasingly restricted the area where we allowed the individuals to be sampled. a) Map showing homogeneous sampling scheme in which we did not impose any spatial restrictions of individuals sampled. b) Intermediate sampling scheme with the region restricted to 7 degrees in each cardinal direction from each of the chosen locations c) Extreme sampling scheme with the sampling region restricted to 2 degrees in each cardinal direction from the chosen locations.

**Figure 3-Figure supplement 2.**
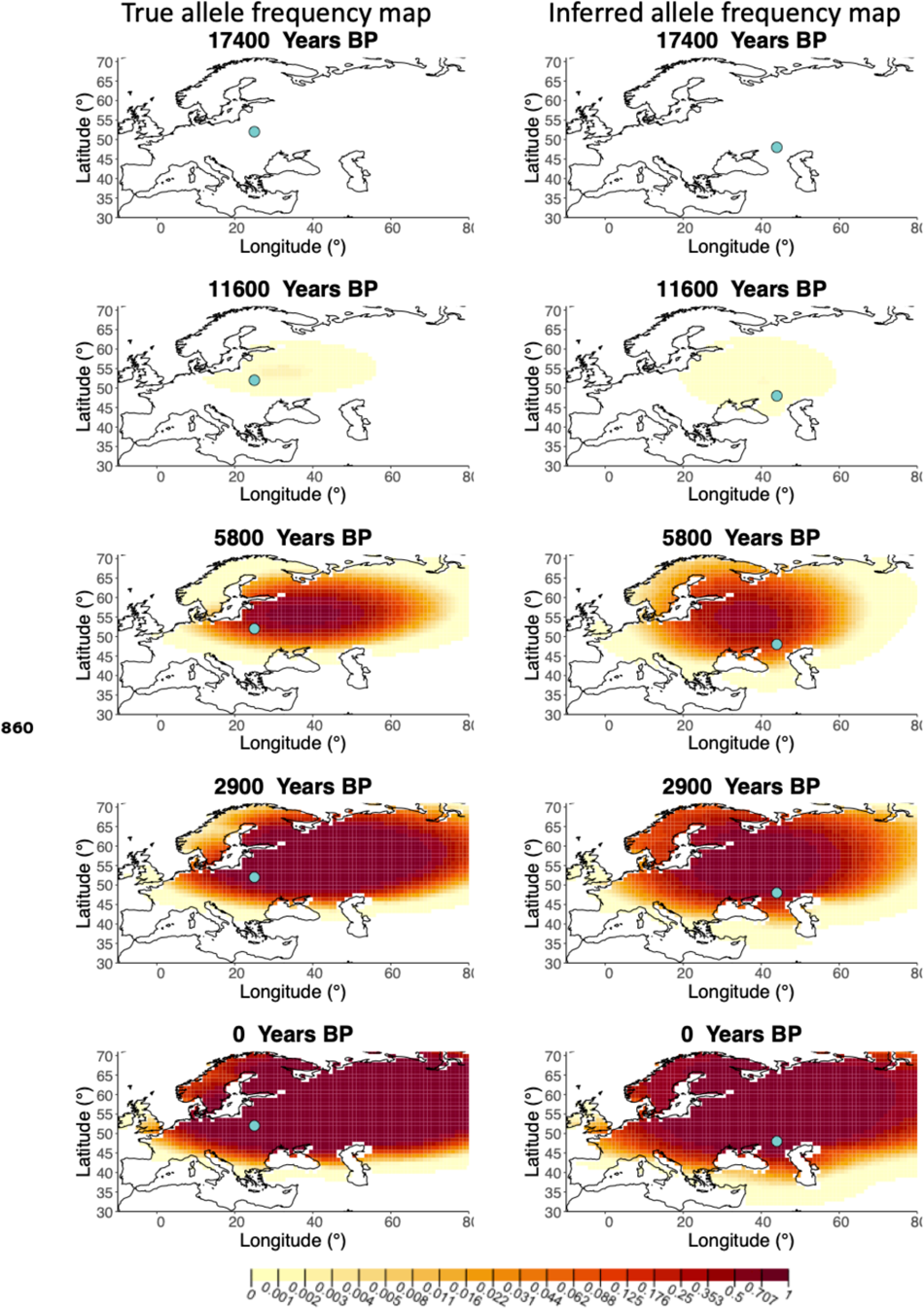
Left-Allele frequency map generated using true parameter values. Right - Allele frequency map generated using parameter estimates for “homogeneous 75%/25%” clustering scheme. Parameter values used to generate the maps are summarised in **Table A3.**

**Figure 3-Figure supplement 3.**
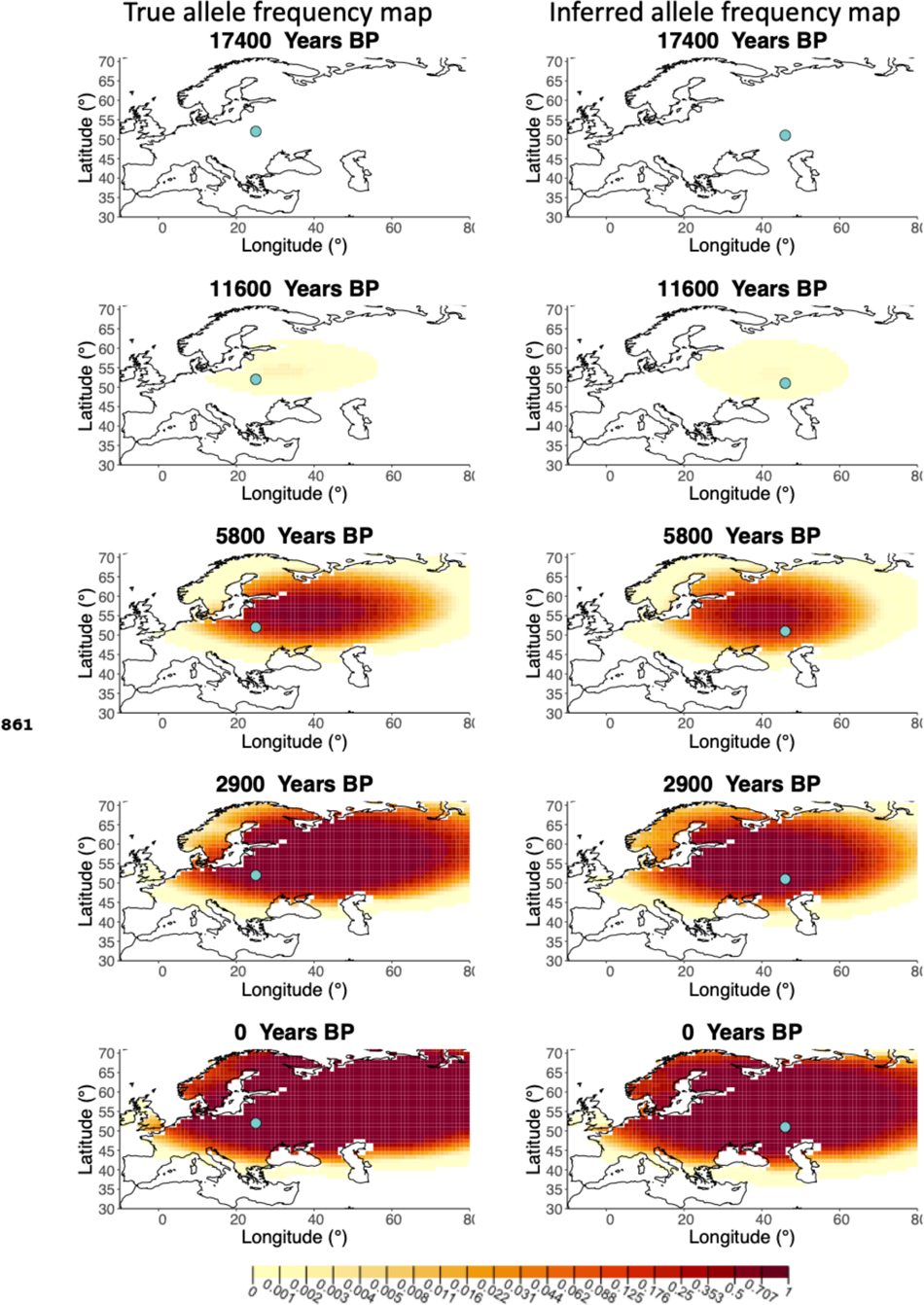
Left-Allele frequency map generated using true parameter values. Right - Allele frequency map generated using parameter estimates for “homogeneous 50%/50%” clustering scheme. Parameter values used to generate the maps are summarised in **Table A3.**

**Figure 3-Figure supplement 4.**
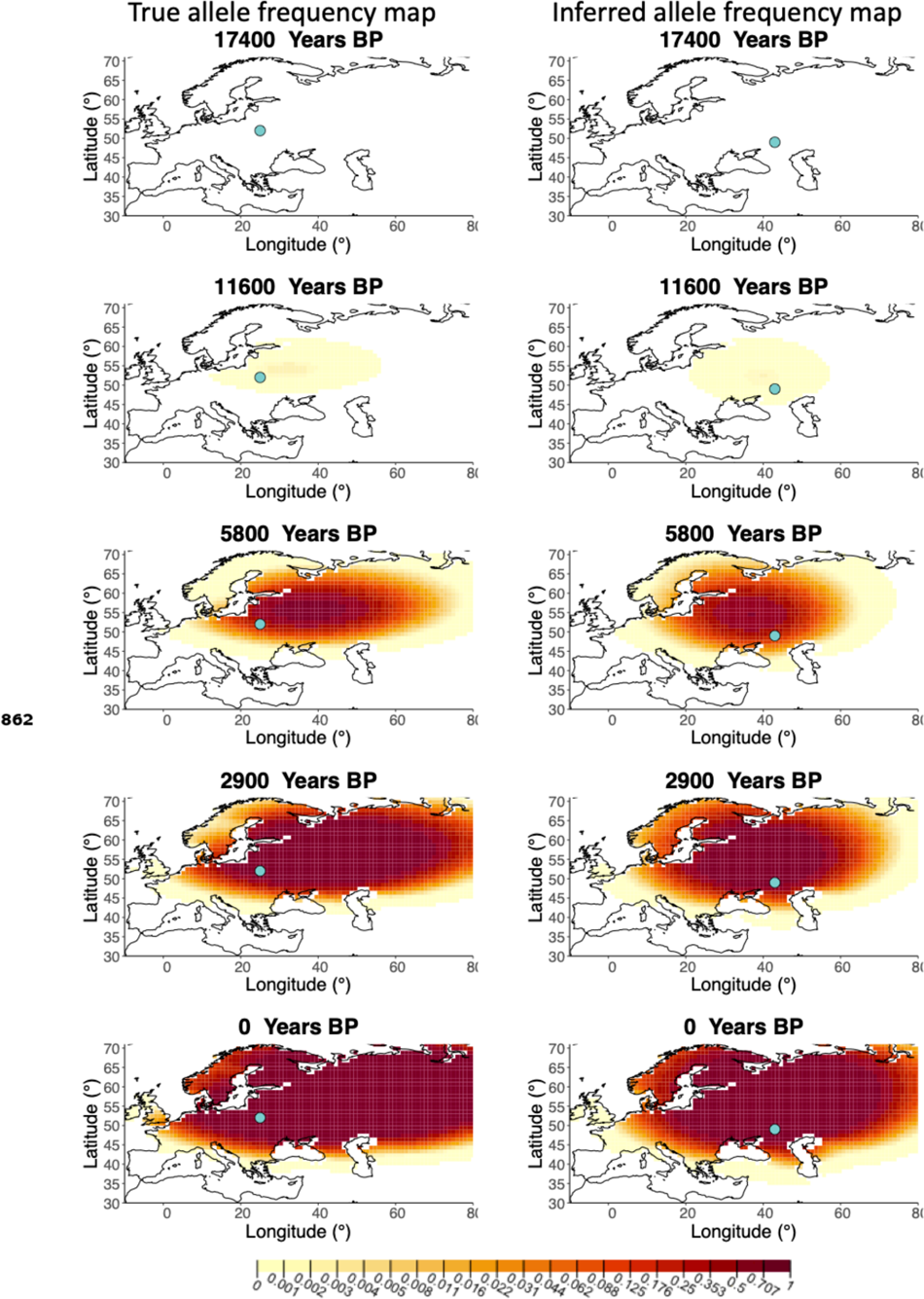
Left-Allele frequency map generated using true parameter values. Right - Allele frequency map generated using parameter estimates for “homogeneous 25%/75%” clustering scheme. Parameter values used to generate the maps are summarised in **Table A3.**

**Figure 3-Figure supplement 5.**
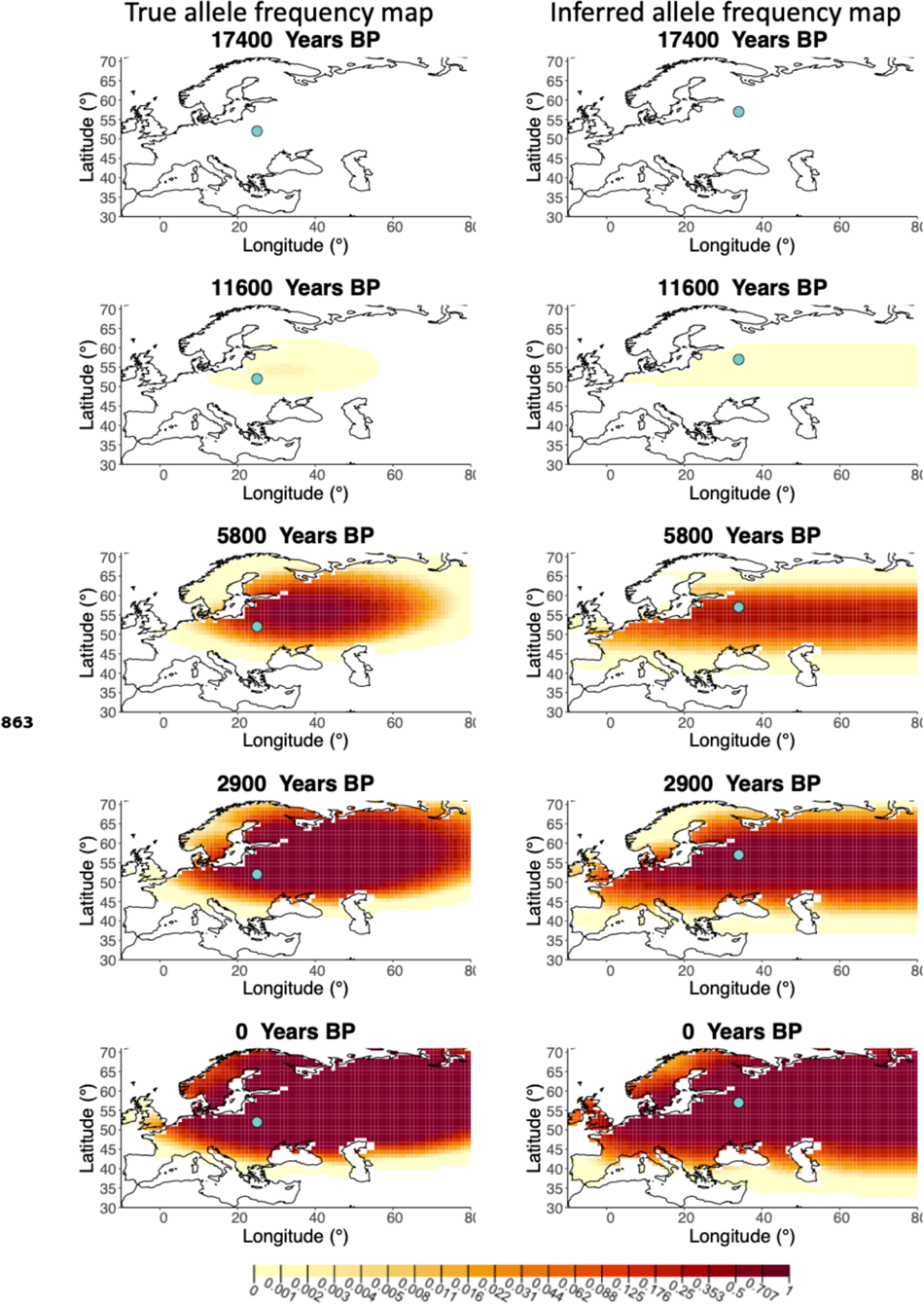
Left - Allele frequency map generated using true parameter values. Right - Allele frequency map generated using parameter estimates for “intermediate 50%/50%” clustering scheme. Parameter values used to generate the maps are summarised in **Table A3.**

**Figure 3-Figure supplement 6.**
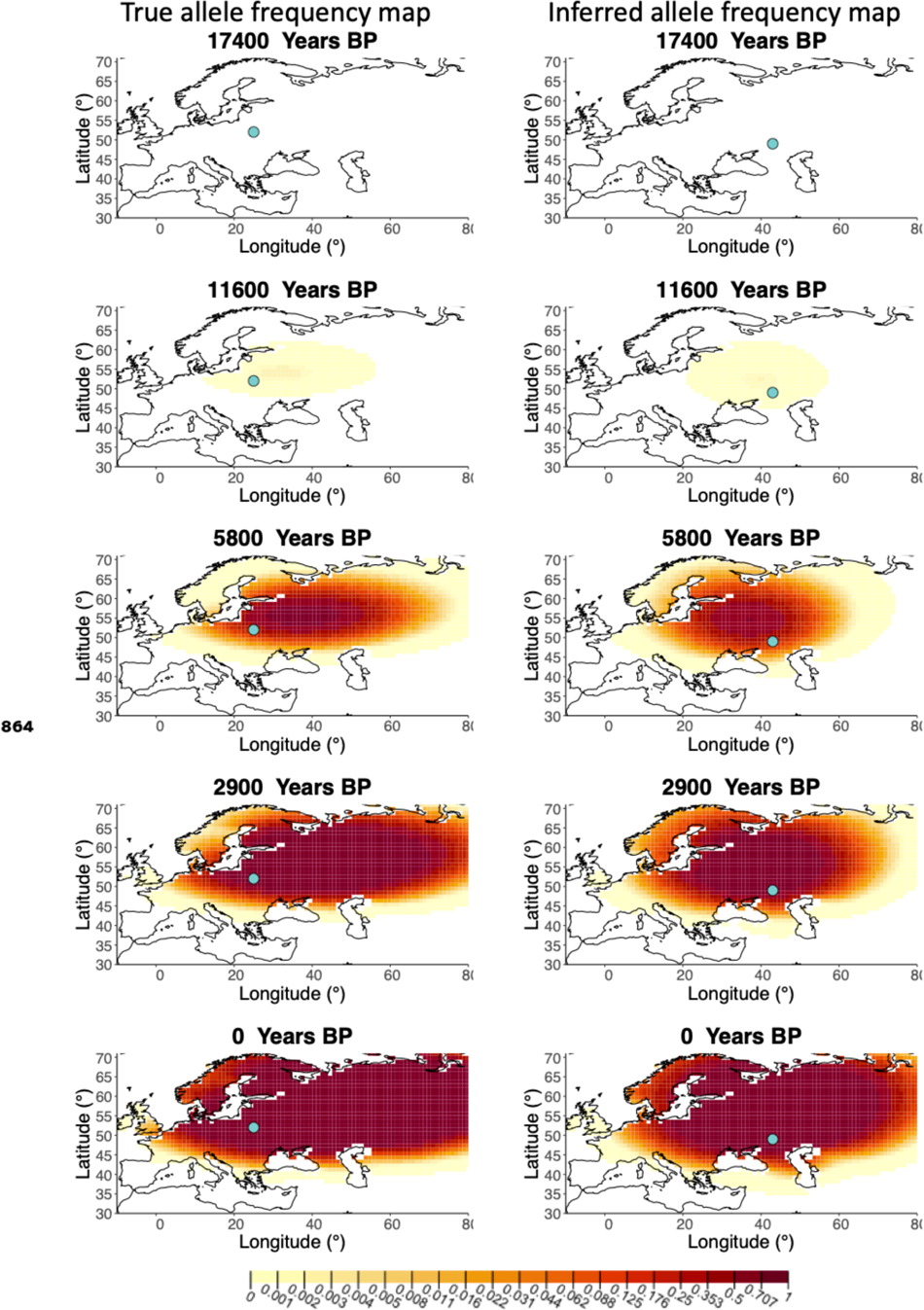
Left - Allele frequency map generated using true parameter values. Right - Allele frequency map generated using parameter estimates for “intermediate 25%/75%” clustering scheme. Parameter values used to generate the maps are summarised in **Table A3.**

**Figure 3-Figure supplement 7.**
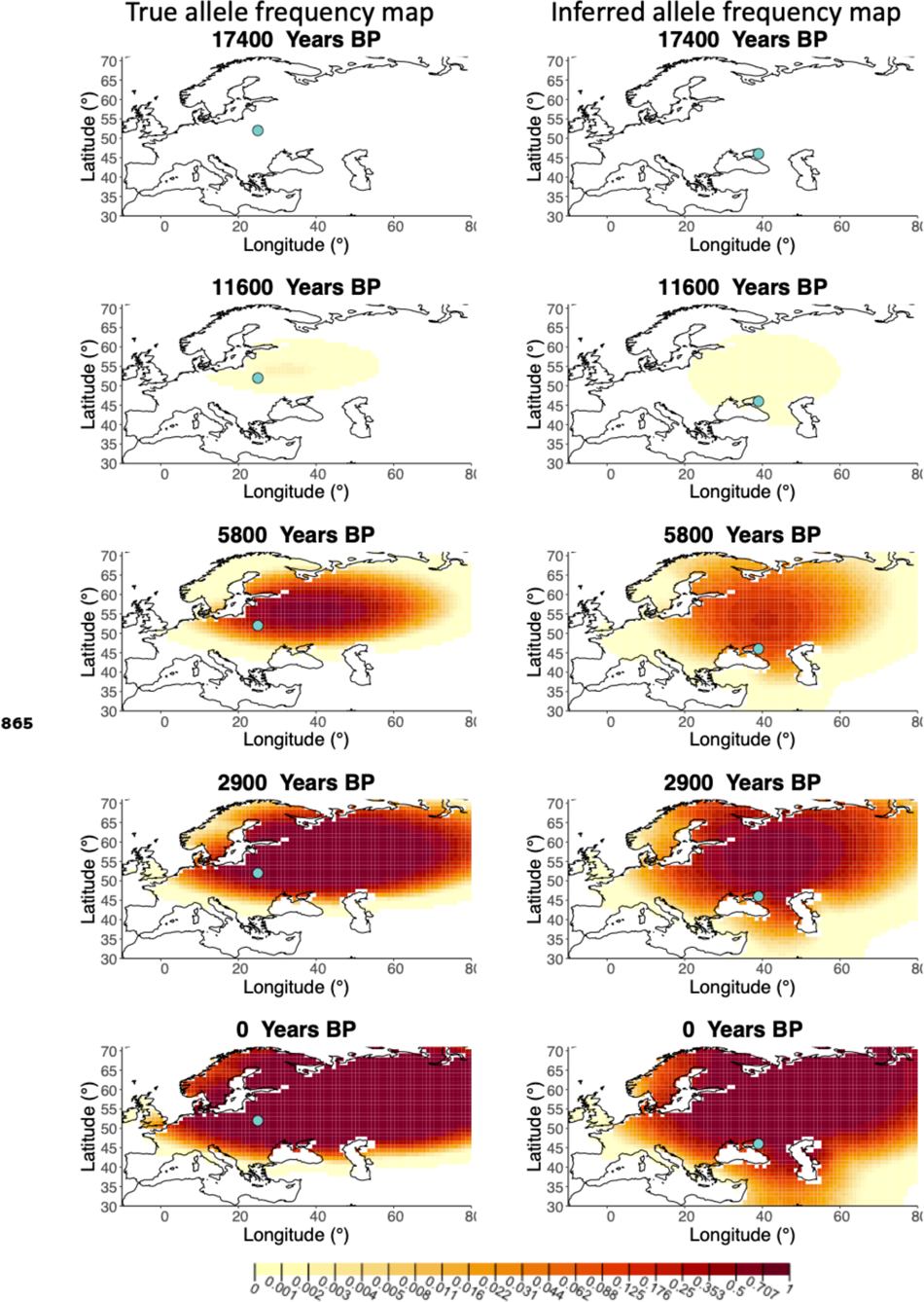
Left - Allele frequency map generated using true parameter values. Right - Allele frequency map generated using parameter estimates for “extreme 75%/25%” clustering scheme. Parameter values used to generate the maps are summarised in **Table A3.**

**Figure 3-Figure supplement 8.**
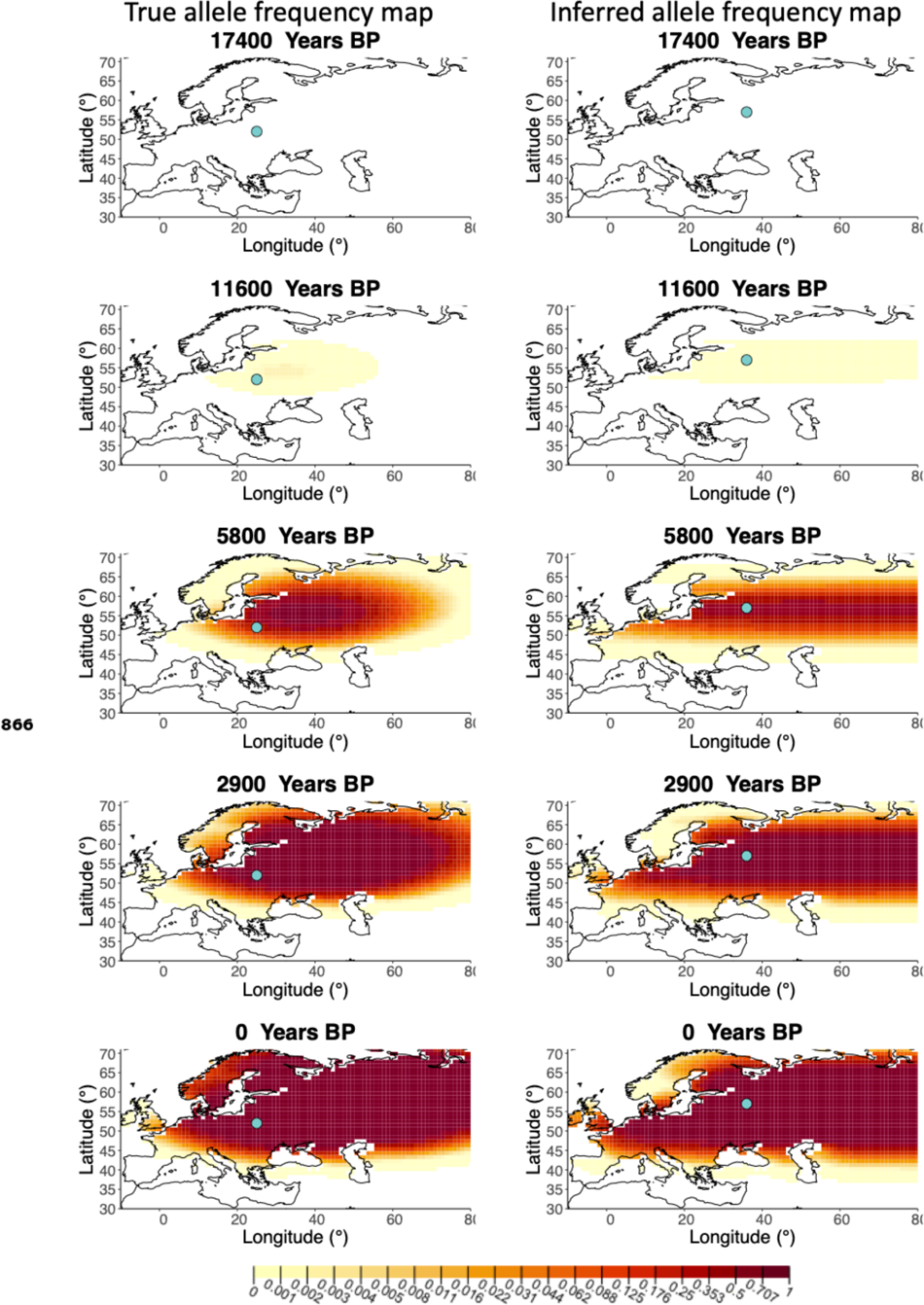
Left - Allele frequency map generated using true parameter values. Right - Allele frequency map generated using parameter estimates for “extreme 50%/50%” clustering scheme. Parameter values used to generate the maps are summarised in **Table A3.**

**Figure 3-Figure supplement 9.**
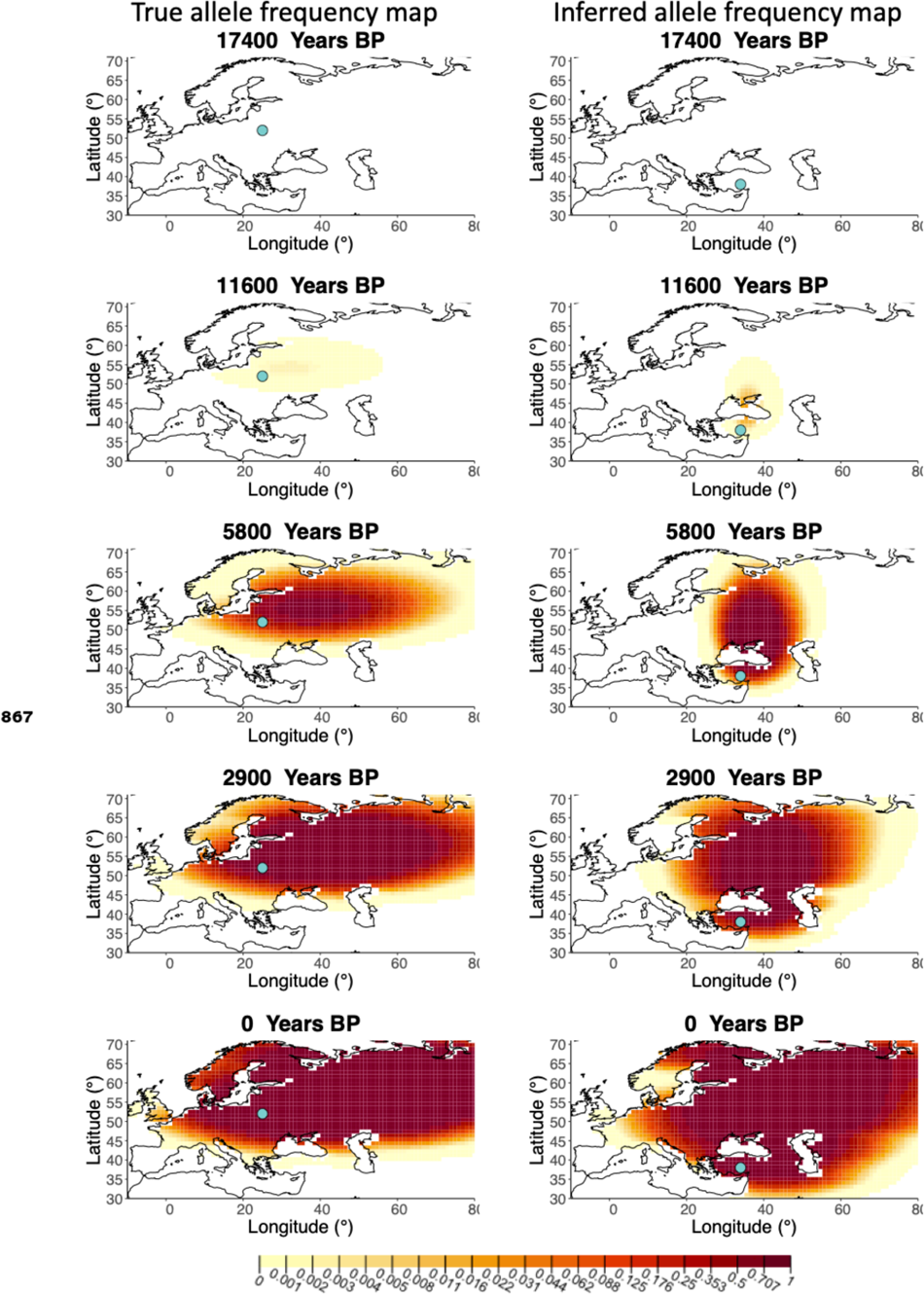
Left - Allele frequency map generated using true parameter values. Right - Allele frequency map generated using parameter estimates for “extreme 25%/75%” clustering scheme. Parameter values used to generate the maps are summarised in **Table A3.**

**Figure 4-Figure supplement 1.**
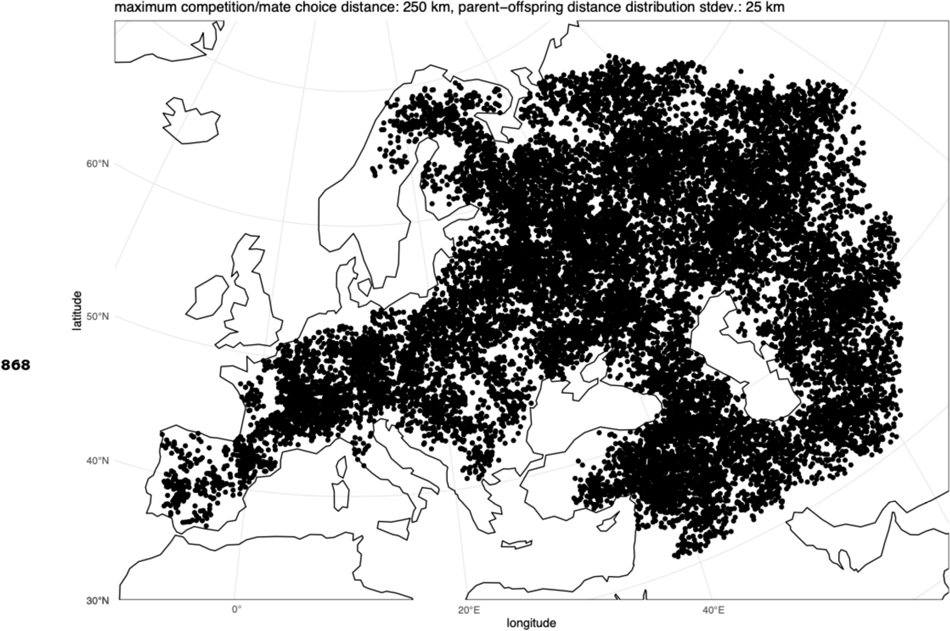
Distribution of individuals across the map under neutrality, showing the tendency of individuals to cluster together.

**Figure 6-Figure supplement 1.**
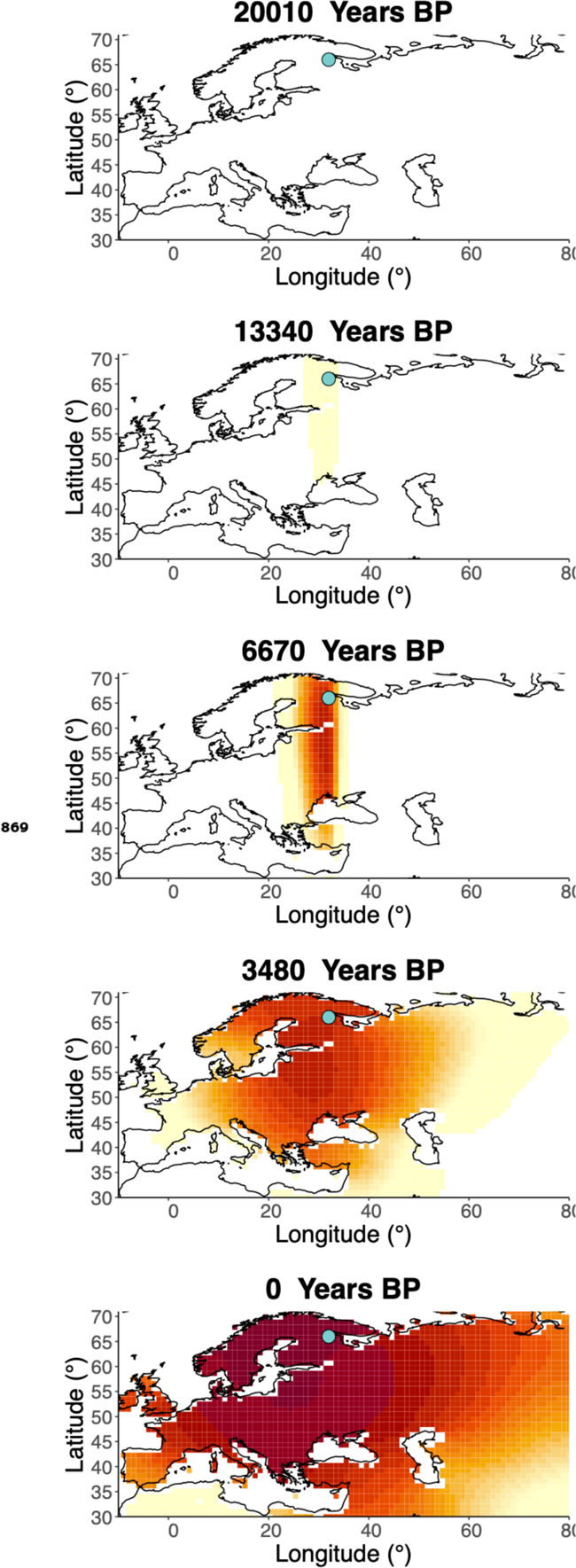
Inferred frequency dynamics of rs4988235(T) using the allele age that was inferred in ***Albers and Mcvean* (2020).**

**Figure 6-Figure supplement 2.**
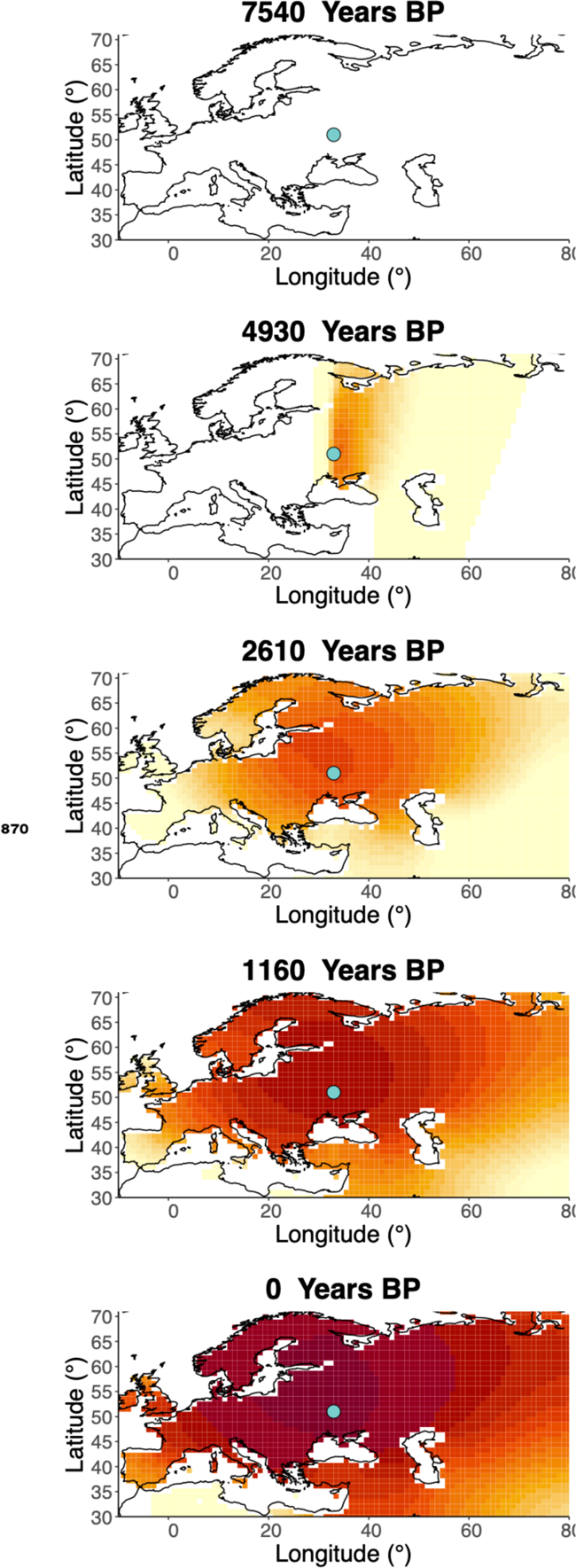
Inferred frequency dynamics of rs4988235(T) when the origin of the allele is moved 10 degrees west from the original estimate.

**Figure 6-Figure supplement 3.**
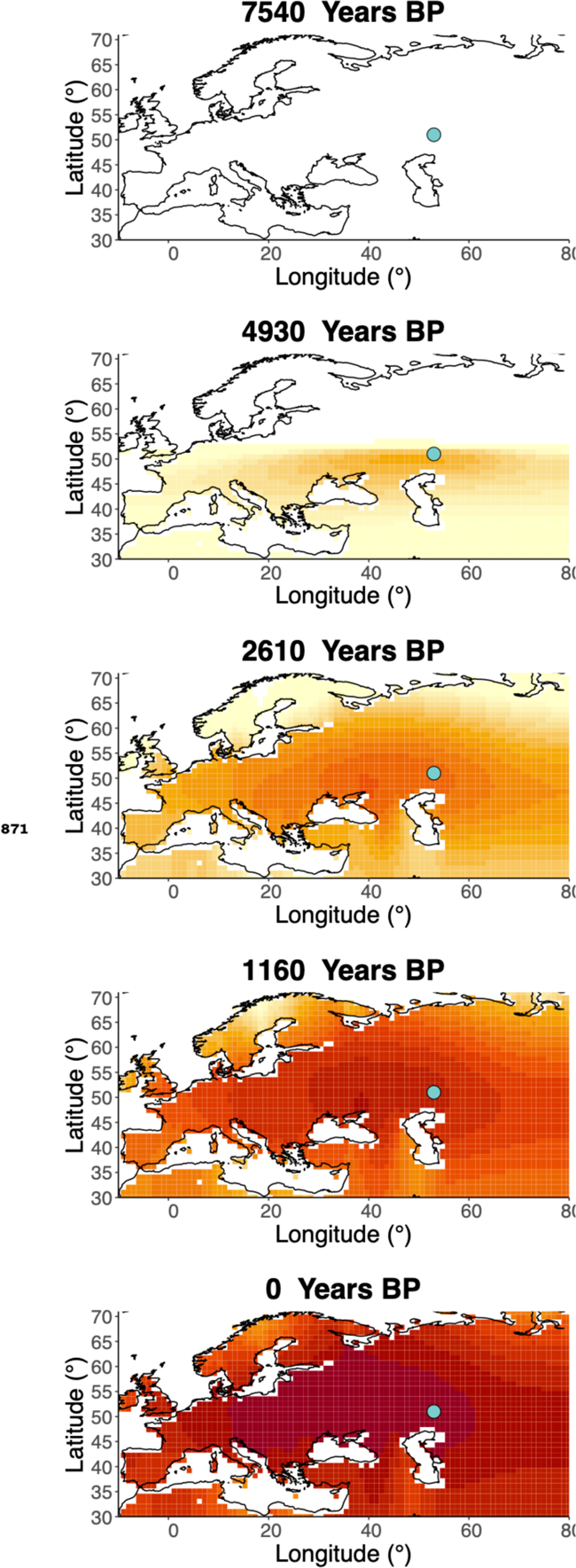
Inferred frequency dynamics of rs4988235(T) when the origin of the allele is moved 10 degrees east from the original estimate.

**Figure 6-Figure supplement 4.**
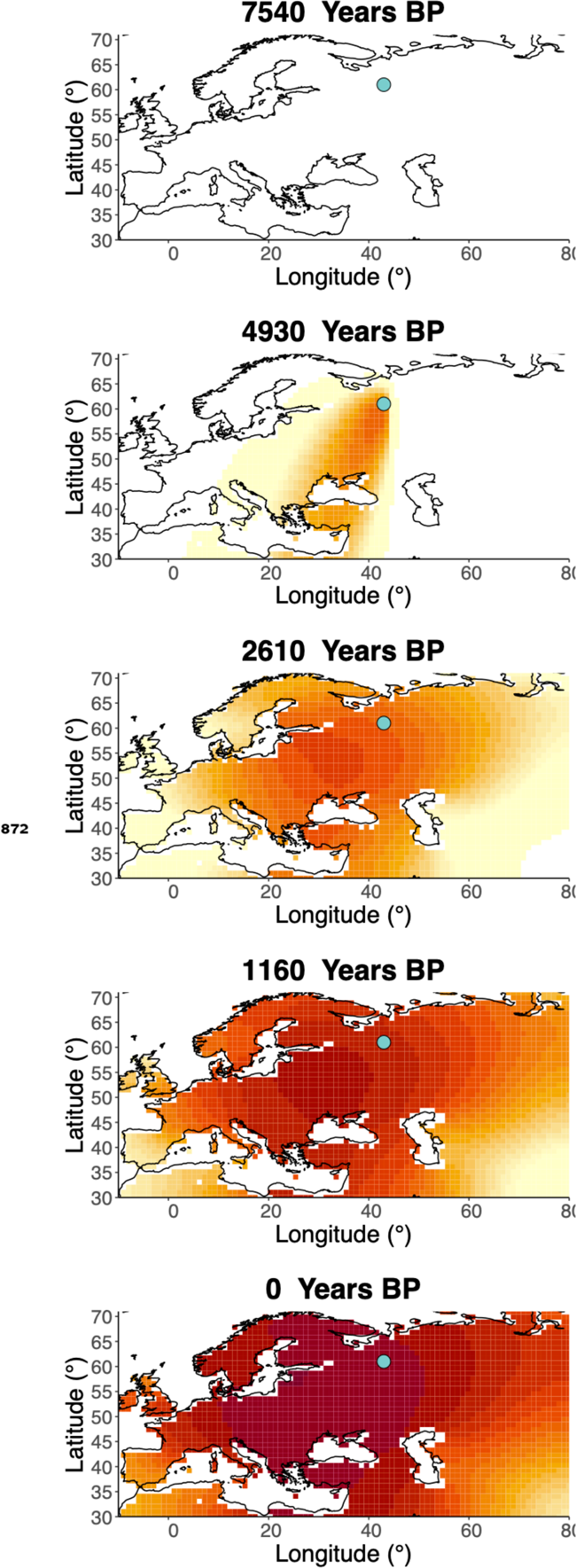
Inferred frequency dynamics of rs4988235(T) when the origin of the allele is moved 10 degrees north from the original estimate.

**Figure 6-Figure supplement 5.**
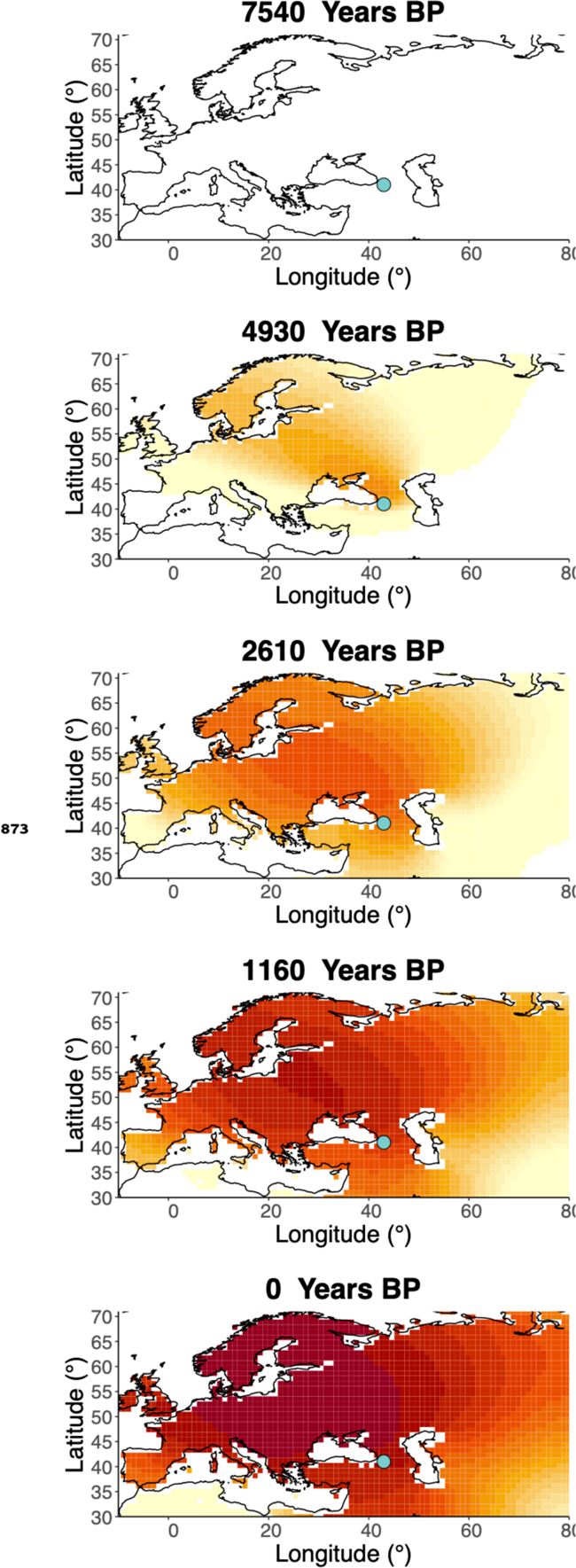
Inferred frequency dynamics of rs4988235(T) when the origin of the allele is moved 10 degrees south from the original estimate.

**Figure 6-Figure supplement 6.**
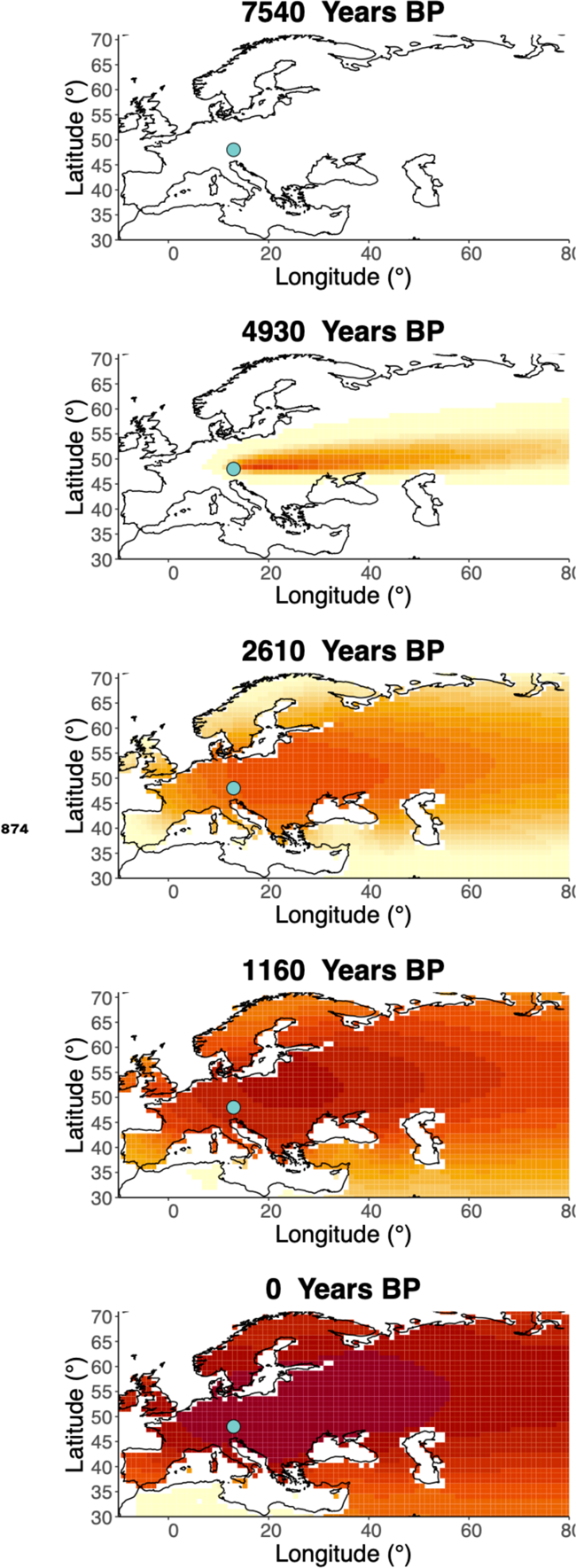
Inferred frequency dynamics of rs4988235(D forcing the geographic origin of the allele to be at the location inferred in ***/tan et al.* (2009).**

**Figure 6-Figure supplement 7.**
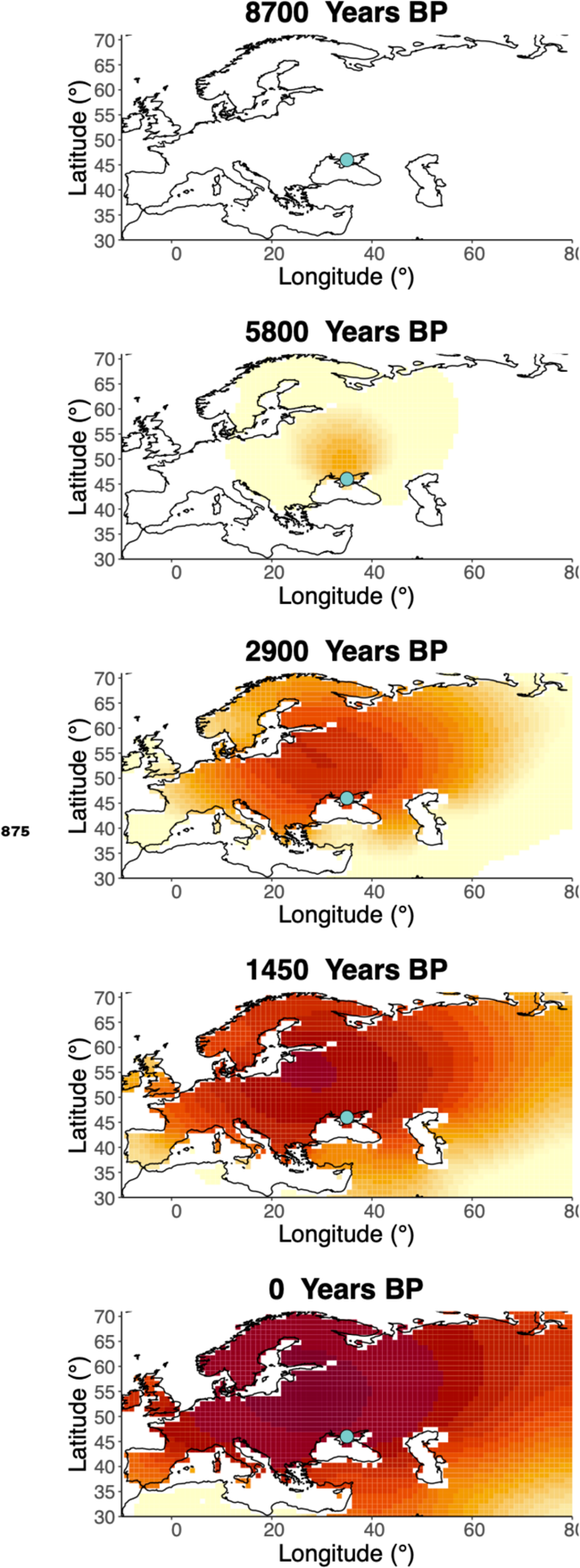
Inferred frequency dynamics of rs4988235(T) assuming the allele age to be the lower end of the 95%credible interval for the allele age inferred in ***/tan et al.* (2009).**

**Figure 6-Figure supplement 8.**
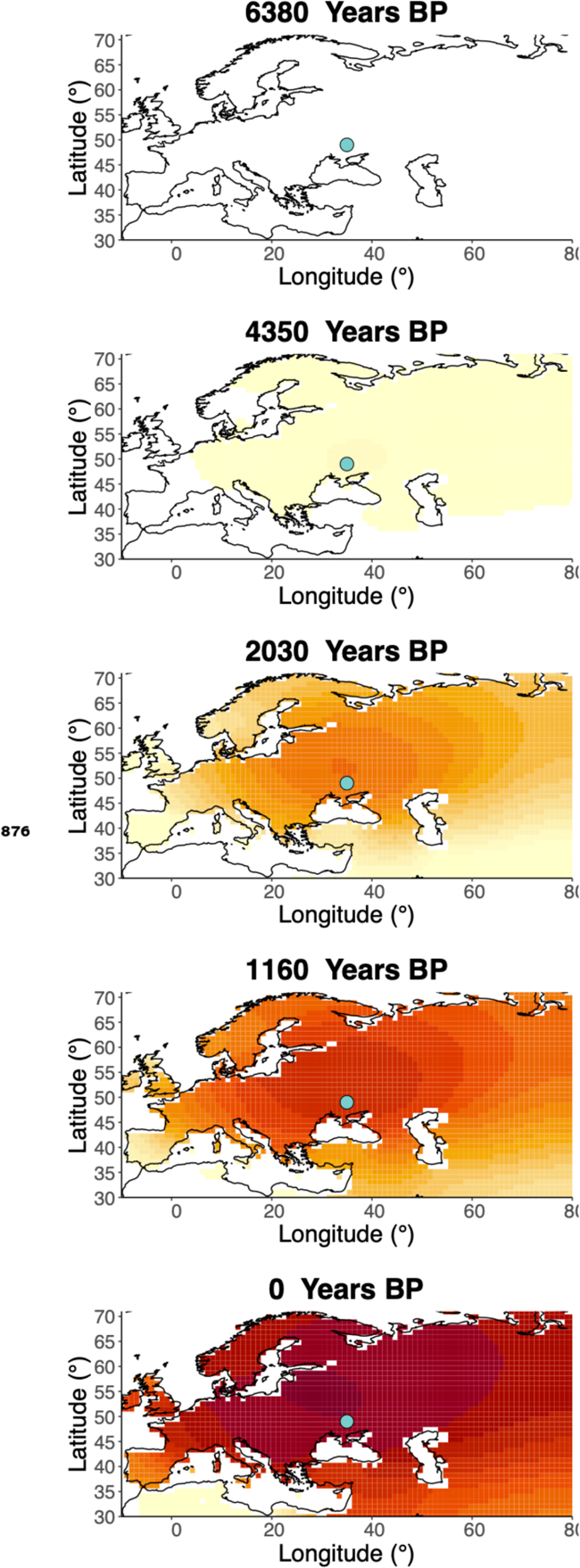
Inferred frequency dynamics of rs4988235(T) assuming the allele age to be the higher end of the 95%credible interval for the allele age inferred in ***/tan et al.* (2009).**

**Figure 6-Figure supplement 9.**
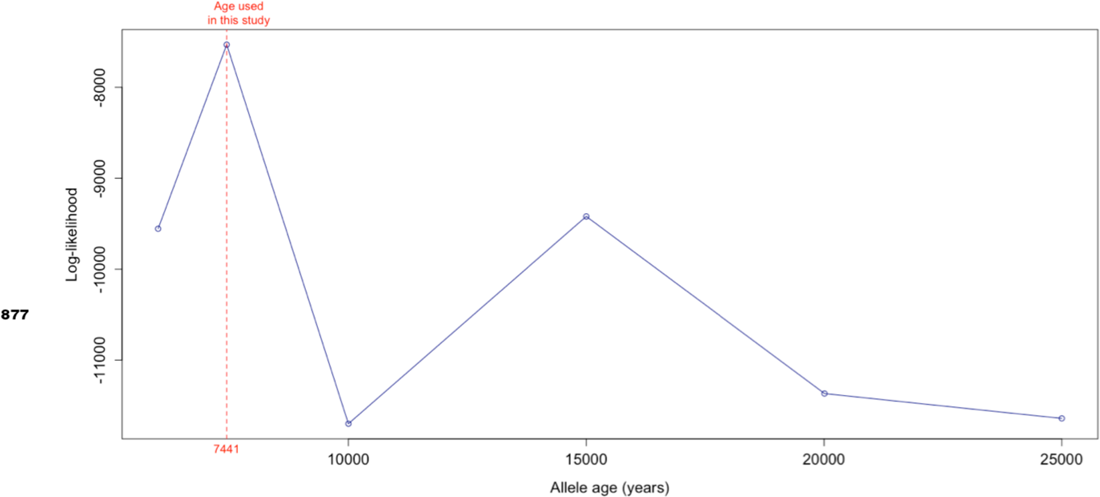
Log-likelihood values for model runs using different ages of the rs4988235(T) allele as input, with the age inferred by ***/tan et al.* (2009)** we use as fixed input highlighted in red.

**Figure 8-Figure supplement 1.**
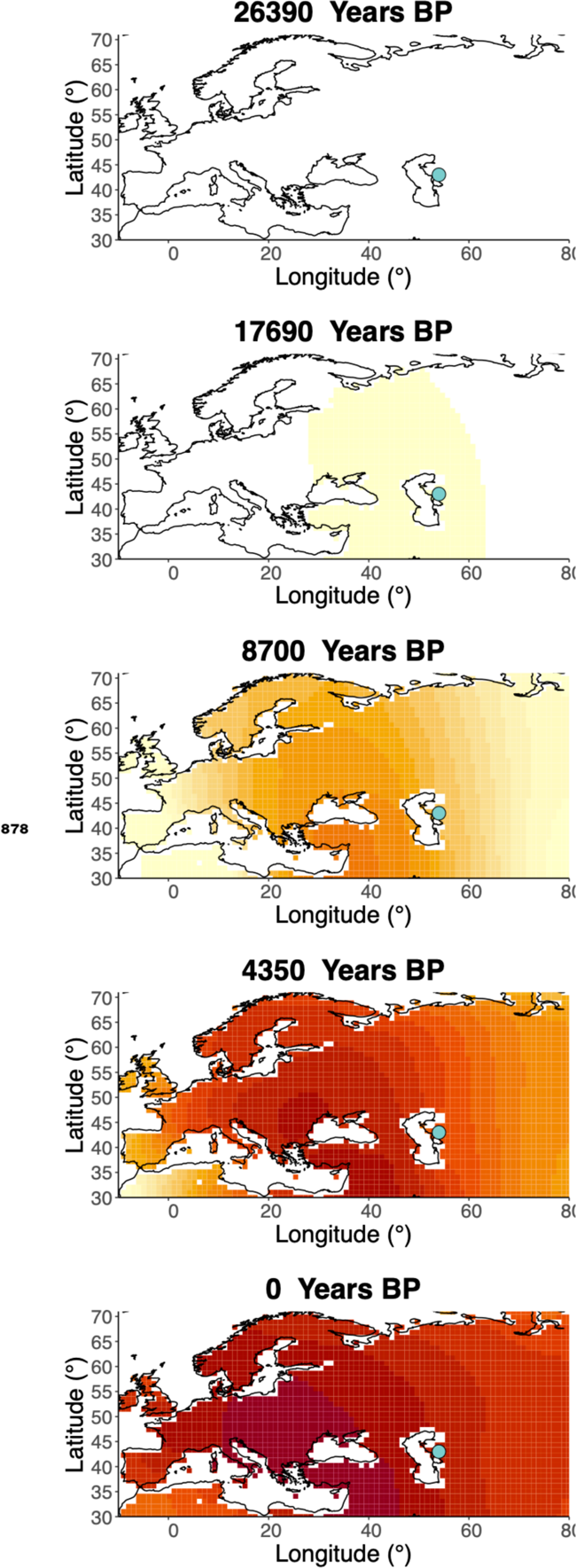
Inferred frequency dynamics of rs1042602(A) when the origin of the allele is moved 10 degrees east from the original estimate.

**Figure 8-Figure supplement 2.**
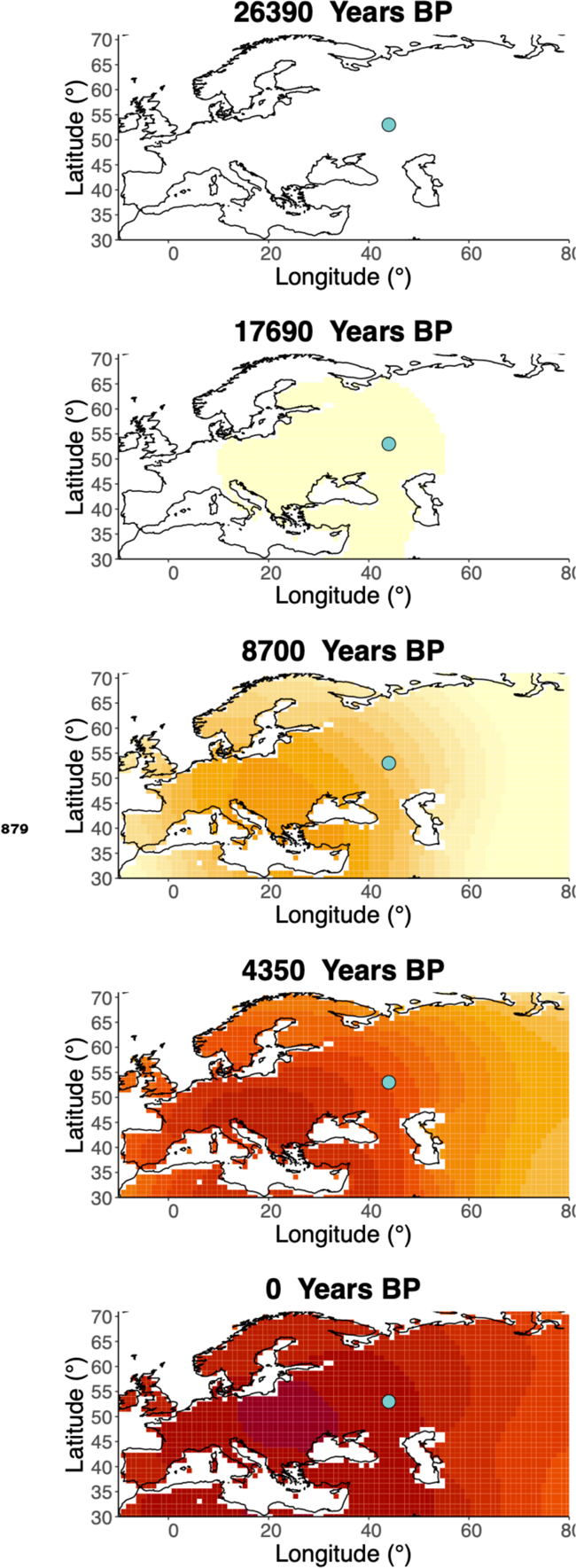
Inferred frequency dynamics of rs1042602(A) when the origin of the allele is moved 10 degrees north from the original estimate.

**Figure 8-Figure supplement 3.**
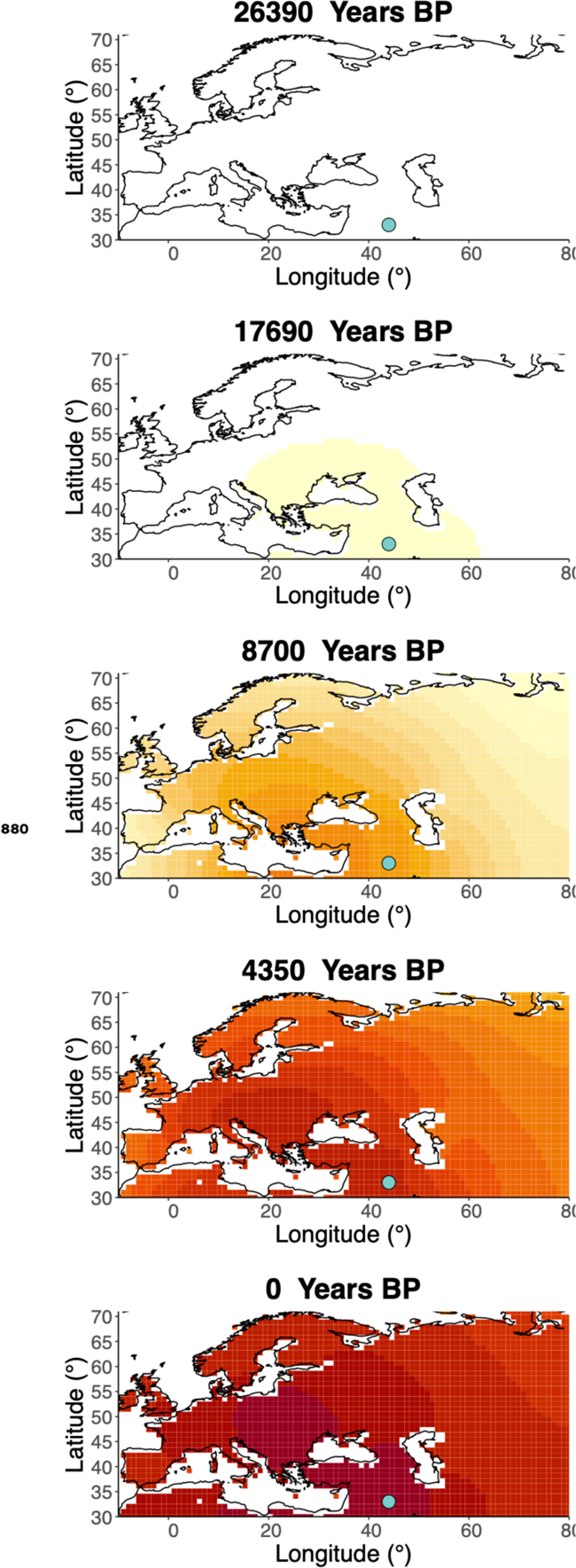
Inferred frequency dynamics of rs1042602(A) when the origin of the allele is moved 10 degrees south from the original estimate.

**Figure 8-Figure supplement 4.**
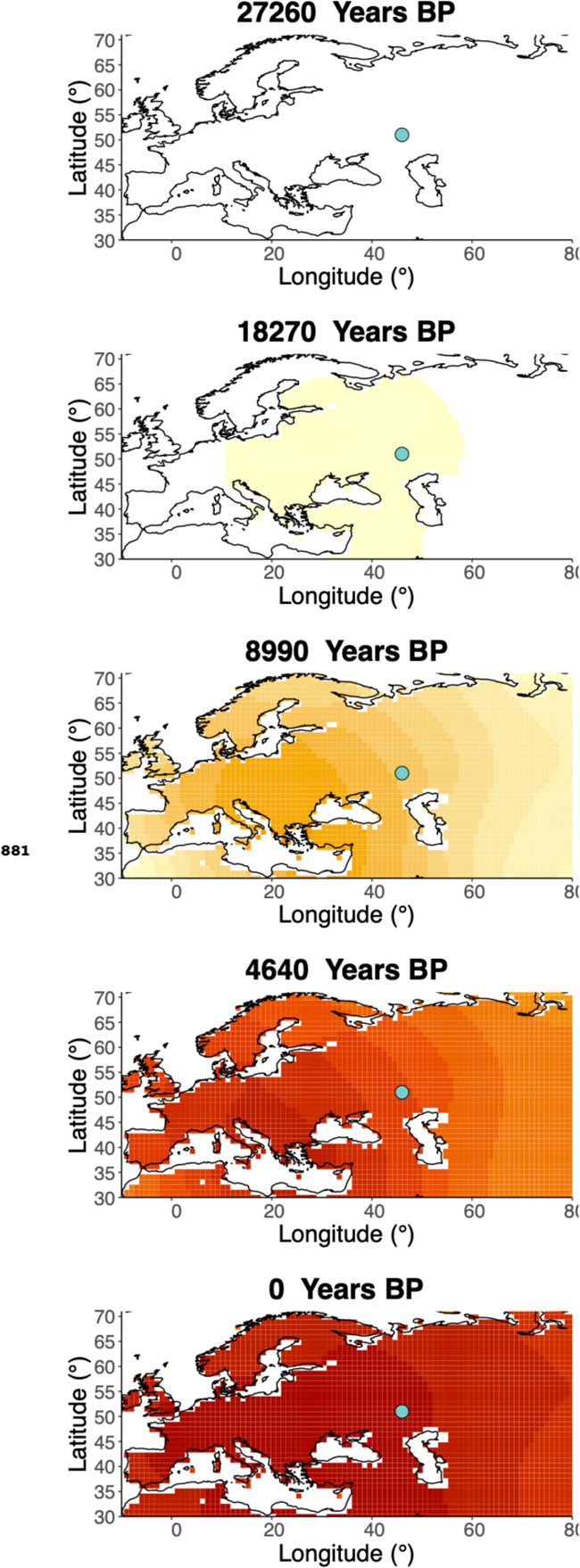
Inferred frequency dynamics of rs1042602(A) assuming the allele age to be the lower end of the 95% confidence interval for the allele age inferred in ***Albers and Mcvean (2020)*.**

**Figure 8-Figure supplement 5.**
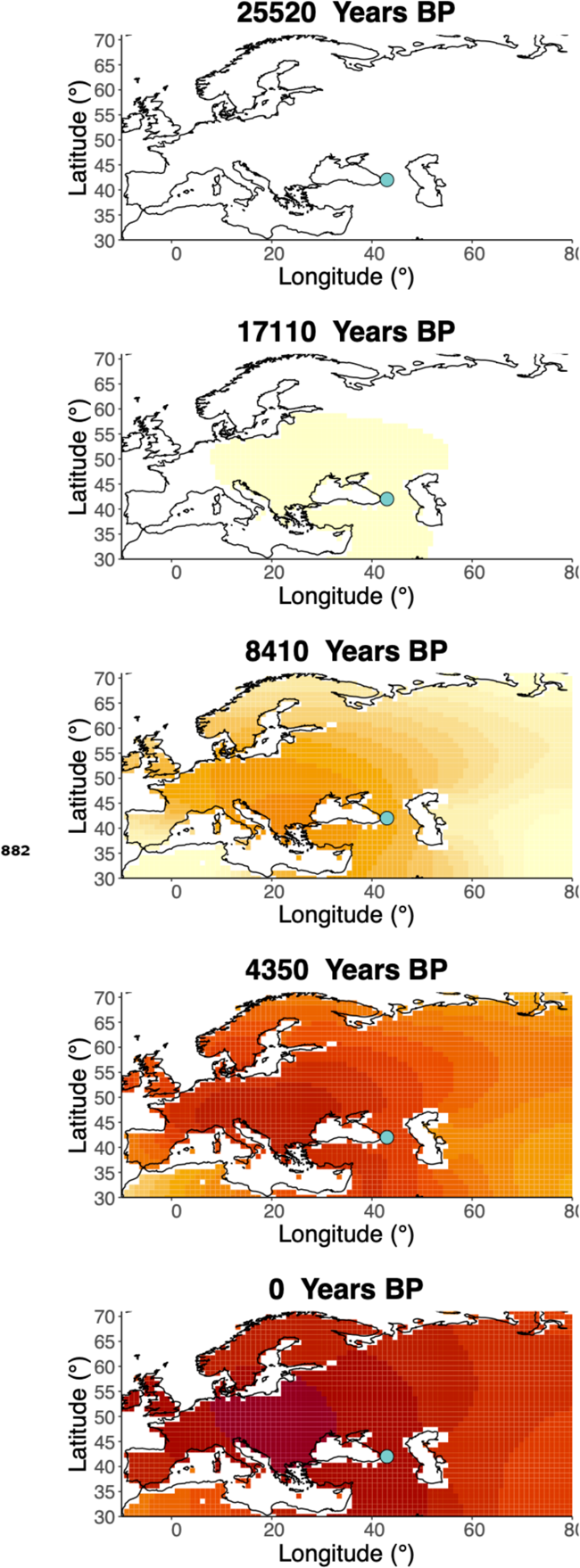
Frequency dynamics of rs1042602(A) assuming the allele age to be the higher end of the 95%confidence interval for the allele age inferred in ***Albers and Mcvean* (2020).**

**Figure 8-Figure supplement 6.**
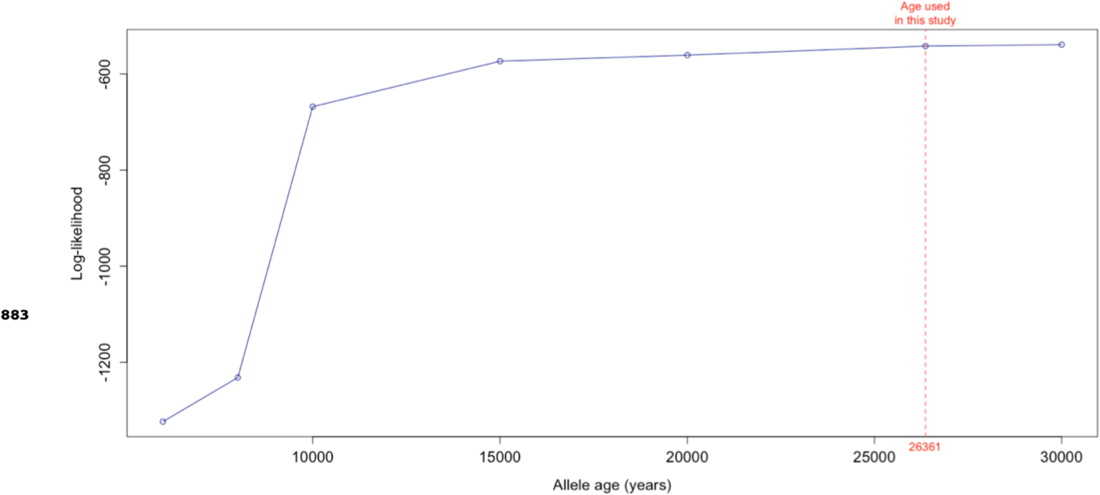
Log-likelihood values for model runs using different ages of the rs1042602(A) allele as input, with the age inferred by ***Albers and Mcvean (2020)*** we use as fixed input highlighted in red.

