## Supplementary Text 1 for "Modelling the spatiotemporal spread of beneficial alleles using ancient genomes"

**Publications that produced data included in the Allen Ancient DNA Resource, compiled by the Reich Lab:**

*[1KGPhase3]:*
***A global reference for human genetic variation.*** *1000 Genomes Project Consortium, Auton A, Brooks LD, Durbin RM, Garrison EP, Kang HM, Korbel JO, Marchini JL, McCarthy S, McVean GA, Abecasis GR. Nature. 2015 Oct 1;526(7571):68-74. doi: 10.1038/nature15393. PMID: 26432245.* [*http://www.internationalgenome.org/data/*](http://www.internationalgenome.org/data/) *[AgranatTamirCell2020]:*
***The Genomic History of the Bronze Age Southern Levant.*** *Agranat-Tamir L, Waldman S, Martin MAS, Gokhman D, Mishol N, Eshel T, Cheronet O, Rohland N, Mallick S, Adamski N, Lawson AM, Mah M, Michel M, Oppenheimer J, Stewardson K, Candilio F, Keating D, Gamarra B, Tzur S, Novak M, Kalisher R, Bechar S, Eshed V, Kennett DJ, Faerman M, Yahalom-Mack N, Monge JM, Govrin Y, Erel Y, Yakir B, Pinhasi R, Carmi S, Finkelstein I, Carmel L, Reich D.Cell. 2020 May 28;181(5):1146-1157.e11. doi: 10.1016/j.cell.2020.04.024. PMID: 32470400.

[AllentoftNature2015]:*
***Population genomics of Bronze Age Eurasia.*** *Allentoft ME, Sikora M, Sjögren KG, Rasmussen S, Rasmussen M, Stenderup J, Damgaard PB, Schroeder H, Ahlström T, Vinner L, Malaspinas AS, Margaryan A, Higham T, Chivall D, Lynnerup N, Harvig L, Baron J, Della Casa P, Dąbrowski P, Duffy PR, Ebel AV, Epimakhov A, Frei K, Furmanek M, Gralak T, Gromov A, Gronkiewicz S, Grupe G, Hajdu T, Jarysz R, Khartanovich V, Khokhlov A, Kiss V, Kolář J, Kriiska A, Lasak I, Longhi C, McGlynn G, Merkevicius A, Merkyte I, Metspalu M, Mkrtchyan R, Moiseyev V, Paja L, Pálfi G, Pokutta D, Pospieszny Ł, Price TD, Saag L, Sablin M, Shishlina N, Smrčka V, Soenov VI, Szeverényi V, Tóth G, Trifanova SV, Varul L, Vicze M, Yepiskoposyan L, Zhitenev V, Orlando L, Sicheritz-Pontén T, Brunak S, Nielsen R, Kristiansen K, Willerslev E. Nature. 2015 Jun 11;522(7555):167-72. doi: 10.1038/nature14507.

[AmorimNatureCommunications2018]:*
***Understanding 6th-century barbarian social organization and migration through paleogenomics.*** *Amorim CEG, Vai S, Posth C, Modi A, Koncz I, Hakenbeck S, La Rocca MC, Mende B, Bobo D, Pohl W, Baricco LP, Bedini E, Francalacci P, Giostra C, Vida T, Winger D, von Freeden U, Ghirotto S, Lari M, Barbujani G, Krause J, Caramelli D, Geary PJ, Veeramah KR. Nat Commun. 2018 Sep 11;9(1):3547. doi: 10.1038/s41467-018-06024-4.

[AntonioGaoMootsScience2019]:*
***Ancient Rome: A genetic crossroads of Europe and the Mediterranean.*** *Antonio ML, Gao Z, Moots HM, Lucci M, Candilio F, Sawyer S, Oberreiter V, Calderon D, Devitofranceschi K, Aikens RC, Aneli S, Bartoli F, Bedini A, Cheronet O, Cotter DJ, Fernandes DM, Gasperetti G, Grifoni R, Guidi A, La Pastina F, Loreti E, Manacorda D, Matullo G, Morretta S, Nava A, Fiocchi Nicolai V, Nomi F, Pavolini C, Pentiricci M, Pergola P, Piranomonte M, Schmidt R, Spinola G, Sperduti A, Rubini M, Bondioli L, Coppa A, Pinhasi R, Pritchard JK. Science. 2019 Nov 8;366(6466):708-714. doi: 10.1126/science.aay6826. PMID: 31699931.

[BergstromScience2020]:*
***Insights into human genetic variation and population history from 929 diverse genomes*** *Bergström A, McCarthy SA, Hui R, Almarri MA, Ayub Q, Danecek P, Chen Y, Felkel S, Hallast P, Kamm J, Blanché H, Deleuze JF, Cann H, Mallick S, Reich D, Sandhu MS, Skoglund P, Scally A, Xue Y, Durbin R, Tyler-Smith C. Science. 2020 Mar 20;367(6484):eaay5012. doi: 10.1126/science.aay5012. PMID: 32193295; PMCID: PMC7115999.

[BongersPNAS2020]:*
***Integration of ancient DNA with transdisciplinary dataset finds strong support for Inca resettlement in the south Peruvian coast.*** *Bongers JL, Nakatsuka N, O'Shea C, Harper TK, Tantaleán H, Stanish C, Fehren-Schmitz L. Proc Natl Acad Sci U S A. 2020 Aug 4;117(31):18359-18368. doi: 10.1073/pnas.2005965117. Epub 2020 Jul 13. PMID: 32661160; PMCID: PMC7414190.

[BraceNatureEcologyEvolution2019] / [BraceDiekmannNatureEcologyEvolution2019]:*
***Ancient genomes indicate population replacement in Early Neolithic Britain.*** *Brace S, Diekmann Y, Booth TJ, van Dorp L, Faltyskova Z, Rohland N, Mallick S, Olalde I, Ferry M, Michel M, Oppenheimer J, Broomandkhoshbacht N, Stewardson K, Martiniano R, Walsh S, Kayser M, Charlton S, Hellenthal G, Armit I, Schulting R, Craig OE, Sheridan A, Parker Pearson M, Stringer C, Reich D, Thomas MG, Barnes I. Nat Ecol Evol. 2019 May;3(5):765-771. doi: 10.1038/s41559-019-0871-9. Epub 2019 Apr 15. Erratum in: Nat Ecol Evol. 2019 May 8;:. PMID: 30988490.

[BroushakiScience2016]:*
***Early Neolithic genomes from the eastern Fertile Crescent.*** *Broushaki F, Thomas MG, Link V, López S, van Dorp L, Kirsanow K, Hofmanová Z, Diekmann Y, Cassidy LM, Díez-Del-Molino D, Kousathanas A, Sell C, Robson HK, Martiniano R, Blöcher J, Scheu A, Kreutzer S, Bollongino R, Bobo D, Davudi H, Munoz O, Currat M, Abdi K, Biglari F, Craig OE, Bradley DG, Shennan S, Veeramah K, Mashkour M, Wegmann D, Hellenthal G, Burger J. Science. 2016 Jul 29;353(6298):499-503. doi: 10.1126/science.aaf7943. Epub 2016 Jul 14. PubMed PMID: 27417496; PubMed Central PMCID: PMC5113750.

[BrunelPNAS2020]:*
***Ancient genomes from present-day France unveil 7,000 years of its demographic history.*** *Brunel S, Bennett EA, Cardin L, Garraud D, Barrand Emam H, Beylier A, Boulestin B, Chenal F, Ciesielski E, Convertini F, Dedet B, Desbrosse-Degobertiere S, Desenne S, Dubouloz J, Duday H, Escalon G, Fabre V, Gailledrat E, Gandelin M, Gleize Y, Goepfert S, Guilaine J, Hachem L, Ilett M, Lambach F, Maziere F, Perrin B, Plouin S, Pinard E, Praud I, Richard I, Riquier V, Roure R, Sendra B, Thevenet C, Thiol S, Vauquelin E, Vergnaud L, Grange T, Geigl EM, Pruvost M. Proc Natl Acad Sci U S A. 2020 Jun 9;117(23):12791-12798. doi: 10.1073/pnas.1918034117. Epub 2020 May 26. PMID: 32457149; PMCID: PMC7293694.

[CassidyPNAS2016]:*
***Neolithic and Bronze Age migration to Ireland and establishment of the insular Atlantic genome.*** *Cassidy LM, Martiniano R, Murphy EM, Teasdale MD, Mallory J, Hartwell B, Bradley DG. Proc Natl Acad Sci U S A. 2016 Jan 12;113(2):368-73. doi: 10.1073/pnas.1518445113. Epub 2015 Dec 28.

[CassidyNature2020]:*
***A dynastic elite in monumental Neolithic society.*** *Cassidy LM, Maoldúin RÓ, Kador T, Lynch A, Jones C, Woodman PC, Murphy E, Ramsey G, Dowd M, Noonan A, Campbell C, Jones ER, Mattiangeli V, Bradley DG. Nature. 2020 Jun;582(7812):384-388. doi: 10.1038/s41586-020-2378-6. Epub 2020 Jun 17. PMID: 32555485.

[DamgaardNature2018]:*
***137 ancient human genomes from across the Eurasian steppes.*** *Damgaard PB, Marchi N, Rasmussen S, Peyrot M, Renaud G, Korneliussen T, Moreno-Mayar JV, Pedersen MW, Goldberg A, Usmanova E, Baimukhanov N, Loman V, Hedeager L, Pedersen AG, Nielsen K, Afanasiev G, Akmatov K, Aldashev A, Alpaslan A, Baimbetov G, Bazaliiskii VI, Beisenov A, Boldbaatar B, Boldgiv B, Dorzhu C, Ellingvag S, Erdenebaatar D, Dajani R, Dmitriev E, Evdokimov V, Frei KM, Gromov A, Goryachev A, Hakonarson H, Hegay T, Khachatryan Z, Khaskhanov R, Kitov E, Kolbina A, Kubatbek T, Kukushkin A, Kukushkin I, Lau N, Margaryan A, Merkyte I, Mertz IV, Mertz VK, Mijiddorj E, Moiyesev V, Mukhtarova G, Nurmukhanbetov B, Orozbekova Z, Panyushkina I, Pieta K, Smrčka V, Shevnina I, Logvin A, Sjögren KG, Štolcová T, Taravella AM, Tashbaeva K, Tkachev A, Tulegenov T, Voyakin D, Yepiskoposyan L, Undrakhbold S, Varfolomeev V, Weber A, Wilson Sayres MA, Kradin N, Allentoft ME, Orlando L, Nielsen R, Sikora M, Heyer E, Kristiansen K, Willerslev E. Nature. 2018 May;557(7705):369-374. doi: 10.1038/s41586-018-0094-2. Epub 2018 May 9.

[DamgaardScience2018]:*
***The first horse herders and the impact of early Bronze Age steppe expansions into Asia.*** *de Barros Damgaard P, Martiniano R, Kamm J, Moreno-Mayar JV, Kroonen G, Peyrot M, Barjamovic G, Rasmussen S, Zacho C, Baimukhanov N, Zaibert V, Merz V, Biddanda A, Merz I, Loman V, Evdokimov V, Usmanova E, Hemphill B, Seguin-Orlando A, Yediay FE, Ullah I, Sjögren KG, Iversen KH, Choin J, de la Fuente C, Ilardo M, Schroeder H, Moiseyev V, Gromov A, Polyakov A, Omura S, Senyurt SY, Ahmad H, McKenzie C, Margaryan A, Hameed A, Samad A, Gul N, Khokhar MH, Goriunova OI, Bazaliiskii VI, Novembre J, Weber AW, Orlando L, Allentoft ME, Nielsen R, Kristiansen K, Sikora M, Outram AK, Durbin R, Willerslev E. Science. 2018 Jun 29;360(6396). pii: eaar7711. doi: 10.1126/science.aar7711. Epub 2018 May 9. PubMed PMID: 29743352.

[EbenesersdottirScience2018]*
***Ancient genomes from Iceland reveal the making of a human population.*** *Ebenesersdóttir SS, Sandoval-Velasco M, Gunnarsdóttir ED, Jagadeesan A, Guòmundsdóttir VB, Thordardóttir VB, Thordardóttir MS, Moore KHS, Siguròsson Á, Magnúsdóttir DN, Jónsson H, Snorradóttir S, Hovig E, Møller P, Kockum I, Olsson T, Alfredsson L, Hansen TF, Werge T, Cavalleri GL, Gilbert E, Lalueza-Fox C, Walser JW 3rd, Kristjánsdóttir S, Gopalakrishnan S, Árnadóttir L, Magnússon ÓP, Gilbert MTP, Stefánsson K, Helgason A. Science. 2018 Jun 1;360(6392):1028-1032. doi: 10.1126/science.aar2625. PMID: 29853688.

[FanGenomeBiology2019]:*
***African evolutionary history inferred from whole genome sequence data of 44 indigenous African populations.*** *Fan S, Kelly DE, Beltrame MH, Hansen MEB, Mallick S, Ranciaro A, Hirbo J, Thompson S, Beggs W, Nyambo T, Omar SA, Meskel DW, Belay G, Froment A, Patterson N, Reich D, Tishkoff SA. Genome Biol. 2019 Apr 26;20(1):82. doi: 10.1186/s13059-019-1679-2. Erratum in: Genome Biol. 2019 Oct 9;20(1):204. PMID: 31023338; PMCID: PMC6485071.

[FeldmanNatureCommunications2019]:*
***Late Pleistocene human genome suggests a local origin for the first farmers of central Anatolia.*** *Feldman M, Fernández-Domínguez E, Reynolds L, Baird D, Pearson J, Hershkovitz I, May H, Goring-Morris N, Benz M, Gresky J, Bianco RA, Fairbairn A, Mustafaolu G, Stockhammer PW, Posth C, Haak W, Jeong C, Krause J. Nat Commun. 2019 Mar 19;10(1):1218. doi: 10.1038/s41467-019-09209-7. PMID: 30890703.

[FeldmanScienceAdvances2019]:*
***Ancient DNA sheds light on the genetic origins of early Iron Age Philistines.*** *Feldman M, Master DM, Bianco RA, Burri M, Stockhammer PW, Mittnik A, Aja AJ, Jeong C, Krause J. Sci Adv. 2019 Jul 3;5(7):eaax0061. doi: 10.1126/sciadv.aax0061. eCollection 2019 Jul. PMID: 31281897.

[FernandesNatureEcologyEvolution2020]:*
***The spread of steppe and Iranian-related ancestry in the islands of the western Mediterranean.*** *Fernandes DM, Mittnik A, Olalde I, Lazaridis I, Cheronet O, Rohland N, Mallick S, Bernardos R, Broomandkhoshbacht N, Carlsson J, Culleton BJ, Ferry M, Gamarra B, Lari M, Mah M, Michel M, Modi A, Novak M, Oppenheimer J, Sirak KA, Stewardson K, Mandl K, Schattke C, Özdoğan KT, Lucci M, Gasperetti G, Candilio F, Salis G, Vai S, Camarós E, Calò C, Catalano G, Cueto M, Forgia V, Lozano M, Marini E, Micheletti M, Miccichè RM, Palombo MR, Ramis D, Schimmenti V, Sureda P, Teira L, Teschler-Nicola M, Kennett DJ, Lalueza-Fox C, Patterson N, Sineo L, Coppa A, Caramelli D, Pinhasi R, Reich D. Nat Ecol Evol. 2020 Mar;4(3):334-345. doi: 10.1038/s41559-020-1102-0. Epub 2020 Feb 24. Erratum in: Nat Ecol Evol. 2020 May;4(5):764. PMID: 32094539; PMCID: PMC7080320.

[FernandesSirakNature2020]:*
***A genetic history of the pre-contact Caribbean.*** *Fernandes DM, Sirak KA, Ringbauer H, Sedig J, Rohland N, Cheronet O, Mah M, Mallick S, Olalde I, Culleton BJ, Adamski N, Bernardos R, Bravo G, Broomandkhoshbacht N, Callan K, Candilio F, Demetz L, Carlson KSD, Eccles L, Freilich S, George RJ, Lawson AM, Mandl K, Marzaioli F, McCool WC, Oppenheimer J, Özdogan KT, Schattke C, Schmidt R, Stewardson K, Terrasi F, Zalzala F, Antúnez CA, Canosa EV, Colten R, Cucina A, Genchi F, Kraan C, La Pastina F, Lucci M, Maggiolo MV, Marcheco-Teruel B, Maria CT, Martínez C, París I, Pateman M, Simms TM, Sivoli CG, Vilar M, Kennett DJ, Keegan WF, Coppa A, Lipson M, Pinhasi R, Reich D. Nature. 2020 Dec 23. doi: 10.1038/s41586-020-03053-2. Epub ahead of print. PMID: 33361817.

[FurtwanglerNatureCommunications2020]:*
***Comparison of target enrichment strategies for ancient pathogen DNA.*** *Furtwängler A, Neukamm J, Böhme L, Reiter E, Vollstedt M, Arora N, Singh P, Cole ST, Knauf S, Calvignac-Spencer S, Krause-Kyora B, Krause J, Schuenemann VJ, Herbig A. Biotechniques. 2020 Dec;69(6):455-459. doi: 10.2144/btn-2020-0100. Epub 2020 Nov 2. PMID: 33135465.

[delaFuentePNAS2018]:*
***Genomic insights into the origin and diversification of late maritime hunter-gatherers from the Chilean Patagonia.*** *De la Fuente C, Ávila-Arcos MC, Galimany J, Carpenter ML, Homburger JR, Blanco A, Contreras P, Cruz Dávalos D, Reyes O, San Roman M, Moreno-Estrada A, Campos PF, Eng C, Huntsman S, Burchard EG, Malaspinas AS, Bustamante CD, Willerslev E, Llop E, Verdugo RA, Moraga M. Proc Natl Acad Sci U S A. 2018 Apr 24;115(17):E4006-E4012. doi: 10.1073/pnas.1715688115. Epub 2018 Apr 9. PubMed PMID: 29632188; PubMed Central PMCID: PMC5924884.

[FernandesScientificReports2018]:*
***A genomic Neolithic time transect of hunter-farmer admixture in central Poland.*** *Fernandes DM, Strapagiel D, Borówka P, Marciniak B, Żądzińska E, Sirak K, Siska V, Grygiel R, Carlsson J, Manica A, Lorkiewicz W, Pinhasi R. Sci Rep. 2018 Oct 5;8(1):14879. doi: 10.1038/s41598-018-33067-w.

[FlegontovNature2019]:*
***Palaeo-Eskimo genetic ancestry and the peopling of Chukotka and North America.*** *Flegontov P, Altinisik NE, Changmai P, Rohland N, Mallick S, Adamski N, Bolnick DA, Broomandkhoshbacht N, Candilio F, Culleton BJ, Flegontova O, Friesen TM, Jeong C, Harper TK, Keating D, Kennett DJ, Kim AM, Lamnidis TC, Lawson AM, Olalde I, Oppenheimer J, Potter BA, Raff J, Sattler RA, Skoglund P, Stewardson K, Vajda EJ, Vasilyev S, Veselovskaya E, Hayes MG, O'Rourke DH, Krause J, Pinhasi R, Reich D, Schiffels S. Nature. 2019 Jun;570(7760):236-240. doi: 10.1038/s41586-019-1251-y. Epub 2019 Jun 5. PMID: 31168094.

[FregelPNAS2018]:*
***Ancient genomes from North Africa evidence prehistoric migrations to the Maghreb from both the Levant and Europe.*** *Fregel R, Méndez FL, Bokbot Y, Martín-Socas D, Camalich-Massieu MD, Santana J, Morales J, Ávila-Arcos MC, Underhill PA, Shapiro B, Wojcik G, Rasmussen M, Soares AER, Kapp J, Sockell A, Rodríguez-Santos FJ, Mikdad A, Trujillo-Mederos A, Bustamante CD. Proc Natl Acad Sci U S A. 2018 Jun 26;115(26):6774-6779. doi: 10.1073/pnas.1800851115. Epub 2018 Jun 12. Erratum in: Proc Natl Acad Sci U S A. 2018 Jul 24;115(30):E7231. PubMed PMID: 29895688; PubMed Central PMCID: PMC6042094.

[FuCurrentBiology2013]: (contamMix used for mitochondrial contamination estimates)*
***A revised timescale for human evolution based on ancient mitochondrial genomes.*** *inFu Q, Mittnik A, Johnson PLF, Bos K, Lari M, Bollongino R, Sun C, Giemsch L, Schmitz R, Burger J, Ronchitelli AM, Martini F, Cremonesi RG, Svoboda J, Bauer P, Caramelli D, Castellano S, Reich D, Pääbo S, Krause J. Curr Biol. 2013 Apr 8;23(7):553-559. doi: 10.1016/j.cub.2013.02.044. Epub 2013 Mar 21. PMID: 23523248.

[FuNature2014]:*
***Genome sequence of a 45,000-year-old modern human from western Siberia.*** *Fu Q, Li H, Moorjani P, Jay F, Slepchenko SM, Bondarev AA, Johnson PL, Aximu-Petri A, Prüfer K, de Filippo C, Meyer M, Zwyns N, Salazar-García DC, Kuzmin YV, Keates SG, Kosintsev PA, Razhev DI, Richards MP, Peristov NV, Lachmann M, Douka K, Higham TF, Slatkin M, Hublin JJ, Reich D, Kelso J, Viola TB, Pääbo S. Nature. 2014 Oct 23;514(7523):445-9. doi: 10.1038/nature13810.

[FuNature2015]:*
***An early modern human from Romania with a recent Neanderthal ancestor.*** *Fu Q, Hajdinjak M, Moldovan OT, Constantin S, Mallick S, Skoglund P, Patterson N, Rohland N, Lazaridis I, Nickel B, Viola B, Prüfer K, Meyer M, Kelso J, Reich D, Pääbo S. Nature. 2015 Aug 13;524(7564):216-9. doi: 10.1038/nature14558. Epub 2015 Jun 22.

[FuNature2016]:*
***The genetic history of Ice Age Europe.*** *Fu Q, Posth C, Hajdinjak M, Petr M, Mallick S, Fernandes D, Furtwängler A, Haak W, Meyer M, Mittnik A, Nickel B, Peltzer A, Rohland N, Slon V, Talamo S, Lazaridis I, Lipson M, Mathieson I, Schiffels S, Skoglund P, Derevianko AP, Drozdov N, Slavinsky V, Tsybankov A, Cremonesi RG, Mallegni F, Gély B, Vacca E, Morales MR, Straus LG, Neugebauer-Maresch C, Teschler-Nicola M, Constantin S, Moldovan OT, Benazzi S, Peresani M, Coppola D, Lari M, Ricci S, Ronchitelli A, Valentin F, Thevenet C, Wehrberger K, Grigorescu D, Rougier H, Crevecoeur I, Flas D, Semal P, Mannino MA, Cupillard C, Bocherens H, Conard NJ, Harvati K, Moiseyev V, Drucker DG, Svoboda J, Richards MP, Caramelli D, Pinhasi R, Kelso J, Patterson N, Krause J, Pääbo S, Reich D. Nature. 2016 Jun 9;534(7606):200-5. doi: 10.1038/nature17993. Epub 2016 May 2.

[GambaNatureCommunications2014]:*
***Genome flux and stasis in a five millennium transect of European prehistory.*** *Gamba C, Jones ER, Teasdale MD, McLaughlin RL, Gonzalez-Fortes G, Mattiangeli V, Domboróczki L, Kővári I, Pap I, Anders A, Whittle A, Dani J, Raczky P, Higham TF, Hofreiter M, Bradley DG, Pinhasi R. Nat Commun. 2014 Oct 21;5:5257. doi: 10.1038/ncomms6257. PubMed PMID: 25334030; PubMed Central PMCID: PMC4218962.

[GokhmanNatureCommunications2020]:*
***Differential DNA methylation of vocal and facial anatomy genes in modern humans.*** *Gokhman D, Nissim-Rafinia M, Agranat-Tamir L, Housman G, García-Pérez R, Lizano E, Cheronet O, Mallick S, Nieves-Colón MA, Li H, Alpaslan-Roodenberg S, Novak M, Gu H, Osinski JM, Ferrando-Bernal M, Gelabert P, Lipende I, Mjungu D, Kondova I, Bontrop R, Kullmer O, Weber G, Shahar T, Dvir-Ginzberg M, Faerman M, Quillen EE, Meissner A, Lahav Y, Kandel L, Liebergall M, Prada ME, Vidal JM, Gronostajski RM, Stone AC, Yakir B, Lalueza-Fox C, Pinhasi R, Reich D, Marques-Bonet T, Meshorer E, Carmel L. Nat Commun. 2020 Mar 4;11(1):1189. doi: 10.1038/s41467-020-15020-6. PMID: 32132541; PMCID: PMC7055320.

[GonzalesFortesCurrentBiology2017]:*
***Paleogenomic Evidence for Multi-generational Mixing between Neolithic Farmers and Mesolithic Hunter-Gatherers in the Lower Danube Basin.*** *González-Fortes G, Jones ER, Lightfoot E, Bonsall C, Lazar C, Grandal-d'Anglade A, Garralda MD, Drak L, Siska V, Simalcsik A, Boroneanţ A, Vidal Romaní JR, Vaqueiro Rodríguez M, Arias P, Pinhasi R, Manica A, Hofreiter M. Paleogenomic Evidence for Multi-generational Mixing between Neolithic Farmers and Mesolithic Hunter-Gatherers in the Lower Danube Basin. Curr Biol. 2017 Jun 19;27(12):1801-1810.e10. doi: 10.1016/j.cub.2017.05.023. Epub 2017 May 25. PubMed PMID: 28552360; PubMed Central PMCID: PMC5483232.

[GonzalesFortesProcRoyalSocB2019]:*
***A western route of prehistoric human migration from Africa into the Iberian Peninsula.*** *González-Fortes G, Tassi F, Trucchi E, Henneberger K, Paijmans JLA, Díez-Del-Molino D, Schroeder H, Susca RR, Barroso-Ruíz C, Bermudez FJ, Barroso-Medina C, Bettencourt AMS, Sampaio HA, Grandal-d'Anglade A, Salas A, de Lombera-Hermida A, Fabregas Valcarce R, Vaquero M, Alonso S, Lozano M, Rodríguez-Alvarez XP, Fernández-Rodríguez C, Manica A, Hofreiter M, Barbujani G. Proc Biol Sci. 2019 Jan 30;286(1895):20182288. doi: 10.1098/rspb.2018.2288. PMID: 30963949.

[GuntherPLoSBiology2018]:*
***Population genomics of Mesolithic Scandinavia: Investigating early postglacial migration routes and high-latitude adaptation.*** *Günther T, Malmström H, Svensson EM, Omrak A, Sánchez-Quinto F, Kılınç GM, Krzewińska M, Eriksson G, Fraser M, Edlund H, Munters AR, Coutinho A, Simões LG, Vicente M, Sjölander A, Jansen Sellevold B, Jørgensen R, Claes P, Shriver MD, Valdiosera C, Netea MG, Apel J, Lidén K, Skar B, Storå J, Götherström A, Jakobsson M. PLoS Biol. 2018 Jan 9;16(1):e2003703. doi: 10.1371/journal.pbio.2003703. eCollection 2018 Jan.

[GuntherPNAS2015]:*
***Ancient genomes link early farmers from Atapuerca in Spain to modern-day Basques.*** *Günther T, Valdiosera C, Malmström H, Ureña I, Rodriguez-Varela R, Sverrisdóttir ÓO, Daskalaki EA, Skoglund P, Naidoo T, Svensson EM, Bermúdez de Castro JM, Carbonell E, Dunn M, Storå J, Iriarte E, Arsuaga JL, Carretero JM, Götherström A, Jakobsson M. Proc Natl Acad Sci U S A. 2015 Sep 22;112(38):11917-22. doi: 10.1073/pnas.1509851112. Epub 2015 Sep 8.

[HaberAJHG2017]:*
***Continuity and Admixture in the Last Five Millennia of Levantine History from Ancient Canaanite and Present-Day Lebanese Genome Sequences.*** *Haber M, Doumet-Serhal C, Scheib C, Xue Y, Danecek P, Mezzavilla M, Youhanna S, Martiniano R, Prado-Martinez J, Szpak M, Matisoo-Smith E, Schutkowski H, Mikulski R, Zalloua P, Kivisild T, Tyler-Smith C. Am J Hum Genet. 2017 Aug 3;101(2):274-282. doi: 10.1016/j.ajhg.2017.06.013. Epub 2017 Jul 27. PubMed PMID: 28757201; PubMed Central PMCID: PMC5544389.

[HaberAJHG2018]:*
***A Transient Pulse of Genetic Admixture from the Crusaders in the Near East Identified from Ancient Genome Sequences.*** *Haber M, Doumet-Serhal C, Scheib CL, Xue Y, Mikulski R, Martiniano R, Fischer-Genz B, Schutkowski H, Kivisild T, Tyler-Smith C. Am J Hum Genet. 2019 May 2;104(5):977-984. doi: 10.1016/j.ajhg.2019.03.015. Epub 2019 Apr 18. PMID: 31006515.

[HaberAJHG2020]:*
***A Genetic History of the Near East from an aDNA Time Course Sampling Eight Points in the Past 4,000 Years*** *Haber M, Nassar J, Almarri MA, Saupe T, Saag L, Griffith SJ, Doumet-Serhal C, Chanteau J, Saghieh-Beydoun M, Xue Y, Scheib CL, Tyler-Smith C. Am J Hum Genet. 2020 Jul 2;107(1):149-157. doi: 10.1016/j.ajhg.2020.05.008. Epub 2020 May 28. PMID: 32470374; PMCID: PMC7332655.

[HajdinjakNature2018]:*
***Reconstructing the genetic history of late Neanderthals.*** *Hajdinjak M, Fu Q, Hübner A, Petr M, Mafessoni F, Grote S, Skoglund P, Narasimham V, Rougier H, Crevecoeur I, Semal P, Soressi M, Talamo S, Hublin JJ, Gušić I, Kućan Ž, Rudan P, Golovanova LV, Doronichev VB, Posth C, Krause J, Korlević P, Nagel S, Nickel B, Slatkin M, Patterson N, Reich D, Prüfer K, Meyer M, Pääbo S, Kelso J. Nature. 2018 Mar 29;555(7698):652-656. doi: 10.1038/nature26151. Epub 2018 Mar 21.

[HarneyMayNatureCommunications2018]:*
***Ancient DNA from Chalcolithic Israel reveals the role of population mixture in cultural transformation.*** *Harney É, May H, Shalem D, Rohland N, Mallick S, Lazaridis I, Sarig R, Stewardson K, Nordenfelt S, Patterson N, Hershkovitz I, Reich D. Nat Commun. 2018 Aug 20;9(1):3336. doi: 10.1038/s41467-018-05649-9. Erratum in: Nat Commun. 2018 Sep 20;9(1):3913.

[HarneyNatureCommunications2019]:*
***Ancient DNA from the skeletons of Roopkund Lake reveals Mediterranean migrants in India.*** *Harney É, Nayak A, Patterson N, Joglekar P, Mushrif-Tripathy V, Mallick S, Rohland N, Sedig J, Adamski N, Bernardos R, Broomandkhoshbacht N, Culleton BJ, Ferry M, Harper TK, Michel M, Oppenheimer J, Stewardson K, Zhang Z, Harashawaradhana, Bartwal MS, Kumar S, Diyundi SC, Roberts P, Boivin N, Kennett DJ, Thangaraj K, Reich D, Rai N. Nat Commun. 2019 Aug 20;10(1):3670. doi: 10.1038/s41467-019-11357-9. PMID: 31431628.

[HofmanovaPNAS2016]:*
***Early farmers from across Europe directly descended from Neolithic Aegeans.*** *Hofmanová Z, Kreutzer S, Hellenthal G, Sell C, Diekmann Y, Díez-Del-Molino D, van Dorp L, López S, Kousathanas A, Link V, Kirsanow K, Cassidy LM, Martiniano R, Strobel M, Scheu A, Kotsakis K, Halstead P, Triantaphyllou S, Kyparissi-Apostolika N, Urem-Kotsou D, Ziota C, Adaktylou F, Gopalan S, Bobo DM, Winkelbach L, Blöcher J, Unterländer M, Leuenberger C, Çilingiroğlu Ç, Horejs B, Gerritsen F, Shennan SJ, Bradley DG, Currat M, Veeramah KR, Wegmann D, Thomas MG, Papageorgopoulou C, Burger J. Proc Natl Acad Sci U S A. 2016 Jun 21;113(25):6886-91. doi: 10.1073/pnas.1523951113. Epub 2016 Jun 6.

[JarveCurrentBiology2019]:*
***Shifts in the Genetic Landscape of the Western Eurasian Steppe Associated with the Beginning and End of the Scythian Dominance.*** *Järve M, Saag L, Scheib CL, Pathak AK, Montinaro F, Pagani L, Flores R, Guellil M, Saag L, Tambets K, Kushniarevich A, Solnik A, Varul L, Zadnikov S, Petrauskas O, Avramenko M, Magomedov B, Didenko S, Toshev G, Bruyako I, Grechko D, Okatenko V, Gorbenko K, Smyrnov O, Heiko A, Reida R, Sapiehin S, Sirotin S, Tairov A, Beisenov A, Starodubtsev M, Vasilev V, Nechvaloda A, Atabiev B, Litvinov S, Ekomasova N, Dzhaubermezov M, Voroniatov S, Utevska O, Shramko I, Khusnutdinova E, Metspalu M, Savelev N, Kriiska A, Kivisild T, Villems R. Curr Biol. 2019 Jul 22;29(14):2430-2441.e10. doi: 10.1016/j.cub.2019.06.019. Epub 2019 Jul 11. PMID: 31303491.

[JeongNatureEcologyEvolution2019]:*
***The genetic history of admixture across inner Eurasia.*** *Jeong C, Balanovsky O, Lukianova E, Kahbatkyzy N, Flegontov P, Zaporozhchenko V, Immel A, Wang CC, Ixan O, Khussainova E, Bekmanov B, Zaibert V, Lavryashina M, Pocheshkhova E, Yusupov Y, Agdzhoyan A, Koshel S, Bukin A, Nymadawa P, Turdikulova S, Dalimova D, Churnosov M, Skhalyakho R, Daragan D, Bogunov Y, Bogunova A, Shtrunov A, Dubova N, Zhabagin M, Yepiskoposyan L, Churakov V, Pislegin N, Damba L, Saroyants L, Dibirova K, Atramentova L, Utevska O, Idrisov E, Kamenshchikova E, Evseeva I, Metspalu M, Outram AK, Robbeets M, Djansugurova L, Balanovska E, Schiffels S, Haak W, Reich D, Krause J. Nat Ecol Evol. 2019 Jun;3(6):966-976. doi: 10.1038/s41559-019-0878-2. Epub 2019 Apr 29. PMID: 31036896.

[JeongPNAS2016]:*
***Long-term genetic stability and a high-altitude East Asian origin for the peoples of the high valleys of the Himalayan arc.*** *Jeong C, Ozga AT, Witonsky DB, Malmström H, Edlund H, Hofman CA, Hagan RW, Jakobsson M, Lewis CM, Aldenderfer MS, Di Rienzo A, Warinner C. Proc Natl Acad Sci U S A. 2016 Jul 5;113(27):7485-90. doi: 10.1073/pnas.1520844113. Epub 2016 Jun 20. PMID: 27325755.

[JeongPNAS2018]*
***Bronze Age population dynamics and the rise of dairy pastoralism on the eastern Eurasian steppe.*** *Jeong C, Wilkin S, Amgalantugs T, Bouwman AS, Taylor WTT, Hagan RW, Bromage S, Tsolmon S, Trachsel C, Grossmann J, Littleton J, Makarewicz CA, Krigbaum J, Burri M, Scott A, Davaasambuu G, Wright J, Irmer F, Myagmar E, Boivin N, Robbeets M, Rühli FJ, Krause J, Frohlich B, Hendy J, Warinner C. Proc Natl Acad Sci U S A. 2018 Nov 27;115(48):E11248-E11255. doi: 10.1073/pnas.1813608115. Epub 2018 Nov 5. PMID: 30397125.

[JonesNatureCommunications2015]:*
***Upper Palaeolithic genomes reveal deep roots of modern Eurasians.*** *Jones ER, Gonzalez-Fortes G, Connell S, Siska V, Eriksson A, Martiniano R, McLaughlin RL, Gallego Llorente M, Cassidy LM, Gamba C, Meshveliani T, Bar-Yosef O, Müller W, Belfer-Cohen A, Matskevich Z, Jakeli N, Higham TF, Currat M, Lordkipanidze D, Hofreiter M, Manica A, Pinhasi R, Bradley DG. Nat Commun. 2015 Nov 16;6:8912. doi: 10.1038/ncomms9912. PMID: 26567969.

[JonesCurrentBiology2017]:*
***The Neolithic Transition in the Baltic Was Not Driven by Admixture with Early European Farmers.*** *Jones ER, Zarina G, Moiseyev V, Lightfoot E, Nigst PR, Manica A, Pinhasi R, Bradley DG. Curr Biol. 2017 Feb 20;27(4):576-582.

[KanzawaKiriyamaJHG2016:]*
***A partial nuclear genome of the Jomons who lived 3000 years ago in Fukushima, Japan.*** *Kanzawa-Kiriyama H, Kryukov K, Jinam TA, Hosomichi K, Saso A, Suwa G, Ueda S, Yoneda M, Tajima A, Shinoda KI, Inoue I, Saitou N. J Hum Genet. 2017 Feb;62(2):213-221. doi: 10.1038/jhg.2016.110. Epub 2016 Sep 1.

[KellerNatureCommunications2012]:*
***New insights into the Tyrolean Iceman's origin and phenotype as inferred by whole-genome sequencing.*** *Keller A1, Graefen A, Ball M, Matzas M, Boisguerin V, Maixner F, Leidinger P, Backes C, Khairat R, Forster M, Stade B, Franke A, Mayer J, Spangler J, McLaughlin S, Shah M, Lee C, Harkins TT, Sartori A, Moreno-Estrada A, Henn B, Sikora M, Semino O, Chiaroni J, Rootsi S, Myres NM, Cabrera VM, Underhill PA, Bustamante CD, Vigl EE, Samadelli M, Cipollini G, Haas J, Katus H, O'Connor BD, Carlson MR, Meder B, Blin N, Meese E, Pusch CM, Zink A. Nat Commun. 2012 Feb 28;3:698. doi: 10.1038/ncomms1701.Paperpile.

[KennettNatureCommunications2017]:*
***Archaeogenomic evidence reveals prehistoric matrilineal dynasty.*** *Kennett DJ, Plog S, George RJ, Culleton BJ, Watson AS, Skoglund P, Rohland N, Mallick S, Stewardson K, Kistler L, LeBlanc SA, Whiteley PM, Reich D, Perry GH. Nat Commun. 2017 Feb 21;8:14115. doi: 10.1038/ncomms14115. PMID: 28221340.

[KilincCurrentBiology2016]:*
***The Demographic Development of the First Farmers in Anatolia.*** *Kılınç GM, Omrak A, Özer F, Günther T, Büyükkarakaya AM, Bıçakçı E, Baird D, Dönertaş HM, Ghalichi A, Yaka R, Koptekin D, Açan SC, Parvizi P, Krzewińska M, Daskalaki EA, Yüncü E, Dağtaş ND, Fairbairn A, Pearson J, Mustafaoğlu G, Erdal YS, Çakan YG, Togan İ, Somel M, Storå J, Jakobsson M, Götherström A. Curr Biol. 2016 Oct 10;26(19):2659-2666. doi: 10.1016/j.cub.2016.07.057. Epub 2016 Aug 4. PMID: 27498567.

[Korneliussen2014]: (used for X-contamination estimates)*
***ANGSD: Analysis of Next Generation Sequencing Data.*** *Korneliussen TS, Albrechtsen A, Nielsen R. BMC Bioinformatics. 2014 Nov 25;15:356. doi: 10.1186/s12859-014-0356-4. PMID: 25420514.

[KrzewinskaCurrentBiology2018]:*
***Genomic and Strontium Isotope Variation Reveal Immigration Patterns in a Viking Age Town.*** *Krzewińska M, Kjellström A, Günther T, Hedenstierna-Jonson C, Zachrisson T, Omrak A, Yaka R, Kılınç GM, Somel M, Sobrado V, Evans J, Knipper C, Jakobsson M, Storå J, Götherström A. Curr Biol. 2018 Sep 10;28(17):2730-2738.e10. doi: 10.1016/j.cub.2018.06.053. Epub 2018 Aug 23. PMID: 30146150.

[KrzewinskaScienceAdvances2018]:*
***Ancient genomes suggest the eastern Pontic-Caspian steppe as the source of western Iron Age nomads.*** *Krzewińska M, Kılınç GM, Juras A, Koptekin D, Chyleński M, Nikitin AG, Shcherbakov N, Shuteleva I, Leonova T, Kraeva L, Sungatov FA, Sultanova AN, Potekhina I, Łukasik S, Krenz-Niedbała M, Dalén L, Sinika V, Jakobsson M, Storå J, Götherström A. Sci Adv. 2018 Oct 3;4(10):eaat4457. doi: 10.1126/sciadv.aat4457. eCollection 2018 Oct. PMID: 30417088.

[LamnidisNatureCommunications2018]:*
***Ancient Fennoscandian genomes reveal origin and spread of Siberian ancestry in Europe.*** *Lamnidis TC, Majander K, Jeong C, Salmela E, Wessman A, Moiseyev V, Khartanovich V, Balanovsky O, Ongyerth M, Weihmann A, Sajantila A, Kelso J, Pääbo S, Onkamo P, Haak W, Krause J, Schiffels S. Nat Commun. 2018 Nov 27;9(1):5018. doi: 10.1038/s41467-018-07483-5. PMID: 30479341.

[LazaridisNature2014]:*
***Ancient human genomes suggest three ancestral populations for present-day Europeans.*** *Lazaridis I, Patterson N, Mittnik A, Renaud G, Mallick S, Kirsanow K, Sudmant PH, Schraiber JG, Castellano S, Lipson M, Berger B, Economou C, Bollongino R, Fu Q, Bos KI, Nordenfelt S, Li H, de Filippo C, Prüfer K, Sawyer S, Posth C, Haak W, Hallgren F, Fornander E, Rohland N, Delsate D, Francken M, Guinet JM, Wahl J, Ayodo G, Babiker HA, Bailliet G, Balanovska E, Balanovsky O, Barrantes R, Bedoya G, Ben-Ami H, Bene J, Berrada F, Bravi CM, Brisighelli F, Busby GB, Cali F, Churnosov M, Cole DE, Corach D, Damba L, van Driem G, Dryomov S, Dugoujon JM, Fedorova SA, Gallego Romero I, Gubina M, Hammer M, Henn BM, Hervig T, Hodoglugil U, Jha AR, Karachanak-Yankova S, Khusainova R, Khusnutdinova E, Kittles R, Kivisild T, Klitz W, Kučinskas V, Kushniarevich A, Laredj L, Litvinov S, Loukidis T, Mahley RW, Melegh B, Metspalu E, Molina J, Mountain J, Näkkäläjärvi K, Nesheva D, Nyambo T, Osipova L, Parik J, Platonov F, Posukh O, Romano V, Rothhammer F, Rudan I, Ruizbakiev R, Sahakyan H, Sajantila A, Salas A, Starikovskaya EB, Tarekegn A, Toncheva D, Turdikulova S, Uktveryte I, Utevska O, Vasquez R, Villena M, Voevoda M, Winkler CA, Yepiskoposyan L, Zalloua P, Zemunik T, Cooper A, Capelli C, Thomas MG, Ruiz-Linares A, Tishkoff SA, Singh L, Thangaraj K, Villems R, Comas D, Sukernik R, Metspalu M, Meyer M, Eichler EE, Burger J, Slatkin M, Pääbo S, Kelso J, Reich D, Krause J. Nature. 2014 Sep 18;513(7518):409-13. doi: 10.1038/nature13673. PMID: 25230663.

[LazaridisNature2016]:*
***Genomic insights into the origin of farming in the ancient Near East.*** *Lazaridis I, Nadel D, Rollefson G, Merrett DC, Rohland N, Mallick S, Fernandes D, Novak M, Gamarra B, Sirak K, Connell S, Stewardson K, Harney E, Fu Q, Gonzalez-Fortes G, Jones ER, Roodenberg SA, Lengyel G, Bocquentin F, Gasparian B, Mongce JM, Gregg M, Eshed V, Mizrahi AS, Meiklejohn C, Gerritsen F, Bejenaru L, Blüher M, Campbell A, Cavalleri G, Comas D, Froguel P, Gilbert E, Kerr SM, Kovacs P, Krause J, McGettigan D, Merrigan M, Merriwether DA, O'Reilly S, Richards MB, Semino O, Shamoon-Pour M, Stefanescu G, Stumvoll M, Tönjes A, Torroni A, Wilson JF, Yengo L, Hovhannisyan NA, Patterson N, Pinhasi R, Reich D. Nature. 2016 Aug 25;536(7617):419-24. Epub 2016 Jul 25. PMID: 27459054.

[LazaridisNature2017]:*
***Genetic origins of the Minoans and Mycenaeans.*** *Lazaridis I, Mittnik A, Patterson N, Mallick S, Rohland N, Pfrengle S, Furtwängler A, Peltzer A, Posth C, Vasilakis A, McGeorge PJP, Konsolaki-Yannopoulou E, Korres G, Martlew H, Michalodimitrakis M, Özsait M, Özsait N, Papathanasiou A, Richards M, Roodenberg SA, Tzedakis Y, Arnott R, Fernandes DM, Hughey JR, Lotakis DM, Navas PA, Maniatis Y, Stamatoyannopoulos JA, Stewardson K, Stockhammer P, Pinhasi R, Reich D, Krause J, Stamatoyannopoulos G. Nature. 2017 Aug 10;548(7666):214-218. doi: 10.1038/nature23310. Epub 2017 Aug 2. PMID: 28783727.

[LindoPNAS2017]:*
***Ancient individuals from the North American Northwest Coast reveal 10,000 years of regional genetic continuity.*** *Lindo J, Achilli A, Perego UA, Archer D, Valdiosera C, Petzelt B, Mitchell J, Worl R, Dixon EJ, Fifield TE, Rasmussen M, Willerslev E, Cybulski JS, Kemp BM, DeGiorgio M, Malhi RS. Proc Natl Acad Sci U S A. 2017 Apr 18;114(16):4093-4098. doi: 10.1073/pnas.1620410114. Epub 2017 Apr 4. PMID: 28377518.

[LindoScienceAdvances2018]:*
***The genetic prehistory of the Andean highlands 7000 years BP though European contact.*** *Lindo J, Haas R, Hofman C, Apata M, Moraga M, Verdugo RA, Watson JT, Viviano Llave C, Witonsky D, Beall C, Warinner C, Novembre J, Aldenderfer M, Di Rienzo A. Sci Adv. 2018 Nov 8;4(11):eaau4921. doi: 10.1126/sciadv.aau4921. eCollection 2018 Nov. PMID: 30417096.

[LinderholmNatureScientificReports2020]:*
***Corded Ware cultural complexity uncovered using genomic and isotopic analysis from south-eastern Poland.*** *Linderholm A, Kılınç GM, Szczepanek A, Włodarczak P, Jarosz P, Belka Z, Dopieralska J, Werens K, Górski J, Mazurek M, Hozer M, Rybicka M, Ostrowski M, Bagińska J, Koman W, Rodríguez-Varela R, Storå J, Götherström A, Krzewińska M. Sci Rep. 2020 Apr 14;10(1):6885. doi: 10.1038/s41598-020-63138-w. PMID: 32303690; PMCID: PMC7165176.

[LipsonCurrentBiology2018]:*
***Population Turnover in Remote Oceania Shortly after Initial Settlement.*** *Lipson M, Skoglund P, Spriggs M, Valentin F, Bedford S, Shing R, Buckley H, Phillip I, Ward GK, Mallick S, Rohland N, Broomandkhoshbacht N, Cheronet O, Ferry M, Harper TK, Michel M, Oppenheimer J, Sirak K, Stewardson K, Auckland K, Hill AVS, Maitland K, Oppenheimer SJ, Parks T, Robson K, Williams TN, Kennett DJ, Mentzer AJ, Pinhasi R, Reich D. Curr Biol. 2018 Apr 2;28(7):1157-1165.e7. doi: 10.1016/j.cub.2018.02.051. Epub 2018 Feb 28. PMID:29501328.

[LipsonNature2017]:*
***Parallel palaeogenomic transects reveal complex genetic history of early European farmers.*** *Lipson M, Szécsényi-Nagy A, Mallick S, Pósa A, Stégmár B, Keerl V, Rohland N, Stewardson K, Ferry M, Michel M, Oppenheimer J, Broomandkhoshbacht N, Harney E, Nordenfelt S, Llamas B, Gusztáv Mende B, Köhler K, Oross K, Bondár M, Marton T, Osztás A, Jakucs J, Paluch T, Horváth F, Csengeri P, Koós J, Sebők K, Anders A, Raczky P, Regenye J, Barna JP, Fábián S, Serlegi G, Toldi Z, GyÖngyvér Nagy E, Dani J, Molnár E, Pálfi G, Márk L, Melegh B, Bánfai Z, Domboróczki L, Fernández-Eraso J, Antonio Mujika-Alustiza J, Alonso Fernández C, Jiménez Echevarría J, Bollongino R, Orschiedt J, Schierhold K, Meller H, Cooper A, Burger J, Bánffy E, Alt KW, Lalueza-Fox C, Haak W, Reich D. Nature. 2017 Nov 16;551(7680):368-372. doi: 10.1038/nature24476. Epub 2017 Nov 8. PMID: 29144465.

[LipsonNature2020]:*
***Ancient West African foragers in the context of African population history.*** *Lipson M, Ribot I, Mallick S, Rohland N, Olalde I, Adamski N, Broomandkhoshbacht N, Lawson AM, López S, Oppenheimer J, Stewardson K, Asombang RN, Bocherens H, Bradman N, Culleton BJ, Cornelissen E, Crevecoeur I, de Maret P, Fomine FLM, Lavachery P, Mindzie CM, Orban R, Sawchuk E, Semal P, Thomas MG, Van Neer W, Veeramah KR, Kennett DJ, Patterson N, Hellenthal G, Lalueza-Fox C, MacEachern S, Prendergast ME, Reich D. Nature. 2020 Jan;577(7792):665-670. doi: 10.1038/s41586-020-1929-1. Epub 2020 Jan 22. PMID: 31969706.

[LipsonScience2018]:*
***Ancient genomes document multiple waves of migration in Southeast Asian prehistory.*** *Lipson M, Cheronet O, Mallick S, Rohland N, Oxenham M, Pietrusewsky M, Pryce TO, Willis A, Matsumura H, Buckley H, Domett K, Nguyen GH, Trinh HH, Kyaw AA, Win TT, Pradier B, Broomandkhoshbacht N, Candilio F, Changmai P, Fernandes D, Ferry M, Gamarra B, Harney E, Kampuansai J, Kutanan W, Michel M, Novak M, Oppenheimer J, Sirak K, Stewardson K, Zhang Z, Flegontov P, Pinhasi R, Reich D. Science. 2018 Jul 6;361(6397):92-95. doi: 10.1126/science.aat3188. Epub 2018 May 17. PMID: 29773666.

[LlorenteScience2015]:*
***Ancient Ethiopian genome reveals extensive Eurasian admixture throughout the African continent.*** *Gallego Llorente M, Jones ER, Eriksson A, Siska V, Arthur KW, Arthur JW, Curtis MC, Stock JT, Coltorti M, Pieruccini P, Stretton S, Brock F, Higham T, Park Y, Hofreiter M, Bradley DG, Bhak J, Pinhasi R, Manica A. Science. 2015 Nov 13;350(6262):820-2. doi: 10.1126/science.aad2879. Epub 2015 Oct 8. Erratum in: Science. 2016 Feb 19;351(6275). pii: aaf3945. doi: 10.1126/science.aaf3945. PMID: 26449472.

[MafessoniPNAS2020]:*
***A high-coverage Neandertal genome from Chagyrskaya Cave*** *Mafessoni F, Grote S, de Filippo C, Slon V, Kolobova KA, Viola B, Markin SV, Chintalapati M, Peyrégne S, Skov L, Skoglund P, Krivoshapkin AI, Derevianko AP, Meyer M, Kelso J, Peter B, Prüfer K, Pääbo S. Proc Natl Acad Sci U S A. 2020 Jun 30;117(26):15132-15136. doi: 10.1073/pnas.2004944117. Epub 2020 Jun 16. PMID: 32546518; PMCID: PMC7334501.

[MalaspinasCurrentBiology2014]:*
***Two ancient human genomes reveal Polynesian ancestry among the indigenous Botocudos of Brazil.*** *Malaspinas AS, Lao O, Schroeder H, Rasmussen M, Raghavan M, Moltke I, Campos PF, Sagredo FS, Rasmussen S, Gonçalves VF, Albrechtsen A, Allentoft ME, Johnson PL, Li M, Reis S, Bernardo DV, DeGiorgio M, Duggan AT, Bastos M, Wang Y, Stenderup J, Moreno-Mayar JV, Brunak S, Sicheritz-Ponten T, Hodges E, Hannon GJ, Orlando L, Price TD, Jensen JD, Nielsen R, Heinemeier J, Olsen J, Rodrigues-Carvalho C, Lahr MM, Neves WA, Kayser M, Higham T, Stoneking M, Pena SD, Willerslev E. Curr Biol. 2014 Nov 3;24(21):R1035-7. doi: 10.1016/j.cub.2014.09.078. Epub 2014 Oct 23. PMID: 25455029.

[MallickNature2016]:*
***The Simons Genome Diversity Project: 300 genomes from 142 diverse populations.*** *Mallick S, Li H, Lipson M, Mathieson I, Gymrek M, Racimo F, Zhao M, Chennagiri N, Nordenfelt S, Tandon A, Skoglund P, Lazaridis I, Sankararaman S, Fu Q, Rohland N, Renaud G, Erlich Y, Willems T, Gallo C, Spence JP, Song YS, Poletti G, Balloux F, van Driem G, de Knijff P, Romero IG, Jha AR, Behar DM, Bravi CM, Capelli C, Hervig T, Moreno-Estrada A, Posukh OL, Balanovska E, Balanovsky O, Karachanak-Yankova S, Sahakyan H, Toncheva D, Yepiskoposyan L, Tyler-Smith C, Xue Y, Abdullah MS, Ruiz-Linares A, Beall CM, Di Rienzo A, Jeong C, Starikovskaya EB, Metspalu E, Parik J, Villems R, Henn BM, Hodoglugil U, Mahley R, Sajantila A, Stamatoyannopoulos G, Wee JT, Khusainova R, Khusnutdinova E, Litvinov S, Ayodo G, Comas D, Hammer MF, Kivisild T, Klitz W, Winkler CA, Labuda D, Bamshad M, Jorde LB, Tishkoff SA, Watkins WS, Metspalu M, Dryomov S, Sukernik R, Singh L, Thangaraj K, Pääbo S, Kelso J, Patterson N, Reich D. Nature. 2016 Oct 13;538(7624):201-206. doi: 10.1038/nature18964. Epub 2016 Sep 21. PMID: 27654912.

[MalmstromProcBiolSci2019]:*
***The genomic ancestry of the Scandinavian Battle Axe Culture people and their relation to the broader Corded Ware horizon.*** *Malmström H, Günther T, Svensson EM, Juras A, Fraser M, Munters AR, Pospieszny L , Tõrv M, Lindström J, Götherström A, Storå J, Jakobsson M. Proc Biol Sci. 2019 Oct 9;286(1912):20191528. doi: 10.1098/rspb.2019.1528. Epub 2019 Oct 9. PMID: 31594508.

[MartinianoNatureCommunications2016]:*
***Genomic signals of migration and continuity in Britain before the Anglo-Saxons.*** *Martiniano R, Caffell A, Holst M, Hunter-Mann K, Montgomery J, Müldner G, McLaughlin RL, Teasdale MD, van Rheenen W, Veldink JH, van den Berg LH, Hardiman O, Carroll M, Roskams S, Oxley J, Morgan C, Thomas MG, Barnes I, McDonnell C, Collins MJ, Bradley DG. Nat Commun. 2016 Jan 19;7:10326. doi: 10.1038/ncomms10326. PMID: 26783717.

[MarcusNatureCommunications2020]:*
***Genetic history from the Middle Neolithic to present on the Mediterranean island of Sardinia.*** *Marcus JH, Posth C, Ringbauer H, Lai L, Skeates R, Sidore C, Beckett J, Furtwängler A, Olivieri A, Chiang CWK, Al-Asadi H, Dey K, Joseph TA, Liu CC, Der Sarkissian C, Radzevičiūtė R, Michel M, Gradoli MG, Marongiu P, Rubino S, Mazzarello V, Rovina D, La Fragola A, Serra RM, Bandiera P, Bianucci R, Pompianu E, Murgia C, Guirguis M, Orquin RP, Tuross N, van Dommelen P, Haak W, Reich D, Schlessinger D, Cucca F, Krause J, Novembre J. Nat Commun. 2020 Feb 24;11(1):939. doi: 10.1038/s41467-020-14523-6. PMID: 32094358; PMCID: PMC7039977.

[MargaryanWillerslevNature2020]:*
***Population genomics of the Viking world.*** *Margaryan A, Lawson DJ, Sikora M, Racimo F, Rasmussen S, Moltke I, Cassidy LM, Jørsboe E, Ingason A, Pedersen MW, Korneliussen T, Wilhelmson H, Buś MM, de Barros Damgaard P, Martiniano R, Renaud G, Bhérer C, Moreno-Mayar JV, Fotakis AK, Allen M, Allmäe R, Molak M, Cappellini E, Scorrano G, McColl H, Buzhilova A, Fox A, Albrechtsen A, Schütz B, Skar B, Arcini C, Falys C, Jonson CH, Błaszczyk D, Pezhemsky D, Turner-Walker G, Gestsdóttir H, Lundstrøm I, Gustin I, Mainland I, Potekhina I, Muntoni IM, Cheng J, Stenderup J, Ma J, Gibson J, Peets J, Gustafsson J, Iversen KH, Simpson L, Strand L, Loe L, Sikora M, Florek M, Vretemark M, Redknap M, Bajka M, Pushkina T, Søvsø M, Grigoreva N, Christensen T, Kastholm O, Uldum O, Favia P, Holck P, Sten S, Arge SV, Ellingvåg S, Moiseyev V, Bogdanowicz W, Magnusson Y, Orlando L, Pentz P, Jessen MD, Pedersen A, Collard M, Bradley DG, Jørkov ML, Arneborg J, Lynnerup N, Price N, Gilbert MTP, Allentoft ME, Bill J, Sindbæk SM, Hedeager L, Kristiansen K, Nielsen R, Werge T, Willerslev E. Nature. 2020 Sep;585(7825):390-396. doi: 10.1038/s41586-020-2688-8. Epub 2020 Sep 16. PMID: 32939067.

[MartinianoPLoSGenetics2017]:*
***The population genomics of archaeological transition in west Iberia: Investigation of ancient substructure using imputation and haplotype-based methods.*** *Martiniano R, Cassidy LM, Ó'Maoldúin R, McLaughlin R, Silva NM, Manco L, Fidalgo D, Pereira T, Coelho MJ, Serra M, Burger J, Parreira R, Moran E, Valera AC, Porfirio E, Boaventura R, Silva AM, Bradley DG. PLoS Genet. 2017 Jul 27;13(7):e1006852. doi: 10.1371/journal.pgen.1006852. eCollection 2017 Jul. PMID: 28749934.

[MathiesonNature2015]:*
***Genome-wide patterns of selection in 230 ancient Eurasians.*** *Mathieson I, Lazaridis I, Rohland N, Mallick S, Patterson N, Roodenberg SA, Harney E, Stewardson K, Fernandes D, Novak M, Sirak K, Gamba C, Jones ER, Llamas B, Dryomov S, Pickrell J, Arsuaga JL, de Castro JM, Carbonell E, Gerritsen F, Khokhlov A, Kuznetsov P, Lozano M, Meller H, Mochalov O, Moiseyev V, Guerra MA, Roodenberg J, Vergès JM, Krause J, Cooper A, Alt KW, Brown D, Anthony D, Lalueza-Fox C, Haak W, Pinhasi R, Reich D. Nature. 2015 Dec 24;528(7583):499-503. doi: 10.1038/nature16152. Epub 2015 Nov 23. PMID: 26595274.

[MathiesonNature2018]:*
***The genomic history of southeastern Europe.*** *Mathieson I, Alpaslan-Roodenberg S, Posth C, Szécsényi-Nagy A, Rohland N, Mallick S, Olalde I, Broomandkhoshbacht N, Candilio F, Cheronet O, Fernandes D, Ferry M, Gamarra B, Fortes GG, Haak W, Harney E, Jones E, Keating D, Krause-Kyora B, Kucukkalipci I, Michel M, Mittnik A, Nägele K, Novak M, Oppenheimer J, Patterson N, Pfrengle S, Sirak K, Stewardson K, Vai S, Alexandrov S, Alt KW, Andreescu R, Antonović D, Ash A, Atanassova N, Bacvarov K, Gusztáv MB, Bocherens H, Bolus M, Boroneanţ A, Boyadzhiev Y, Budnik A, Burmaz J, Chohadzhiev S, Conard NJ, Cottiaux R, Čuka M, Cupillard C, Drucker DG, Elenski N, Francken M, Galabova B, Ganetsovski G, Gély B, Hajdu T, Handzhyiska V, Harvati K, Higham T, Iliev S, Janković I, Karavanić I, Kennett DJ, Komšo D, Kozak A, Labuda D, Lari M, Lazar C, Leppek M, Leshtakov K, Vetro DL, Los D, Lozanov I, Malina M, Martini F, McSweeney K, Meller H, Menđušić M, Mirea P, Moiseyev V, Petrova V, Price TD, Simalcsik A, Sineo L, Šlaus M, Slavchev V, Stanev P, Starović A, Szeniczey T, Talamo S, Teschler-Nicola M, Thevenet C, Valchev I, Valentin F, Vasilyev S, Veljanovska F, Venelinova S, Veselovskaya E, Viola B, Virag C, Zaninović J, Zäuner S, Stockhammer PW, Catalano G, Krauß R, Caramelli D, Zariņa G, Gaydarska B, Lillie M, Nikitin AG, Potekhina I, Papathanasiou A, Borić D, Bonsall C, Krause J, Pinhasi R, Reich D. Nature. 2018 Mar 8;555(7695):197-203. doi: 10.1038/nature25778. Epub 2018 Feb 21. PMID: 29466330.

[McCollScience2018]:*
***The prehistoric peopling of Southeast Asia.*** *McColl H, Racimo F, Vinner L, Demeter F, Gakuhari T, Moreno-Mayar JV, van Driem G, Gram Wilken U, Seguin-Orlando A, de la Fuente Castro C, Wasef S, Shoocongdej R, Souksavatdy V, Sayavongkhamdy T, Saidin MM, Allentoft ME, Sato T, Malaspinas AS, Aghakhanian FA, Korneliussen T, Prohaska A, Margaryan A, de Barros Damgaard P, Kaewsutthi S, Lertrit P, Nguyen TMH, Hung HC, Minh Tran T, Nghia Truong H, Nguyen GH, Shahidan S, Wiradnyana K, Matsumae H, Shigehara N, Yoneda M, Ishida H, Masuyama T, Yamada Y, Tajima A, Shibata H, Toyoda A, Hanihara T, Nakagome S, Deviese T, Bacon AM, Duringer P, Ponche JL, Shackelford L, Patole-Edoumba E, Nguyen AT, Bellina-Pryce B, Galipaud JC, Kinaston R, Buckley H, Pottier C, Rasmussen S, Higham T, Foley RA, Lahr MM, Orlando L, Sikora M, Phipps ME, Oota H, Higham C, Lambert DM, Willerslev E. Science. 2018 Jul 6;361(6397):88-92. doi: 10.1126/science.aat3628. PMID: 29976827.

[MeyerScience2012]:*
***A high-coverage genome sequence from an archaic Denisovan individual.*** *Meyer M, Kircher M, Gansauge MT, Li H, Racimo F, Mallick S, Schraiber JG, Jay F, Prüfer K, de Filippo C, Sudmant PH, Alkan C, Fu Q, Do R, Rohland N, Tandon A, Siebauer M, Green RE, Bryc K, Briggs AW, Stenzel U, Dabney J, Shendure J, Kitzman J, Hammer MF, Shunkov MV, Derevianko AP, Patterson N, Andrés AM, Eichler EE, Slatkin M, Reich D, Kelso J, Pääbo S. Science. 2012 Oct 12;338(6104):222-6. doi: 10.1126/science.1224344. Epub 2012 Aug 30. PMID: 22936568.

[MittnikNatureCommunications2018]:*
***The genetic prehistory of the Baltic Sea region.*** *Mittnik A, Wang CC, Pfrengle S, Daubaras M, Zariņa G, Hallgren F, Allmäe R, Khartanovich V, Moiseyev V, Tõrv M, Furtwängler A, Andrades Valtueña A, Feldman M, Economou C, Oinonen M, Vasks A, Balanovska E, Reich D, Jankauskas R, Haak W, Schiffels S, Krause J. Nat Commun. 2018 Jan 30;9(1):442. doi: 10.1038/s41467-018-02825-9. Erratum in: Nat Commun. 2018 Apr 11;9(1):1494. PMID: 29382937.

[MittnikScience2019]:*
***Kinship-based social inequality in Bronze Age Europe.*** *Mittnik A, Massy K, Knipper C, Wittenborn F, Friedrich R, Pfrengle S, Burri M, Carlichi-Witjes N, Deeg H, Furtwängler A, Harbeck M, von Heyking K, Kociumaka C, Kucukkalipci I, Lindauer S, Metz S, Staskiewicz A, Thiel A, Wahl J, Haak W, Pernicka E, Schiffels S, Stockhammer PW, Krause J. Science. 2019 Nov 8;366(6466):731-734. doi: 10.1126/science.aax6219. Epub 2019 Oct 10. PMID: 31601705.

[MondalNatureGenetics2016]:*
***Genomic analysis of Andamanese provides insights into ancient human migration into Asia and adaptation.*** *Mondal M, Casals F, Xu T, Dall'Olio GM, Pybus M, Netea MG, Comas D, Laayouni H, Li Q, Majumder PP, Bertranpetit J. Nat Genet. 2016 Sep;48(9):1066-70. doi: 10.1038/ng.3621. Epub 2016 Jul 25. PMID: 27455350.

[MorenoMayarNature2017]:*
***Terminal Pleistocene Alaskan genome reveals first founding population of Native Americans.*** *Moreno-Mayar JV, Potter BA, Vinner L, Steinrücken M, Rasmussen S, Terhorst J, Kamm JA, Albrechtsen A, Malaspinas AS, Sikora M, Reuther JD, Irish JD, Malhi RS, Orlando L, Song YS, Nielsen R, Meltzer DJ, Willerslev E. Nature. 2018 Jan 11;553(7687):203-207. doi: 10.1038/nature25173. Epub 2018 Jan 3. PMID: 29323294.

[MorenoMayarScience2018]:*
***Early human dispersals within the Americas.*** *Moreno-Mayar JV, Vinner L, de Barros Damgaard P, de la Fuente C, Chan J, Spence JP, Allentoft ME, Vimala T, Racimo F, Pinotti T, Rasmussen S, Margaryan A, Iraeta Orbegozo M, Mylopotamitaki D, Wooller M, Bataille C, Becerra-Valdivia L, Chivall D, Comeskey D, Devièse T, Grayson DK, George L, Harry H, Alexandersen V, Primeau C, Erlandson J, Rodrigues-Carvalho C, Reis S, Bastos MQR, Cybulski J, Vullo C, Morello F, Vilar M, Wells S, Gregersen K, Hansen KL, Lynnerup N, Mirazón Lahr M, Kjær K, Strauss A, Alfonso-Durruty M, Salas A, Schroeder H, Higham T, Malhi RS, Rasic JT, Souza L, Santos FR, Malaspinas AS, Sikora M, Nielsen R, Song YS, Meltzer DJ, Willerslev E. Science. 2018 Dec 7;362(6419). pii: eaav2621. doi: 10.1126/science.aav2621. Epub 2018 Nov 8. PMID: 30409807.

[NagelePosthScience2020]:*
***Genomic insights into the early peopling of the Caribbean.*** *Nägele K, Posth C, Iraeta Orbegozo M, Chinique de Armas Y, Hernández Godoy ST, González Herrera UM, Nieves-Colón MA, Sandoval-Velasco M, Mylopotamitaki D, Radzeviciute R, Laffoon J, Pestle WJ, Ramos-Madrigal J, Lamnidis TC, Schaffer WC, Carr RS, Day JS, Arredondo Antúnez C, Rangel Rivero A, Martínez-Fuentes AJ, Crespo-Torres E, Roksandic I, Stone AC, Lalueza-Fox C, Hoogland M, Roksandic M, Hofman CL, Krause J, Schroeder H. Science. 2020 Jul 24;369(6502):456-460. doi: 10.1126/science.aba8697. Epub 2020 Jun 4. PMID: 32499399.

[NakatsukaCell2020]:*
***A Paleogenomic Reconstruction of the Deep Population History of the Andes.*** *Nakatsuka N, Lazaridis I, Barbieri C, Skoglund P, Rohland N, Mallick S, Posth C, Harkins-Kinkaid K, Ferry M, Harney É, Michel M, Stewardson K, Novak-Forst J, Capriles JM, Durruty MA, Álvarez KA, Beresford-Jones D, Burger R, Cadwallader L, Fujita R, Isla J, Lau G, Aguirre CL, LeBlanc S, Maldonado SC, Meddens F, Messineo PG, Culleton BJ, Harper TK, Quilter J, Politis G, Rademaker K, Reindel M, Rivera M, Salazar L, Sandoval JR, Santoro CM, Scheifler N, Standen V, Barreto MI, Espinoza IF, Tomasto-Cagigao E, Valverde G, Kennett DJ, Cooper A, Krause J, Haak W, Llamas B, Reich D, Fehren-Schmitz L. Cell. 2020 May 28;181(5):1131-1145.e21. doi: 10.1016/j.cell.2020.04.015. Epub 2020 May 7. PMID: 32386546; PMCID: PMC7304944.

[NarasimhanPattersonScience2019]:*
***The formation of human populations in South and Central Asia.*** *Narasimhan VM, Patterson N, Moorjani P, Rohland N, Bernardos R, Mallick S, Lazaridis I, Nakatsuka N, Olalde I, Lipson M, Kim AM, Olivieri LM, Coppa A, Vidale M, Mallory J, Moiseyev V, Kitov E, Monge J, Adamski N, Alex N, Broomandkhoshbacht N, Candilio F, Callan K, Cheronet O, Culleton BJ, Ferry M, Fernandes D, Freilich S, Gamarra B, Gaudio D, Hajdinjak M, Harney É, Harper TK, Keating D, Lawson AM, Mah M, Mandl K, Michel M, Novak M, Oppenheimer J, Rai N, Sirak K, Slon V, Stewardson K, Zalzala F, Zhang Z, Akhatov G, Bagashev AN, Bagnera A, Baitanayev B, Bendezu-Sarmiento J, Bissembaev AA, Bonora GL, Chargynov TT, Chikisheva T, Dashkovskiy PK, Derevianko A, Dobeš M, Douka K, Dubova N, Duisengali MN, Enshin D, Epimakhov A, Fribus AV, Fuller D, Goryachev A, Gromov A, Grushin SP, Hanks B, Judd M, Kazizov E, Khokhlov A, Krygin AP, Kupriyanova E, Kuznetsov P, Luiselli D, Maksudov F, Mamedov AM, Mamirov TB, Meiklejohn C, Merrett DC, Micheli R, Mochalov O, Mustafokulov S, Nayak A, Pettener D, Potts R, Razhev D, Rykun M, Sarno S, Savenkova TM, Sikhymbaeva K, Slepchenko SM, Soltobaev OA, Stepanova N, Svyatko S, Tabaldiev K, Teschler-Nicola M, Tishkin AA, Tkachev VV, Vasilyev S, Velemínský P, Voyakin D, Yermolayeva A, Zahir M, Zubkov VS, Zubova A, Shinde VS, Lalueza-Fox C, Meyer M, Anthony D, Boivin N, Thangaraj K, Kennett DJ, Frachetti M, Pinhasi R, Reich D. Science. 2019 Sep 6;365(6457). pii: eaat7487. doi: 10.1126/science.aat7487. PMID: 31488661.

[Nieves-ColónMolecularBiologyandEvolution2020]:*
***Ancient DNA Reconstructs the Genetic Legacies of Precontact Puerto Rico Communities*** *Nieves-Colón MA, Pestle WJ, Reynolds AW, Llamas B, de la Fuente C, Fowler K, Skerry KM, Crespo-Torres E, Bustamante CD, Stone AC. Mol Biol Evol. 2020 Mar 1;37(3):611-626. doi: 10.1093/molbev/msz267. PMID: 31710665.

[NikitinScientificReports2019]:*
***Interactions between earliest Linearbandkeramik farmers and central European hunter gatherers at the dawn of European Neolithization.*** *Nikitin AG, Stadler P, Kotova N, Teschler-Nicola M, Price TD, Hoover J, Kennett DJ, Lazaridis I, Rohland N, Lipson M, Reich D. Sci Rep. 2019 Dec 20;9(1):19544. doi: 10.1038/s41598-019-56029-2. PMID: 31863024.

[NingCurrentBiology2019]:*
***Ancient Genomes Reveal Yamnaya-Related Ancestry and a Potential Source of Indo-European Speakers in Iron Age Tianshan.*** *Ning C, Wang CC, Gao S, Yang Y, Zhang X, Wu X, Zhang F, Nie Z, Tang Y, Robbeets M, Ma J, Krause J, Cui Y. Curr Biol. 2019 Aug 5;29(15):2526-2532.e4. doi: 10.1016/j.cub.2019.06.044. Epub 2019 Jul 25. PMID: 31353181.

[NingNatureCommunications2020]:*
***Ancient genomes from northern China suggest links between subsistence changes and human migration.****Ning C, Li T, Wang K, Zhang F, Li T, Wu X, Gao S, Zhang Q, Zhang H, Hudson MJ, Dong G, Wu S, Fang Y, Liu C, Feng C, Li W, Han T, Li R, Wei J, Zhu Y, Zhou Y, Wang CC, Fan S, Xiong Z, Sun Z, Ye M, Sun L, Wu X, Liang F, Cao Y, Wei X, Zhu H, Zhou H, Krause J, Robbeets M, Jeong C, Cui Y. Nat Commun. 2020 Jun 1;11(1):2700. doi: 10.1038/s41467-020-16557-2. PMID: 32483115; PMCID: PMC7264253.

[OlaldeMBE2015]:*
***A Common Genetic Origin for Early Farmers from Mediterranean Cardial and Central European LBK Cultures.*** *Olalde I, Schroeder H, Sandoval-Velasco M, Vinner L, Lobón I, Ramirez O, Civit S, García Borja P, Salazar-García DC, Talamo S, María Fullola J, Xavier Oms F, Pedro M, Martínez P, Sanz M, Daura J, Zilhão J, Marquès-Bonet T, Gilbert MT, Lalueza-Fox C. Mol Biol Evol. 2015 Dec;32(12):3132-42. doi: 10.1093/molbev/msv181. Epub 2015 Sep 2. PMID: 26337550.

[OlaldeNature2014]:*
***Derived immune and ancestral pigmentation alleles in a 7,000-year-old Mesolithic European.*** *Olalde I, Allentoft ME, Sánchez-Quinto F, Santpere G, Chiang CW, DeGiorgio M, Prado-Martinez J, Rodríguez JA, Rasmussen S, Quilez J, Ramírez O, Marigorta UM, Fernández-Callejo M, Prada ME, Encinas JM, Nielsen R, Netea MG, Novembre J, Sturm RA, Sabeti P, Marquès-Bonet T, Navarro A, Willerslev E, Lalueza-Fox C. Nature. 2014 Mar 13;507(7491):225-8. doi: 10.1038/nature12960. Epub 2014 Jan 26. PMID: 24463515.

[OlaldeNature2018]:*
***The Beaker phenomenon and the genomic transformation of northwest Europe.*** *Olalde I, Brace S, Allentoft ME, Armit I, Kristiansen K, Booth T, Rohland N, Mallick S, Szécsényi-Nagy A, Mittnik A, Altena E, Lipson M, Lazaridis I, Harper TK, Patterson N, Broomandkhoshbacht N, Diekmann Y, Faltyskova Z, Fernandes D, Ferry M, Harney E, de Knijff P, Michel M, Oppenheimer J, Stewardson K, Barclay A, Alt KW, Liesau C, Ríos P, Blasco C, Miguel JV, García RM, Fernández AA, Bánffy E, Bernabò-Brea M, Billoin D, Bonsall C, Bonsall L, Allen T, Büster L, Carver S, Navarro LC, Craig OE, Cook GT, Cunliffe B, Denaire A, Dinwiddy KE, Dodwell N, Ernée M, Evans C, Kuchařík M, Farré JF, Fowler C, Gazenbeek M, Pena RG, Haber-Uriarte M, Haduch E, Hey G, Jowett N, Knowles T, Massy K, Pfrengle S, Lefranc P, Lemercier O, Lefebvre A, Martínez CH, Olmo VG, Ramírez AB, Maurandi JL, Majó T, McKinley JI, McSweeney K, Mende BG, Modi A, KulcsáR G, Kiss V, Czene A, Patay R, Endrődi A, Köhler K, Hajdu T, Szeniczey T, Dani J, Bernert Z, Hoole M, Cheronet O, Keating D, Velemínský P, Dobeš M, Candilio F, Brown F, Fernández RF, Herrero-Corral AM, Tusa S, Carnieri E, Lentini L, Valenti A, Zanini A, Waddington C, Delibes G, Guerra-Doce E, Neil B, Brittain M, Luke M, Mortimer R, Desideri J, Besse M, Brücken G, Furmanek M, Hałuszko A, Mackiewicz M, Rapiński A, Leach S, Soriano I, Lillios KT, Cardoso JL, Pearson MP, Włodarczak P, Price TD, Prieto P, Rey PJ, Risch R, Rojo Guerra MA, Schmitt A, Serralongue J, Silva AM, Smrčka V, Vergnaud L, Zilhão J, Caramelli D, Higham T, Thomas MG, Kennett DJ, Fokkens H, Heyd V, Sheridan A, Sjögren KG, Stockhammer PW, Krause J, Pinhasi R, Haak W, Barnes I, Lalueza-Fox C, Reich D. Nature. 2018 Mar 8;555(7695):190-196. doi: 10.1038/nature25738. Epub 2018 Feb 21. Erratum in: Nature. 2018 Mar 21;555(7697):543. PMID: 29466337.

[OlaldeScience2019]:*
***The genomic history of the Iberian Peninsula over the past 8000 years.*** *Output encoded/decoded text Olalde I, Mallick S, Patterson N, Rohland N, Villalba-Mouco V, Silva M, Dulias K, Edwards CJ, Gandini F, Pala M, Soares P, Ferrando-Bernal M, Adamski N, Broomandkhoshbacht N, Cheronet O, Culleton BJ, Fernandes D, Lawson AM, Mah M, Oppenheimer J, Stewardson K, Zhang Z, Jiménez Arenas JM, Toro Moyano IJ, Salazar-García DC, Castanyer P, Santos M, Tremoleda J, Lozano M, García Borja P, Fernández-Eraso J, Mujika-Alustiza JA, Barroso C, Bermúdez FJ, Viguera Mínguez E, Burch J, Coromina N, Vivó D, Cebrià A, Fullola JM, García-Puchol O, Morales JI, Oms FX, Majó T, Vergès JM, Díaz-Carvajal A, Ollich-Castanyer I, López-Cachero FJ, Silva AM, Alonso-Fernández C, Delibes de Castro G, Jiménez Echevarría J, Moreno-Márquez A, Pascual Berlanga G, Ramos-García P, Ramos-Muñoz J, Vijande Vila E, Aguilella Arzo G, Esparza Arroyo Á, Lillios KT, Mack J, Velasco-Vázquez J, Waterman A, Benítez de Lugo Enrich L, Benito Sánchez M, Agustí B, Codina F, de Prado G, Estalrrich A, Fernández Flores Á, Finlayson C, Finlayson G, Finlayson S, Giles-Guzmán F, Rosas A, Barciela González V, García Atiénzar G, Hernández Pérez MS, Llanos A, Carrión Marco Y, Collado Beneyto I, López-Serrano D, Sanz Tormo M, Valera AC, Blasco C, Liesau C, Ríos P, Daura J, de Pedro Michó MJ, Diez-Castillo AA, Flores Fernández R, Francès Farré J, Garrido-Pena R, Gonçalves VS, Guerra-Doce E, Herrero-Corral AM, Juan-Cabanilles J, López-Reyes D, McClure SB, Merino Pérez M, Oliver Foix A, Sanz Borràs M, Sousa AC, Vidal Encinas JM, Kennett DJ, Richards MB, Werner Alt K, Haak W, Pinhasi R, Lalueza-Fox C, Reich D. Science. 2019 Mar 15;363(6432):1230-1234. doi: 10.1126/science.aav4040. PMID: 30872528.

[OmrakCurrentBiology2016]:*
***Genomic Evidence Establishes Anatolia as the Source of the European Neolithic Gene Pool.*** *Omrak A, Günther T, Valdiosera C, Svensson EM, Malmström H, Kiesewetter H, Aylward W, Storå J, Jakobsson M, Götherström A. Curr Biol. 2016 Jan 25;26(2):270-275. doi: 10.1016/j.cub.2015.12.019. Epub 2015 Dec 31. PMID: 26748850.

[PattersonGenetics2012]:*
***Ancient admixture in human history.*** *Patterson N, Moorjani P, Luo Y, Mallick S, Rohland N, Zhan Y, Genschoreck T, Webster T, Reich D. Genetics. 2012 Nov;192(3):1065-93. doi: 10.1534/genetics.112.145037. Epub 2012 Sep 7. PMID: 22960212.

[Pickrellnaturecommunications2012]:*
***The genetic prehistory of southern Africa.*** *Pickrell JK, Patterson N, Barbieri C, Berthold F, Gerlach L, Güldemann T, Kure B, Mpoloka SW, Nakagawa H, Naumann C, Lipson M, Loh PR, Lachance J, Mountain J, Bustamante CD, Berger B, Tishkoff SA, Henn BM, Stoneking M, Reich D, Pakendorf B. Nat Commun. 2012;3:1143. doi: 10.1038/ncomms2140. PMID: 23072811.

[PosthNakatsukaCell2018]:*
***Reconstructing the Deep Population History of Central and South America.*** *Posth C, Nakatsuka N, Lazaridis I, Skoglund P, Mallick S, Lamnidis TC, Rohland N, Nägele K, Adamski N, Bertolini E, Broomandkhoshbacht N, Cooper A, Culleton BJ, Ferraz T, Ferry M, Furtwängler A, Haak W, Harkins K, Harper TK, Hünemeier T, Lawson AM, Llamas B, Michel M, Nelson E, Oppenheimer J, Patterson N, Schiffels S, Sedig J, Stewardson K, Talamo S, Wang CC, Hublin JJ, Hubbe M, Harvati K, Nuevo Delaunay A, Beier J, Francken M, Kaulicke P, Reyes-Centeno H, Rademaker K, Trask WR, Robinson M, Gutierrez SM, Prufer KM, Salazar-García DC, Chim EN, Müller Plumm Gomes L, Alves ML, Liryo A, Inglez M, Oliveira RE, Bernardo DV, Barioni A, Wesolowski V, Scheifler NA, Rivera MA, Plens CR, Messineo PG, Figuti L, Corach D, Scabuzzo C, Eggers S, DeBlasis P, Reindel M, Méndez C, Politis G, Tomasto-Cagigao E, Kennett DJ, Strauss A, Fehren-Schmitz L, Krause J, Reich D. Cell. 2018 Nov 15;175(5):1185-1197.e22. doi: 10.1016/j.cell.2018.10.027. Epub 2018 Nov 8. PMID: 30415837.

[PosthNatureEcologyEvolution2018]:*
***Language continuity despite population replacement in Remote Oceania.*** *Posth C, Nägele K, Colleran H, Valentin F, Bedford S, Kami KW, Shing R, Buckley H, Kinaston R, Walworth M, Clark GR, Reepmeyer C, Flexner J, Maric T, Moser J, Gresky J, Kiko L, Robson KJ, Auckland K, Oppenheimer SJ, Hill AVS, Mentzer AJ, Zech J, Petchey F, Roberts P, Jeong C, Gray RD, Krause J, Powell A. Nat Ecol Evol. 2018 Apr;2(4):731-740. doi: 10.1038/s41559-018-0498-2. Epub 2018 Feb 27. PMID: 29487365.

[PrendergastLipsonSawchukScience2019]:*
***Ancient DNA reveals a multistep spread of the first herders into sub-Saharan Africa.*** *Prendergast ME, Lipson M, Sawchuk EA, Olalde I, Ogola CA, Rohland N, Sirak KA, Adamski N, Bernardos R, Broomandkhoshbacht N, Callan K, Culleton BJ, Eccles L, Harper TK, Lawson AM, Mah M, Oppenheimer J, Stewardson K, Zalzala F, Ambrose SH, Ayodo G, Gates HL Jr, Gidna AO, Katongo M, Kwekason A, Mabulla AZP, Mudenda GS, Ndiema EK, Nelson C, Robertshaw P, Kennett DJ, Manthi FK, Reich D. Science. 2019 Jul 5;365(6448). pii: eaaw6275. doi: 10.1126/science.aaw6275. Epub 2019 May 30. PMID: 31147405.

[Pruefer2017]:*
***A high-coverage Neandertal genome from Vindija Cave in Croatia.*** *Prüfer K, de Filippo C, Grote S, Mafessoni F, Korlević P, Hajdinjak M, Vernot B, Skov L, Hsieh P, Peyrégne S, Reher D, Hopfe C, Nagel S, Maricic T, Fu Q, Theunert C, Rogers R, Skoglund P, Chintalapati M, Dannemann M, Nelson BJ, Key FM, Rudan P, Kućan Ž, Gušić I, Golovanova LV, Doronichev VB, Patterson N, Reich D, Eichler EE, Slatkin M, Schierup MH, Andrés AM, Kelso J, Meyer M, Pääbo S. Science. 2017 Nov 3;358(6363):655-658. doi: 10.1126/science.aao1887. Epub 2017 Oct 5. PMID: 28982794.

[PrueferNature2013]:*
***The complete genome sequence of a Neanderthal from the Altai Mountains.*** *Prüfer K, Racimo F, Patterson N, Jay F, Sankararaman S, Sawyer S, Heinze A, Renaud G, Sudmant PH, de Filippo C, Li H, Mallick S, Dannemann M, Fu Q, Kircher M, Kuhlwilm M, Lachmann M, Meyer M, Ongyerth M, Siebauer M, Theunert C, Tandon A, Moorjani P, Pickrell J, Mullikin JC, Vohr SH, Green RE, Hellmann I, Johnson PL, Blanche H, Cann H, Kitzman JO, Shendure J, Eichler EE, Lein ES, Bakken TE, Golovanova LV, Doronichev VB, Shunkov MV, Derevianko AP, Viola B, Slatkin M, Reich D, Kelso J, Pääbo S. Nature. 2014 Jan 2;505(7481):43-9. doi: 10.1038/nature12886. Epub 2013 Dec 18. PMID: 24352235.

[RaghavanNature2013]:*
***Upper Palaeolithic Siberian genome reveals dual ancestry of Native Americans.*** *Raghavan M, Skoglund P, Graf KE, Metspalu M, Albrechtsen A, Moltke I, Rasmussen S, Stafford TW Jr, Orlando L, Metspalu E, Karmin M, Tambets K, Rootsi S, Mägi R, Campos PF, Balanovska E, Balanovsky O, Khusnutdinova E, Litvinov S, Osipova LP, Fedorova SA, Voevoda MI, DeGiorgio M, Sicheritz-Ponten T, Brunak S, Demeshchenko S, Kivisild T, Villems R, Nielsen R, Jakobsson M, Willerslev E.

[RaghavanNature2014]:*
***Upper Palaeolithic Siberian genome reveals dual ancestry of Native Americans.*** *Raghavan M, Skoglund P, Graf KE, Metspalu M, Albrechtsen A, Moltke I, Rasmussen S, Stafford TW Jr, Orlando L, Metspalu E, Karmin M, Tambets K, Rootsi S, Mägi R, Campos PF, Balanovska E, Balanovsky O, Khusnutdinova E, Litvinov S, Osipova LP, Fedorova SA, Voevoda MI, DeGiorgio M, Sicheritz-Ponten T, Brunak S, Demeshchenko S, Kivisild T, Villems R, Nielsen R, Jakobsson M, Willerslev E. Nature. 2014 Jan 2;505(7481):87-91. doi: 10.1038/nature12736. Epub 2013 Nov 20. PMID: 24256729.

[RaghavanScience2014]:*
***The genetic prehistory of the New World Arctic.*** *Raghavan M, DeGiorgio M, Albrechtsen A, Moltke I, Skoglund P, Korneliussen TS, Grønnow B, Appelt M, Gulløv HC, Friesen TM, Fitzhugh W, Malmström H, Rasmussen S, Olsen J, Melchior L, Fuller BT, Fahrni SM, Stafford T Jr, Grimes V, Renouf MA, Cybulski J, Lynnerup N, Lahr MM, Britton K, Knecht R, Arneborg J, Metspalu M, Cornejo OE, Malaspinas AS, Wang Y, Rasmussen M, Raghavan V, Hansen TV, Khusnutdinova E, Pierre T, Dneprovsky K, Andreasen C, Lange H, Hayes MG, Coltrain J, Spitsyn VA, Götherström A, Orlando L, Kivisild T, Villems R, Crawford MH, Nielsen FC, Dissing J, Heinemeier J, Meldgaard M, Bustamante C, O'Rourke DH, Jakobsson M, Gilbert MT, Nielsen R, Willerslev E. Science. 2014 Aug 29;345(6200):1255832. doi: 10.1126/science.1255832. PMID: 25170159.

[RaghavanScience2015]:*
***POPULATION GENETICS. Genomic evidence for the Pleistocene and recent population history of Native Americans.*** *Raghavan M, Steinrücken M, Harris K, Schiffels S, Rasmussen S, DeGiorgio M, Albrechtsen A, Valdiosera C, Ávila-Arcos MC, Malaspinas AS, Eriksson A, Moltke I, Metspalu M, Homburger JR, Wall J, Cornejo OE, Moreno-Mayar JV, Korneliussen TS, Pierre T, Rasmussen M, Campos PF, de Barros Damgaard P, Allentoft ME, Lindo J, Metspalu E, Rodríguez-Varela R, Mansilla J, Henrickson C, Seguin-Orlando A, Malmström H, Stafford T Jr, Shringarpure SS, Moreno-Estrada A, Karmin M, Tambets K, Bergström A, Xue Y, Warmuth V, Friend AD, Singarayer J, Valdes P, Balloux F, Leboreiro I, Vera JL, Rangel-Villalobos H, Pettener D, Luiselli D, Davis LG, Heyer E, Zollikofer CPE, Ponce de León MS, Smith CI, Grimes V, Pike KA, Deal M, Fuller BT, Arriaza B, Standen V, Luz MF, Ricaut F, Guidon N, Osipova L, Voevoda MI, Posukh OL, Balanovsky O, Lavryashina M, Bogunov Y, Khusnutdinova E, Gubina M, Balanovska E, Fedorova S, Litvinov S, Malyarchuk B, Derenko M, Mosher MJ, Archer D, Cybulski J, Petzelt B, Mitchell J, Worl R, Norman PJ, Parham P, Kemp BM, Kivisild T, Tyler-Smith C, Sandhu MS, Crawford M, Villems R, Smith DG, Waters MR, Goebel T, Johnson JR, Malhi RS, Jakobsson M, Meltzer DJ, Manica A, Durbin R, Bustamante CD, Song YS, Nielsen R, Willerslev E. Science. 2015 Aug 21;349(6250):aab3884. doi: 10.1126/science.aab3884. Epub 2015 Jul 21. PMID: 26198033.

[RasmussenNature2010]:*
***Ancient human genome sequence of an extinct Palaeo-Eskimo.*** *Rasmussen M, Li Y, Lindgreen S, Pedersen JS, Albrechtsen A, Moltke I, Metspalu M, Metspalu E, Kivisild T, Gupta R, Bertalan M, Nielsen K, Gilbert MT, Wang Y, Raghavan M, Campos PF, Kamp HM, Wilson AS, Gledhill A, Tridico S, Bunce M, Lorenzen ED, Binladen J, Guo X, Zhao J, Zhang X, Zhang H, Li Z, Chen M, Orlando L, Kristiansen K, Bak M, Tommerup N, Bendixen C, Pierre TL, Grønnow B, Meldgaard M, Andreasen C, Fedorova SA, Osipova LP, Higham TF, Ramsey CB, Hansen TV, Nielsen FC, Crawford MH, Brunak S, Sicheritz-Pontén T, Villems R, Nielsen R, Krogh A, Wang J, Willerslev E. Nature. 2010 Feb 11;463(7282):757-62. doi: 10.1038/nature08835. PMID: 20148029.

[RasmussenNature2014]:*
***The genome of a Late Pleistocene human from a Clovis burial site in western Montana.*** *Rasmussen M, Anzick SL, Waters MR, Skoglund P, DeGiorgio M, Stafford TW Jr, Rasmussen S, Moltke I, Albrechtsen A, Doyle SM, Poznik GD, Gudmundsdottir V, Yadav R, Malaspinas AS, White SS 5th, Allentoft ME, Cornejo OE, Tambets K, Eriksson A, Heintzman PD, Karmin M, Korneliussen TS, Meltzer DJ, Pierre TL, Stenderup J, Saag L, Warmuth VM, Lopes MC, Malhi RS, Brunak S, Sicheritz-Ponten T, Barnes I, Collins M, Orlando L, Balloux F, Manica A, Gupta R, Metspalu M, Bustamante CD, Jakobsson M, Nielsen R, Willerslev E. Nature. 2014 Feb 13;506(7487):225-9. doi: 10.1038/nature13025. PMID: 24522598.

[RasmussenNature2015]:*
***The ancestry and affiliations of Kennewick Man.*** *Rasmussen M, Sikora M, Albrechtsen A, Korneliussen TS, Moreno-Mayar JV, Poznik GD, Zollikofer CPE, de León MP, Allentoft ME, Moltke I, Jónsson H, Valdiosera C, Malhi RS, Orlando L, Bustamante CD, Stafford TW Jr, Meltzer DJ, Nielsen R, Willerslev E. Nature. 2015 Jul 23;523(7561):455-458. doi: 10.1038/nature14625.

[RodriguezVarelaCurrentBiology2017]:*
***Genomic Analyses of Pre-European Conquest Human Remains from the Canary Islands Reveal Close Affinity to Modern North Africans.*** *Rodríguez-Varela R, Günther T, Krzewińska M, Storå J, Gillingwater TH, MacCallum M, Arsuaga JL, Dobney K, Valdiosera C, Jakobsson M, Götherström A, Girdland-Flink L. Curr Biol. 2017 Nov 6;27(21):3396-3402.e5. doi: 10.1016/j.cub.2017.09.059. Epub 2017 Oct 26. Erratum in: Curr Biol. 2018 May 21;28(10 ):1677-1679. PMID: 29107554.

[RivollatScienceAdvance2020]:*
***Ancient genome-wide DNA from France highlights the complexity of interactions between Mesolithic hunter-gatherers and Neolithic farmers.*** *Rivollat M, Jeong C, Schiffels S, Küçükkalıpçı İ, Pemonge MH, Rohrlach AB, Alt KW, Binder D, Friederich S, Ghesquière E, Gronenborn D, Laporte L, Lefranc P, Meller H, Réveillas H, Rosenstock E, Rottier S, Scarre C, Soler L, Wahl J, Krause J, Deguilloux MF, Haak W. Sci Adv. 2020 May 29;6(22):eaaz5344. doi: 10.1126/sciadv.aaz5344. PMID: 32523989; PMCID: PMC7259947.

[SaagCurrentBiology2017]:*
***Extensive Farming in Estonia Started through a Sex-Biased Migration from the Steppe.****Saag L, Varul L, Scheib CL, Stenderup J, Allentoft ME, Saag L, Pagani L, Reidla M, Tambets K, Metspalu E, Kriiska A, Willerslev E, Kivisild T, Metspalu M.Curr Biol. 2017 Jul 24;27(14):2185-2193.e6. doi: 10.1016/j.cub.2017.06.022. Epub 2017 Jul 14. PMID: 28712569.

[SaagCurrentBiology2019]:*
***The Arrival of Siberian Ancestry Connecting the Eastern Baltic to Uralic Speakers further East.*** *Saag L, Laneman M, Varul L, Malve M, Valk H, Razzak MA, Shirobokov IG, Khartanovich VI, Mikhaylova ER, Kushniarevich A, Scheib CL, Solnik A, Reisberg T, Parik J, Saag L, Metspalu E, Rootsi S, Montinaro F, Remm M, Mägi R, D'Atanasio E, Crema ER, D'Atanasio E, Crema ER, Díez-Del-Molino D, Thomas MG, Kriiska A, Kivisild T, Villems R, Lang V, Metspalu M, Tambets K. Curr Biol. 2019 May 20;29(10):1701-1711.e16. doi: 10.1016/j.cub.2019.04.026. Epub 2019 May 9. PMID: 31080083.

[ScheibScience2018]:*
***Ancient human parallel lineages within North America contributed to a coastal expansion.*** *Scheib CL, Li H, Desai T, Link V, Kendall C, Dewar G, Griffith PW, Mörseburg A, Johnson JR, Potter A, Kerr SL, Endicott P, Lindo J, Haber M, Xue Y, Tyler-Smith C, Sandhu MS, Lorenz JG, Randall TD, Faltyskova Z, Pagani L, Danecek P, O'Connell TC, Martz P, Boraas AS, Byrd BF, Leventhal A, Cambra R, Williamson R, Lesage L, Holguin B, Ygnacio-De Soto E, Rosas J, Metspalu M, Stock JT, Manica A, Scally A, Wegmann D, Malhi RS, Kivisild T. Science. 2018 Jun 1;360(6392):1024-1027. doi: 10.1126/science.aar6851. PMID: 29853687.

[SanchezQuintoPNAS2019]:*
***Megalithic tombs in western and northern Neolithic Europe were linked to a kindred society.*** *Sánchez-Quinto F, Malmström H, Fraser M, Girdland-Flink L, Svensson EM, Simões LG, George R, Hollfelder N, Burenhult G, Noble G, Britton K, Talamo S, Curtis N, Brzobohata H, Sumberova R, Götherström A, Storå J, Jakobsson M. Proc Natl Acad Sci U S A. 2019 May 7;116(19):9469-9474. doi: 10.1073/pnas.1818037116. Epub 2019 Apr 15. PMID: 30988179.

[ScheunemannNatureCommunications2017]:*
***Ancient Egyptian mummy genomes suggest an increase of Sub-Saharan African ancestry in post-Roman periods.*** *Schuenemann VJ, Peltzer A, Welte B, van Pelt WP, Molak M, Wang CC, Furtwängler A, Urban C, Reiter E, Nieselt K, Teßmann B, Francken M, Harvati K, Haak W, Schiffels S, Krause J. Nat Commun. 2017 May 30;8:15694. doi: 10.1038/ncomms15694. PMID: 28556824.

[SchiffelsNatureCommunications2016]:*
***Iron Age and Anglo-Saxon genomes from East England reveal British migration history.*** *Schiffels S, Haak W, Paajanen P, Llamas B, Popescu E, Loe L, Clarke R, Lyons A, Mortimer R, Sayer D, Tyler-Smith C, Cooper A, Durbin R. Nat Commun. 2016 Jan 19;7:10408. doi: 10.1038/ncomms10408. PMID: 26783965.

[SchlebuschScience2017]:*
***Southern African ancient genomes estimate modern human divergence to 350,000 to 260,000 years ago.*** *Schlebusch CM, Malmström H, Günther T, Sjödin P, Coutinho A, Edlund H, Munters AR, Vicente M, Steyn M, Soodyall H, Lombard M, Jakobsson M. Science. 2017 Nov 3;358(6363):652-655. doi: 10.1126/science.aao6266. Epub 2017 Sep 28. PMID: 28971970.

[SchroederPNAS2018]:*
***Origins and genetic legacies of the Caribbean Taino.*** *Schroeder H, Sikora M, Gopalakrishnan S, Cassidy LM, Maisano Delser P, Sandoval Velasco M, Schraiber JG, Rasmussen S, Homburger JR, Ávila-Arcos MC, Allentoft ME, Moreno-Mayar JV, Renaud G, Gómez-Carballa A, Laffoon JE, Hopkins RJA, Higham TFG, Carr RS, Schaffer WC, Day JS, Hoogland M, Salas A, Bustamante CD, Nielsen R, Bradley DG, Hofman CL, Willerslev E. Proc Natl Acad Sci U S A. 2018 Mar 6;115(10):2341-2346. doi: 10.1073/pnas.1716839115. Epub 2018 Feb 20. PMID: 29463742.

[SchroederPNAS2019]:*
***Unraveling ancestry, kinship, and violence in a Late Neolithic mass grave.*** *Schroeder H, Margaryan A, Szmyt M, Theulot B, Wlodarczak P, Rasmussen S, Gopalakrishnan S, Szczepanek A, Konopka T, Jensen TZT, Witkowska B, Wilk S, Przybyla MM, Pospieszny L, Sjögren KG, Belka Z, Olsen J, Kristiansen K, Willerslev E, Frei KM, Sikora M, Johannsen NN, Allentoft ME. Proc Natl Acad Sci U S A. 2019 May 28;116(22):10705-10710. doi: 10.1073/pnas.1820210116. Epub 2019 May 6. PMID: 31061125.

[Seguin-OrlandoScience2014]:odarczak P, Rasmussen S, Gopalakrishnan S, Szczepanek A, Konopka T, Jensen TZT, Witkowska B, Wilk S, Przybya MM, Pospieszny*
***Paleogenomics. Genomic structure in Europeans dating back at least 36,200 years.*** *Seguin-Orlando A, Korneliussen TS, Sikora M, Malaspinas AS, Manica A, Moltke I, Albrechtsen A, Ko A, Margaryan A, Moiseyev V, Goebel T, Westaway M, Lambert D, Khartanovich V, Wall JD, Nigst PR, Foley RA, Lahr MM, Nielsen R, Orlando L, Willerslev E. Science. 2014 Nov 28;346(6213):1113-8. doi: 10.1126/science.aaa0114. Epub 2014 Nov 6. PMID: 25378462.

[ShindeNarasimhanCell2019]:*
***An Ancient Harappan Genome Lacks Ancestry from Steppe Pastoralists or Iranian Farmers.*** *Shinde V, Narasimhan VM, Rohland N, Mallick S, Mah M, Lipson M, Nakatsuka N, Adamski N, Broomandkhoshbacht N, Ferry M, Lawson AM, Michel M, Oppenheimer J, Stewardson K, Jadhav N, Kim YJ, Chatterjee M, Munshi A, Panyam A, Waghmare P, Yadav Y, Patel H, Kaushik A, Thangaraj K, Meyer M, Patterson N, Rai N, Reich D. Cell. 2019 Oct 17;179(3):729-735.e10. doi: 10.1016/j.cell.2019.08.048. Epub 2019 Sep 5. PMID: 31495572.

[SikoraNature2019]:*
***The population history of northeastern Siberia since the Pleistocene.*** *Sikora M, Pitulko VV, Sousa VC, Allentoft ME, Vinner L, Rasmussen S, Margaryan A, de Barros Damgaard P, de la Fuente C, Renaud G, Yang MA, Fu Q, Dupanloup I, Giampoudakis K, Nogués-Bravo D, Rahbek C, Kroonen G, Peyrot M, McColl H, Vasilyev SV, Veselovskaya E, Gerasimova M, Pavlova EY, Chasnyk VG, Nikolskiy PA, Gromov AV, Khartanovich VI, Moiseyev V, Grebenyuk PS, Fedorchenko AY, Lebedintsev AI, Slobodin SB, Malyarchuk BA, Martiniano R, Meldgaard M, Arppe L, Palo JU, Sundell T, Mannermaa K, Putkonen M, Alexandersen V, Primeau C, Baimukhanov N, Malhi RS, Sjögren KG, Kristiansen K, Wessman A, Sajantila A, Lahr MM, Durbin R, Nielsen R, Meltzer DJ, Excoffier L, Willerslev E. Nature. 2019 Jun;570(7760):182-188. doi: 10.1038/s41586-019-1279-z. Epub 2019 Jun 5. PMID: 31168093.

[SikoraScience2017]:*
***Ancient genomes show social and reproductive behavior of early Upper Paleolithic foragers.*** *Sikora M, Seguin-Orlando A, Sousa VC, Albrechtsen A, Korneliussen T, Ko A, Rasmussen S, Dupanloup I, Nigst PR, Bosch MD, Renaud G, Allentoft ME, Margaryan A, Vasilyev SV, Veselovskaya EV, Borutskaya SB, Deviese T, Comeskey D, Higham T, Manica A, Foley R, Meltzer DJ, Nielsen R, Excoffier L, Mirazon Lahr M, Orlando L, Willerslev E. Science. 2017 Nov 3;358(6363):659-662. doi: 10.1126/science.aao1807. Epub 2017 Oct 5.

[SiskaScienceAdvances2017]:*
***Ancient genomes show social and reproductive behavior of early Upper Paleolithic foragers.*** *Sikora M, Seguin-Orlando A, Sousa VC, Albrechtsen A, Korneliussen T, Ko A, Rasmussen S, Dupanloup I, Nigst PR, Bosch MD, Renaud G, Allentoft ME, Margaryan A, Vasilyev SV, Veselovskaya EV, Borutskaya SB, Deviese T, Comeskey D, Higham T, Manica A, Foley R, Meltzer DJ, Nielsen R, Excoffier L, Mirazon Lahr M, Orlando L, Willerslev E. Science. 2017 Nov 3;358(6363):659-662. doi: 10.1126/science.aao1807. Epub 2017 Oct 5. PMID: 28982795.

[SkoglundCell2017]:*
***Reconstructing Prehistoric African Population Structure.*** *Skoglund P, Thompson JC, Prendergast ME, Mittnik A, Sirak K, Hajdinjak M, Salie T, Rohland N, Mallick S, Peltzer A, Heinze A, Olalde I, Ferry M, Harney E, Michel M, Stewardson K, Cerezo-Román JI, Chiumia C, Crowther A, Gomani-Chindebvu E, Gidna AO, Grillo KM, Helenius IT, Hellenthal G, Helm R, Horton M, López S, Mabulla AZP, Parkington J, Shipton C, Thomas MG, Tibesasa R, Welling M, Hayes VM, Kennett DJ, Ramesar R, Meyer M, Pääbo S, Patterson N, Morris AG, Boivin N, Pinhasi R, Krause J, Reich D. Cell. 2017 Sep 21;171(1):59-71.e21. doi: 10.1016/j.cell.2017.08.049. PMID: 28938123.

[SkoglundNature2015]:*
***Genetic evidence for two founding populations of the Americas.*** *Skoglund P, Mallick S, Bortolini MC, Chennagiri N, Hünemeier T, Petzl-Erler ML, Salzano FM, Patterson N, Reich D. Nature. 2015 Sep 3;525(7567):104-8. doi: 10.1038/nature14895. Epub 2015 Jul 21. PMID: 26196601.

[SkoglundNature2016]:*
***Genomic insights into the peopling of the Southwest Pacific.*** *Skoglund P, Posth C, Sirak K, Spriggs M, Valentin F, Bedford S, Clark GR, Reepmeyer C, Petchey F, Fernandes D, Fu Q, Harney E, Lipson M, Mallick S, Novak M, Rohland N, Stewardson K, Abdullah S, Cox MP, Friedlaender FR, Friedlaender JS, Kivisild T, Koki G, Kusuma P, Merriwether DA, Ricaut FX, Wee JT, Patterson N, Krause J, Pinhasi R, Reich D. Nature. 2016 Oct 27;538(7626):510-513. doi: 10.1038/nature19844. Epub 2016 Oct 3. PMID: 27698418.

[SkoglundScience2014]:*
***Genomic diversity and admixture differs for Stone-Age Scandinavian foragers and farmers.*** *Skoglund P, Malmström H, Omrak A, Raghavan M, Valdiosera C, Günther T, Hall P, Tambets K, Parik J, Sjögren KG, Apel J, Willerslev E, Storå J, Götherström A, Jakobsson M. Science. 2014 May 16;344(6185):747-50. doi: 10.1126/science.1253448. Epub 2014 Apr 24. PMID: 24762536.

[SkourtaniontiCell2020]:*
***Genomic History of Neolithic to Bronze Age Anatolia, Northern Levant, and Southern Caucasus.*** *Skourtanioti E, Erdal YS, Frangipane M, Balossi Restelli F, Yener KA, Pinnock F, Matthiae P, Özbal R, Schoop UD, Guliyev F, Akhundov T, Lyonnet B, Hammer EL, Nugent SE, Burri M, Neumann GU, Penske S, Ingman T, Akar M, Shafiq R, Palumbi G, Eisenmann S, D'Andrea M, Rohrlach AB, Warinner C, Jeong C, Stockhammer PW, Haak W, Krause J. Cell. 2020 May 28;181(5):1158-1175.e28. doi: 10.1016/j.cell.2020.04.044. PMID: 32470401.

[SlonNature2018]:*
***The genome of the offspring of a Neanderthal mother and a Denisovan father.*** *Slon V, Mafessoni F, Vernot B, de Filippo C, Grote S, Viola B, Hajdinjak M, Peyrégne S, Nagel S, Brown S, Douka K, Higham T, Kozlikin MB, Shunkov MV, Derevianko AP, Kelso J, Meyer M, Prüfer K, Pääbo S.Nature. 2018 Sep;561(7721):113-116. doi: 10.1038/s41586-018-0455-x. Epub 2018 Aug 22. PMID: 30135579.

[SullivanScienceAdvances2018]:*
***Ancient genome-wide analyses infer kinship structure in an Early Medieval Alemannic graveyard.*** *O'Sullivan N, Posth C, Coia V, Schuenemann VJ, Price TD, Wahl J, Pinhasi R, Zink A, Krause J, Maixner F. Sci Adv. 2018 Sep 5;4(9):eaao1262. doi: 10.1126/sciadv.aao1262. PMID: 30191172; PMCID: PMC6124919.

[TeschlerNicolaCommunicationsBiology2020]:*
***Ancient DNA reveals monozygotic newborn twins from the Upper Palaeolithic*** *Teschler-Nicola M, Fernandes D, Händel M, Einwögerer T, Simon U, Neugebauer-Maresch C, Tangl S, Heimel P, Dobsak T, Retzmann A, Prohaska T, Irrgeher J, Kennett DJ, Olalde I, Reich D, Pinhasi R. Commun Biol. 2020 Nov 6;3(1):650. doi: 10.1038/s42003-020-01372-8. PMID: 33159107; PMCID: PMC7648643.

[UnterlanderNatureCommunications2017]:*
***Ancestry and demography and descendants of Iron Age nomads of the Eurasian Steppe.*** *Unterländer M, Palstra F, Lazaridis I, Pilipenko A, Hofmanová Z, Groß M, Sell C, Blöcher J, Kirsanow K, Rohland N, Rieger B, Kaiser E, Schier W, Pozdniakov D, Khokhlov A, Georges M, Wilde S, Powell A, Heyer E, Currat M, Reich D, Samashev Z, Parzinger H, Molodin VI, Burger J. Nat Commun. 2017 Mar 3;8:14615. doi: 10.1038/ncomms14615. PMID: 28256537.

[ValdioseraPNAS2018]:*
***Four millennia of Iberian biomolecular prehistory illustrate the impact of prehistoric migrations at the far end of Eurasia.*** *Valdiosera C, Günther T, Vera-Rodríguez JC, Ureña I, Iriarte E, Rodríguez-Varela R, Simões LG, Martínez-Sánchez RM, Svensson EM, Malmström H, Rodríguez L, Bermúdez de Castro JM, Carbonell E, Alday A, Hernández Vera JA, Götherström A, Carretero JM, Arsuaga JL, Smith CI, Jakobsson M. Proc Natl Acad Sci U S A. 2018 Mar 27;115(13):3428-3433. doi: 10.1073/pnas.1717762115. Epub 2018 Mar 12. PMID: 29531053.

[vandeLoosdrechtScience2018]:*
***Pleistocene North African genomes link Near Eastern and sub-Saharan African human populations.*** *van de Loosdrecht M, Bouzouggar A, Humphrey L, Posth C, Barton N, Aximu-Petri A, Nickel B, Nagel S, Talbi EH, El Hajraoui MA, Amzazi S, Hublin JJ, Pääbo S, Schiffels S, Meyer M, Haak W, Jeong C, Krause J. Science. 2018 May 4;360(6388):548-552. doi: 10.1126/science.aar8380. Epub 2018 Mar 15.

[VanDenBrink2017]:*
***A Late Bronze Age II clay coffin from Tel Shaddudin the Central Jezreel Valley, Israel: context andhistorical implications.*** *Edwin C. M. van den Brink, Ron Beeri, Dan Kirzner, Enno Bron, Anat Cohen-Weinberger, Elisheva Kamaisky, Tamar Gonen, Lilly Gershuny, Yossi Nagar, Daphna Ben-Tor,Naama Sukenik, Orit Shamir, Edward F. Maher & David Reich Levant, 49:2, 105-135, DOI:10.1080/00758914.2017.1368204.

[VeeramahPNAS2018]:*
***Population genomic analysis of elongated skulls reveals extensive female-biased immigration in Early Medieval Bavaria.*** *Veeramah KR, Rott A, Groß M, van Dorp L, López S, Kirsanow K, Sell C, Blöcher J, Wegmann D, Link V, Hofmanová Z, Peters J, Trautmann B, Gairhos A, Haberstroh J, Päffgen B, Hellenthal G, Haas-Gebhard B, Harbeck M, Burger J. Proc Natl Acad Sci U S A. 2018 Mar 27;115(13):3494-3499. doi: 10.1073/pnas.1719880115. Epub 2018 Mar 12. pPMID: 29531040.

[VyasAJPA2017]:*
***Testing support for the northern and southern dispersal routes out of Africa: an analysis of Levantine and southern Arabian populations.*** *Vyas DN, Al-Meeri A, Mulligan CJ. Am J Phys Anthropol. 2017 Dec;164(4):736-749. doi: 10.1002/ajpa.23312. Epub 2017 Sep 15. PMID: 28913852.

[WangNatureCommunications2019]:*
***Ancient human genome-wide data from a 3000-year interval in the Caucasus corresponds with eco-geographic regions.*** *Wang CC, Reinhold S, Kalmykov A, Wissgott A, Brandt G, Jeong C, Cheronet O, Ferry M, Harney E, Keating D, Mallick S, Rohland N, Stewardson K, Kantorovich AR, Maslov VE, Petrenko VG, Erlikh VR, Atabiev BC, Magomedov RG, Kohl PL, Alt KW, Pichler SL, Gerling C, Meller H, Vardanyan B, Yeganyan L, Rezepkin AD, Mariaschk D, Berezina N, Gresky J, Fuchs K, Knipper C, Schiffels S, Balanovska E, Balanovsky O, Mathieson I, Higham T, Berezin YB, Buzhilova A, Trifonov V, Pinhasi R, Belinskij AB, Reich D, Hansen S, Krause J, Haak W. Nat Commun. 2019 Feb 4;10(1):590. doi: 10.1038/s41467-018-08220-8. PMID: 30713341.

[WangBioRxiv2020]:*
***The Genomic Formation of Human Populations in East Asia.*** *Chuan-Chao Wang, Hui-Yuan Yeh, Alexander N Popov, Hu-Qin Zhang, Hirofumi Matsumura, Kendra Sirak, Olivia Cheronet, Alexey Kovalev, Nadin Rohland, Alexander M. Kim, Rebecca Bernardos, Dashtseveg Tumen, Jing Zhao, Yi-Chang Liu, Jiun-Yu Liu, Matthew Mah, Swapan Mallick, Ke Wang, Zhao Zhang, Nicole Adamski, Nasreen Broomandkhoshbacht, Kimberly Callan, Brendan J. Culleton, Laurie Eccles, Ann Marie Lawson, Megan Michel, Jonas Oppenheimer, Kristin Stewardson, Shaoqing Wen, Shi Yan, Fatma Zalzala, Richard Chuang, Ching-Jung Huang, Chung-Ching Shiung, Yuri G. Nikitin, Andrei V. Tabarev, Alexey A. Tishkin, Song Lin, Zhou-Yong Sun, Xiao-Ming Wu, Tie-Lin Yang, Xi Hu, Liang Chen, Hua Du, Jamsranjav Bayarsaikhan, Enkhbayar Mijiddorj, Diimaajav Erdenebaatar, Tumur-Ochir Iderkhangai, Erdene Myagmar, Hideaki Kanzawa-Kiriyama, Msato Nishino, Ken-ichi Shinoda, Olga A. Shubina, Jianxin Guo, Qiongying Deng, Longli Kang, Dawei Li, Dongna Li, Rong Lin, Wangwei Cai, Rukesh Shrestha, Ling-Xiang Wang, Lanhai Wei, Guangmao Xie, Hongbing Yao, Manfei Zhang, Guanglin He, Xiaomin Yang, Rong Hu, Martine Robbeets, Stephan Schiffels, Douglas J. Kennett, Li Jin, Hui Li, Johannes Krause, Ron Pinhasi, David Reich bioRxiv 2020.03.25.004606; doi: https://doi.org/10.1101/2020.03.25.004606.

[WangSciAdv2020]:*
***Ancient genomes reveal complex patterns of population movement, interaction, and replacement in sub-Saharan Africa.*** *Wang K, Goldstein S, Bleasdale M, Clist B, Bostoen K, Bakwa-Lufu P, Buck LT, Crowther A, Dème A, McIntosh RJ, Mercader J, Ogola C, Power RC, Sawchuk E, Robertshaw P, Wilmsen EN, Petraglia M, Ndiema E, Manthi FK, Krause J, Roberts P, Boivin N, Schiffels S. Sci Adv. 2020 Jun 12;6(24):eaaz0183. doi: 10.1126/sciadv.aaz0183. PMID: 32582847; PMCID: PMC7292641.

[YangCurrentBiology2017]:*
***40,000-Year-Old Individual from Asia Provides Insight into Early Population Structure in Eurasia.*** *Yang MA, Gao X, Theunert C, Tong H, Aximu-Petri A, Nickel B, Slatkin M, Meyer M, Pääbo S, Kelso J, Fu Q. Curr Biol. 2017 Oct 23;27(20):3202-3208.e9. doi: 10.1016/j.cub.2017.09.030. Epub 2017 Oct 12. PMID: 29033327.

[YangScience2020]:*
***Ancient DNA indicates human population shifts and admixture in northern and southern China.*** *Yang MA, Fan X, Sun B, Chen C, Lang J, Ko YC, Tsang CH, Chiu H, Wang T, Bao Q, Wu X, Hajdinjak M, Ko AM, Ding M, Cao P, Yang R, Liu F, Nickel B, Dai Q, Feng X, Zhang L, Sun C, Ning C, Zeng W, Zhao Y, Zhang M, Gao X, Cui Y, Reich D, Stoneking M, Fu Q. Science. 2020 Jul 17;369(6501):282-288. doi: 10.1126/science.aba0909. Epub 2020 May 14. PMID: 32409524.

[YuCell2020]:*
***Paleolithic to Bronze Age Siberians Reveal Connections with First Americans and across Eurasia.*** *Yu H, Spyrou MA, Karapetian M, Shnaider S, Radzevičiūtė R, Nägele K, Neumann GU, Penske S, Zech J, Lucas M, LeRoux P, Roberts P, Pavlenok G, Buzhilova A, Posth C, Jeong C, Krause J. Cell. 2020 Jun 11;181(6):1232-1245.e20. doi: 10.1016/j.cell.2020.04.037. Epub 2020 May 20. PMID: 32437661.

[ZallouaScientificReports2018]:*
***Ancient DNA of Phoenician remains indicates discontinuity in the settlement history of Ibiza.*** *Zalloua P, Collins CJ, Gosling A, Biagini SA, Costa B, Kardailsky O, Nigro L, Khalil W, Calafell F, Matisoo-Smith E. Sci Rep. 2018 Dec 4;8(1):17567. doi: 10.1038/s41598-018-35667-y. PMID: 30514893.*
